# Evaluating rotations as a strategy to delay the evolution of insecticide resistance in vectors of human diseases

**DOI:** 10.1101/2022.10.07.511276

**Authors:** Ian Hastings, Sam Jones, Andy South

## Abstract

**Introduction:** Insecticide resistance threatens the control of important human vector-borne diseases such as malaria and dengue. The current de facto strategy is to target the insect vectors using a sequential deployment of insecticides i.e. use one insecticide (usually the cheapest available) until resistance has made it ineffective and then replace it with the next insecticide in the repertoire. Rotations of insecticides are often advocated as a potentially superior method of using the insecticide repertoire to delay the evolution of resistance. Testing this hypothesis in the field is logistically demanding and an *in silico* approach offers a much faster, flexible and transparent method of evaluating rotations.

**Methods:** We develop an *in silico* approach to evaluate rotations using sequential deployment as the baseline. We explored a wide range of deployment scenarios, underlying genetics of resistance, and incorporated costs of resistance and gene flow to/from untreated refugia.

**Results:** We found that, under most circumstances, resistance to all the insecticides in the repertoire were reached at very similar times for rotations and sequences. Any advantages of one strategy over the other tended to be small (typically <10%) and unpredictable.

**Conclusions:** Operational factors, such as cost or supply-chain security, may therefore largely determine the optimal choice of insecticide deployment strategies.

## 1. Introduction

The evolution of insecticide resistance (IR) is a major problem in agricultural pests and in insects responsible for transmitting a range of human infections, most notably malaria and dengue. This has stimulated a considerable body of field and modelling work on the evolution of resistance under different insecticide or drug strategies [1]. Insects cause large economic costs in agriculture and there has been substantial research on how best to deploy insecticides to minimise such costs while reducing selection driving the evolution of IR. Conversely, there is only a relatively small evidence base to support operational decisions about the use of insecticides in public health (e.g. [2–4]) and most discussion of IRM in public health has been imported directly from agricultural practices [5]. There are key differences between insecticide deployments used in agriculture and to control vectors of diseases. These are described in Table 1 and discussed later but, in summary, agriculture tends to frequently apply a wide range of insecticides, often against multiple pest species and lifecycle stages. In contrast, public health applications tend to use extremely infrequent applications of long-lasting insecticides directed at a few key species responsible for local disease transmission.

**Table 1.** Key factors driving differences in the use of insecticides in agriculture and as public health tools to control disease transmission.

| Factors | Agriculture in high-income settings | Public Health in low income countries |
| --- | --- | --- |
| Market forces | High; competition for market share results in large numbers of available products and formulations. | Low; absence of profits means small number of available products, often repurposed from agriculture. |
| Local decision making | Farmers often subjected to high volumes of advertising, advice from growing associations and sometimes central government, usually advocating IRM strategies based on widespread availability of multiple products. | Local implementers often have no choice with product choice dictated by central government or by donor organisations. Financial restrictions dictate this usually results in widespread coverage of a single insecticide (i.e. the cheapest available product). |
| Target species | Numerous | Often focussed on one or two key vector species |
| Lifecycle stages | Often need to target multiple stages e.g. adults and caterpillars | Usual focussed on adult female mosquitoes. Some larval source control in Aedes where urban breeding sites easily identified. Anopheles breeding sites often too numerous to identify and target |
| Ecological impact | Public opinion and regulatory bodies often require that pest control must minimise ecological impact on non-target species. Insecticides have different potencies on different species hence rotations may minimise ecological impact | Local concerns about ecological impact of disease control is generally a negligible consideration compared to the need for disease control. |
| Persistence of insecticides | Generally very short (weeks) as insecticides washed off crops by rain and/or degraded by sunlight. Often considered desirable as it minimises insecticide residues when food crops reach market | Designed to be long (month/years). |
| Frequency of insecticide use | Weeks/months during growing season. | Months/years and often targeted to disease transmission season. |

This paper uses an in silico approach to investigate the effectiveness of insecticide rotations which are widely advocated as one approach for IRM in public health [5]. We compare them to the baseline “sequential” approach which is the current default method. The ‘rules’ for deploying these policies, as considered here, are as follows:

- Sequential deployment is the practice of using an insecticide until resistance to it reaches a certain threshold (usually based on measuring the level of IR in bottle or cone bioassays) which is deemed to indicate a lack of effectiveness, then switching to a different insecticide. We assume that once switching has “used up” all the insecticides in the repertoire, the first one can be re-deployed if resistance has declined since it was last used; this is what will occur in practice. Cost is usually the major driver of this strategy as most vector control occurs in low-resource regions and public health agencies start with the cheapest (pyrethroids), then move to next cheapest. Funding a change in insecticides is often challenging and ineffective insecticides may continue to be deployed simply because the alternative insecticides are locally unaffordable.
- Rotation of insecticides is a policy where the currently-deployed insecticide is changed at set time intervals irrespective of resistance levels. Different rotation time intervals can be considered, e.g. indoor residual spraying for malaria vector control may be repeated every year (around 10 Anophelene mosquito generations) and insecticide treated nets may be replaced every two years (around 20 generations). Multiples of these intervals could also be used as the rotation interval. Rotations used in public health therefore invariably cover multiple generations, in contrast with the situation in agriculture where a rotation is sometimes more tightly defined as having an interval of a single generation [6] and multiple spraying of crops with different insecticides may occur over a single growing season (Table 1). Once funding is in place for rotations, it avoids the financial hurdle of having to periodically find increased resources for the switches to more expensive insecticides inherent in sequential policy.

Examples of IRM discussed specifically in relation to public health can be found in Curtis [7] who modelled rotations of insecticide as a IRM and Rowland [8] who discussed costs of resistance and selection coefficients for IR and used these to construct a rotations model tested against field data. More recently, Dusfour and colleagues [2] discussed Aedes (the vector for dengue, Zika and yellow fever) and reviewed the (lack of) evidence supporting different IRM strategies, concluding that “Until now, it has been near impossible to anticipate…….. how a given IRM strategy will slow or reverse the evolution of resistance in targeted populations”. We agree with this remark and this manuscript addresses this knowledge gap.

## 2. Methods

A population genetic model was developed to simulate the evolution of insecticide resistance in response to insecticides deployed as rotations and sequences. The algebra is described in the supplementary information. The model uses a similar approach to that previously used to compare insecticide mixtures and sequences (Levick et al., 2017; South & Hastings, 2018). The implementation is simpler here because only one insecticide is deployed at a time so linkage disequilibrium can be ignored. The simpler implementation allows the model to be run for an unlimited number of insecticides. The model runs on a timestep of mosquito generations and resistance allele frequency is recorded at each generation.

### 2.1 Genetics of resistance and the impact of the mutations(s) on insect fitness and mortality

The fitness of different genotypes is determined by parameters representing the impact of insecticide on each of the three genotypes. We also allow a fitness cost to be associated with the resistance allele. Full details are in the SI but here we provide the general principles. Notably, (i) fully sensitive insects (genotype SS) may survive contact with the insecticide (ie. their mortality need not be 100% as mosquitoes often encounter low concentrations of insecticide [9]) and (ii) fully resistant insects (genotype RR) may still be killed by contact with the insecticide (i.e. mortality need not be 0% as high insecticide concentrations often kill “resistant” genotypes[9]).

If the mosquito encounters an insecticide (denoted by subscript +)

- The SS genotype has its fitness 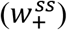 determined by the “effectiveness” of the insecticide i.e. its ability to kill SS genotypes. The fitness is 1-effectiveness
- The RR genotypes has its fitness 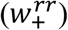 determined by “resistance restoration” which determines the extent to which the resistance mutation allows the RR genotype to survive insecticide contact. It is defined relative to the SS genotype so a “resistance restoration” value of 0 means it has the same fitness as the SS genotype while a value of 1 means it always survives contact with the insecticide.
- The SR genotypes has a fitness 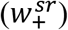 intermediate between that of SS and RR as determined by the “dominance” of the resistant mutation.

If the mosquito does not encounter an insecticide (denoted by subscript -)

- The SS genotype does not incur a cost i.e. has fitness 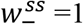
- The RR genotype bears the full “cost”, c, so has fitness 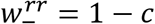
- The SR genotype has the cost that lies between zero and the full costs so it fitness 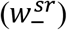 depends on the “dominance” of the cost

Each insecticide is assumed to have a corresponding resistance allele with no cross-resistance to the others.

The parameters described in Table 2 determine the fitness of insect genotypes as summarised above and as shown graphically on Figure 1; they are discussed in more detail in the SI and elsewhere [10, 11]. These “genetic” fitnesses are then updated to construct “overall” fitnesses that depend on how often the mosquitoes meet insecticides, noting that the overall fitnesses of males and females may differ as females may encounter insecticides more frequently while foraging for a human blood meal. Differences in the overall fitness of the genotypes drives selection and increases the resistance allele frequencies over the generations. We set initial frequencies (i.e. at generation zero) to start the selection process (Table 2).

**Table 2.** Parameter definitions, their range of values explored in the simulations, and their values used in the plots.

| **Input** | **Description** | **Range\*** | **Fig 4** | **Figs 5 &6** | **Fig 7** | **Fig 8<sup>+</sup>** |
| --- | --- | --- | --- | --- | --- | --- |
| **Insecticide number** | Number of independent insecticides available | 2→5 | 3 | 3 | 3 | 3 |
| **Coverage** | Proportion of the insect population in the treated intervention area (only applies to scenario with refugia) | 0.1→0.9 | 1 | 1 | 0.3 | 1 |
| **Dispersal** | Proportion of population exchanged between intervention and refugia areas per generation, 0=none, 1=random mixing | 0.1→0.9 | 0 | 0 | 0.1 | 0 |
| **Exposure** | Proportion of females exposed to insecticide in the intervention sites | 0.4→0.9 | 0.75 | 0.75 | 0.75 | 0.65 |
| **Male exposure** | Proportion of males exposed to insecticide in the intervention sites as a proportion of female exposure | 0→1 | 0.34 | 0.34 | 0.34 | 0.087 |
| **Effectiveness** | Proportion of susceptible (SS) insects killed by exposure to insecticide | 0.7<sup>s</sup>→1 | 0.98 | 0.98 | 0.98 | 1, 0.96, 0.78 |
| **Resistance restoration** | Ability of fully resistant (RR) genotype to restore fitness of insects exposed to insecticide | 0.1→0.9 | 0.12 | 0.12 | 0.12 | 0.36, 0.11, 0.86 |
| **Dominance of resistance** | sets fitness of heterozygous (SR) insects between that of SS & RR in presence of insecticide | 0→1 | 0.96 | 0.96 | 0.96 | 0.63, 0.95, 0.42 |
| **Cost of resistance** | fitness of resistant (RR) insects in absence of insecticide | 0→0.1 | 0 | 0.02 | 0 | 0.099,<br>0.088,<br>0.062 |
| **Dominance of cost** | sets fitness of heterozygous (SR) insects between that of SS & RR in absence of insecticide | 0→1 | n/a | 0.32 | n/a | 0.6,<br>0.49,<br>0.14 |
| **Frequency** | Starting frequency of resistance alleles within the population | 0.001→0.1 | 0.001 | 0.001 | 0.001 | 0.0059 |

\*Ranges are uniform except for starting frequency which is log-uniform
\$ 0.9 when using mortality as the threshold for switching
<sup>+</sup> where 3 values are given they correspond to insecticide 1,2 and 3 respectively

**Figure 1.**
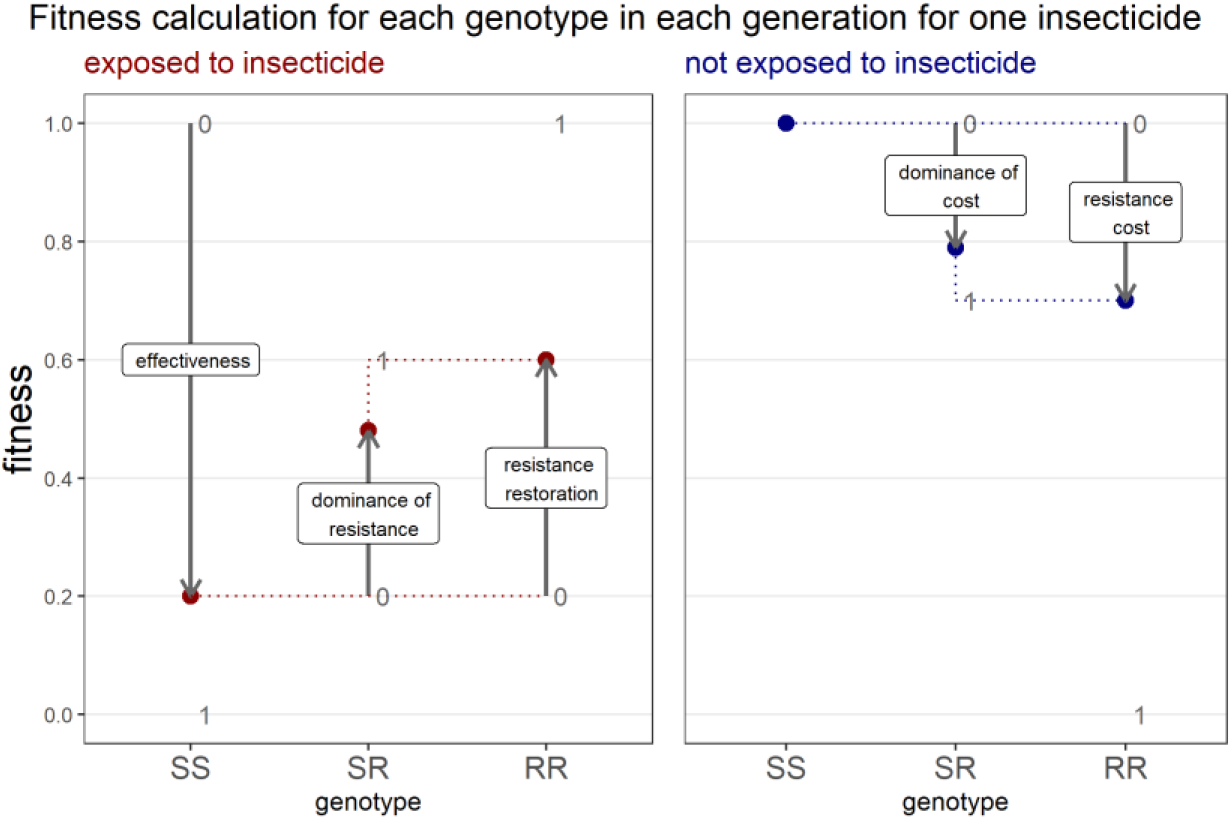
The effect of model inputs on the fitness of genotypes ending resistance to a single insecticide. Fitness is shown on the y-axis and the different genotypes (SS, SR, RR) on the x axis. Insects exposed to the insecticide will have fitness determined by the left panel as follows: insecticide effectiveness sets the fitness for SS, resistance restoration ‘restores’ a portion of the fitness for RR and dominance of resistance determines how the fitness for SR lies between that of SS and RR. Insects not exposed to the insecticide will have fitness determined by the right panel as follows: fitness of SS is set to 1 by definition, resistance cost determines the fitness of RR and again dominance of cost determines how the fitness for SR sits between that of SS and RR. In this example effectiveness=0.8, resistance restoration=0.5 which ‘restores’ half of the fitness lost due to the insecticide, dominance of resistance=0.7 which sets the fitness of the SR closer to RR than SS. Resistance cost=0.3 which reduces fitness in the absence of the insecticide from 1 to 0.7, and dominance of cost=0.8 which sets fitness of SR close to RR. Reprinted from [11].

The frequency of resistance determines how much impact the insecticide(s) have on the mosquito population so we calculate two mortality measures: ‘field’ and ‘bioassay’ mortalities. Field mortality quantifies the impact of the intervention in terms of killing mosquitoes and will be used later to evaluate relative performances of sequences and rotations in their ability to kill during their field application. Field mortality, μ_f_, depends on the ‘exposure’ (i.e. percentage of female mosquitoes exposed to the insecticide in the intervention site(s); Table 2)

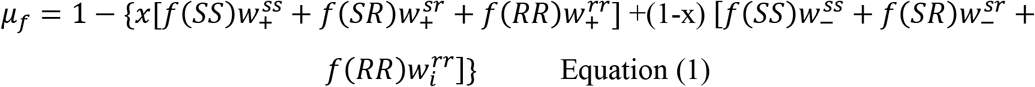

Where x is female exposure to the insecticide and f() is the frequency of the genotypes. The term in the curly brackets is the overall ‘fitness’ i.e. ability to survive in the presence of insecticide deployment, so 1 minus this term is the mortality. Note that this is mortality of females in the interventions site which is appropriate as only female mosquitoes are vectors of human diseases.

Bioassay mortality is a measure used for surveillance, monitoring and, in practice, is often the metric used to determine when a an insecticide is “failing” and needs to be replaced. Bioassay mortality is calculated as in Equation (1) noting that exposure to insecticides in assays is obviously 100% so that x=1 and the equations reduces to

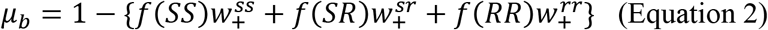

Noting that this makes the implicit assumption that mortality in a bio-assay reflects mortality occurring in the field (and hence driving resistance); for further discussion on this see, for example, [12, 13].

### 2.2 Insecticides and their deployment strategies

The model allows a single insecticide to be deployed at any one time. There are a defined number of available insecticides in a “repertoire” which is one of the inputs of the calculations (Table 2). Insecticide properties were varied in each run according to the input ranges in Table 2. In one set of runs all insecticides in the repertoire were assumed to have identical properties. In another set of runs insecticides were allowed to have different properties randomly selected from the inputs described in Table 2 (i.e., effectiveness, resistance restoration, dominance of resistance, cost of resistance and dominance of cost). The runs with identical insecticides are obviously unrealistic but allowed us to calculate the effect of insecticide properties on model outputs (where properties differ the outputs are largely driven, and obscured, by the insecticide least susceptible to resistance evolution). The runs with different insecticides provide a closer representation to the behaviour we expect in the field where different insecticides are likely to have different properties.

The proportion of insects exposed to the insecticide in the intervention site is defined by the “exposure” parameter. There is differential exposure of male and females in mosquito vectors as only female seek human blood meals and are more likely to encounter insecticides used for protective measures such as bednets and wall-spraying. The “male exposure” parameter allows exposure of males to be less than that of females.

We allow for the presence of untreated refugia whose size is specified by the ‘coverage’ input defined as the proportion of the insect populations in the intervention site(s). Movement between intervention and untreated “refugia” sites is quantified by the ‘dispersal’ parameter.

Insecticides can only be changed on an annual basis, which is approximately ten generations of Anopheles spp that transmit malaria in the most badly affected regions of Sub-Saharan Africa. This annual basis occurs because public-health insecticides are designed to be highly persistent, and this is the most common situation for Insecticide Residual Spraying (IRS) where houses are most often sprayed once per year. The two key strategies are defined as follows:

- *Rotations*. The insecticide is automatically changed at the end of each year and replaced by the next insecticide in a pre-defined sequence. If resistance to the next pre-planned insecticide exceeds a resistance threshold (see later), then it is not used, and the rotation uses the next insecticide in the pre-planned sequence. If the currently deployed insecticide is the only one remaining below threshold, we assume it be used again in that year (which we argue is what would occur in practice).
- *Sequences*. An insecticide change is considered each year but only implemented if resistance to the current insecticide exceeds a resistance threshold. The replacement insecticide is chosen from any remaining insecticides that are below their own resistance thresholds.

The decision on whether to continue with the current insecticide (sequences) or whether to use the next insecticide in the rotation is made immediately before the switch is (potentially) made. This is done of the basis of resistance status at the end of the previous generation. In both strategies it is permissible to switch back to an insecticide that had previously been deployed and reached its resistance threshold, providing resistance to the insecticide had declined to a level below this threshold. This decline can occur during periods of non-deployment because natural selection acts against the resistance allele (due to their carrying a fitness cost) and/or because gene flow in/out of a refugia had reduced the resistance allele frequency in the intervention site.

### 2.3. Evaluating the difference between rotations and sequences

The model runs on a timestep of a generation and the resistance allele frequency is recorded at each generation. The operational lifespan of the strategy in each run is reached when resistance to the current insecticide and all potential replacements exceed the resistance threshold. However, if this does not occur after five hundred generations (approximately 50 years) we terminate the simulations on the basis that 50 years more than exceeds the timescale of operational planning.

Two resistance threshold criteria were investigated as criteria to trigger a change of insecticides.

- Resistance allele frequency. This was a common criterion used in previous theoretical modelling when, typically, a threshold of >50% was used (e.g. [10, 11, 14]). We investigate this threshold for consistency with previous work but note its drawbacks (i) the gene encoding resistance may not be identified in real deployment and hence would not be available for policy makers as a basis of decision making (ii) several genes may actually act to encode resistance (iii) it does not have a direct operational interpretation because the same allele frequency may have different impacts on mortality depending on their level of dominance
- Mosquito mortality in bioassay. This is obtained from resistance frequency as described in Equation 2. We use a mortality of <90% as the switching threshold, consistent with the WHO definition for “confirmed” resistance following a diagnostic test [15]. Note that, when using the mortality threshold, we have to avoid runs that started with all insecticides already above this mortality threshold (i.e. insecticide effectiveness against SS genotypes must be >0.9; Table 2). Mortality appears to be rarely used as a criterion in modelling work (but see Gould [16]) but we argue it is more representative of the switching criterion likely to be used in practice

Each model run was analysed to produce two outputs used to compare rotations and sequences. Firstly, the number of generations taken until all insecticides are above the resistance threshold; this is the “operational lifespan” of the insecticide armoury. Secondly the mean field mortality imposed on the insects during the operational lifespan (obtained from equation 1). The ‘best’ runs are those with the longest operational lifespans and/or highest mean mortalities. We quantify the relative performance of sequential and rotations policy (see later discussion of Figures 2 and 3) using the metric, z. Comparing strategies based on operational lifespan requires

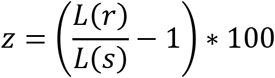

where L(r) and L(s) are the operational lifespans of rotations and sequential use respectively. Alternately, for a comparison based on mean mortality during the operational lifespan

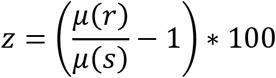

Where μ(r) and μ(s) are the mean mortalities in rotations and sequences respectively.

**Figure 2.**
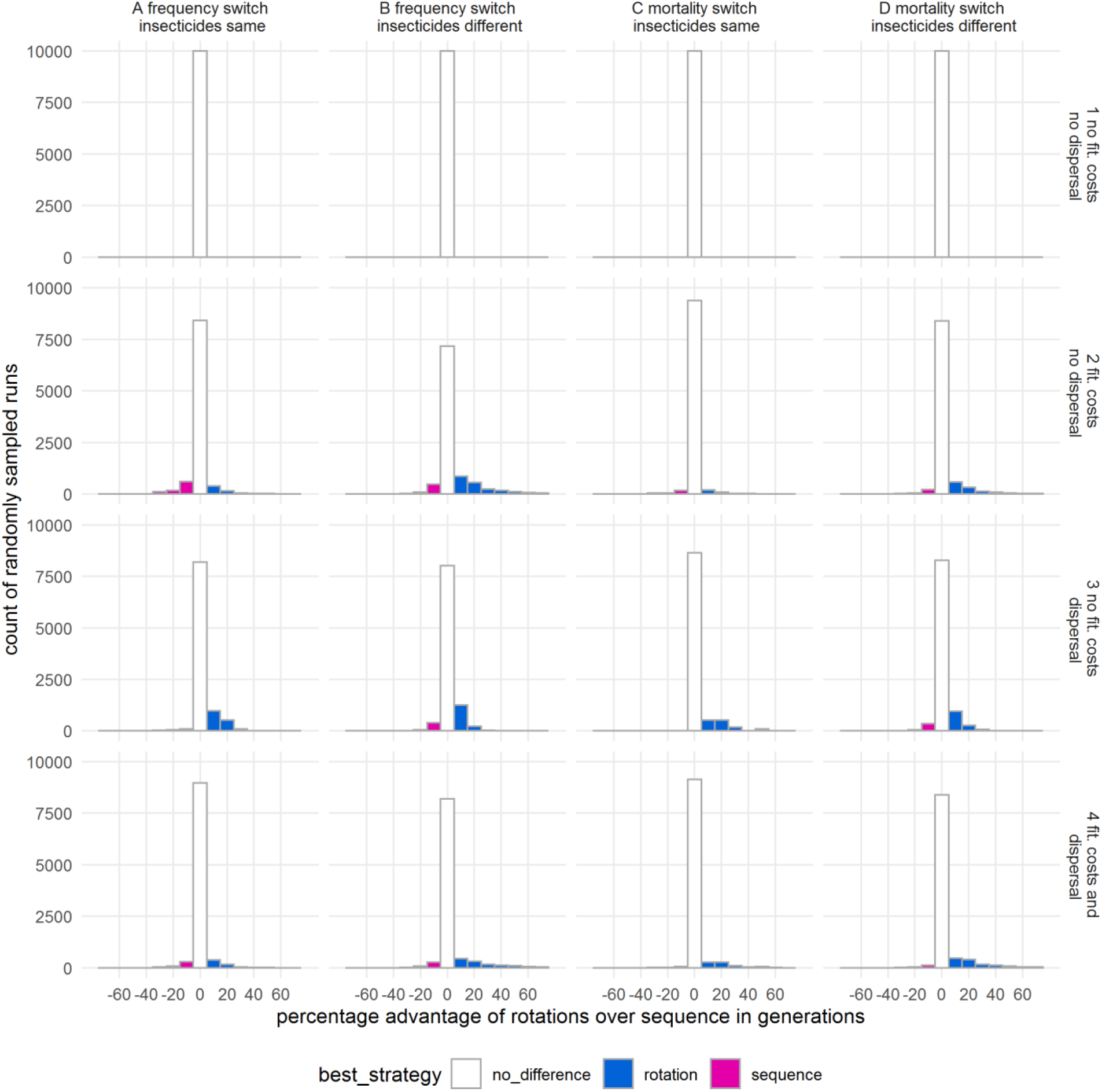
Evaluation of rotations and sequential deployments based on the operational lifespan of a policy. This was defined in two ways:

- If the switching criterion was frequency>50% the operational lifespan was defined as the time until resistance exceeds 50% for all available insecticides.
- If the switching criterion was assay mortality <90% the operational lifespan was defined as the time until mortality was <90% for all available insecticides. Each panel showing the results of 10,000 runs sampling the parameter space specified in Table 2. Each run calculated the operational lifespan of sequences and rotations using the same set of parameters. Column A: each insecticide had identical properties; switching criterion was resistance allele frequency exceeding 50% Column B: insecticides had different properties; switching criterion was resistance allele frequency exceeding 50% Column C: each insecticide had identical properties; switching criterion was mortality <90% Column D: insecticides had different properties; switching criterion was mortality <90% Row 1: no costs of resistance or dispersal into/out of an untreated refugia. Row 2: costs are present; no dispersal to/from refugia Row 3: costs absent; untreated refugia present Row 4: costs and refugia both present A “best strategy” is identified and colour-coded if there is more than a 10% difference, otherwise it is classified as ‘no difference’.

**Figure 3.**
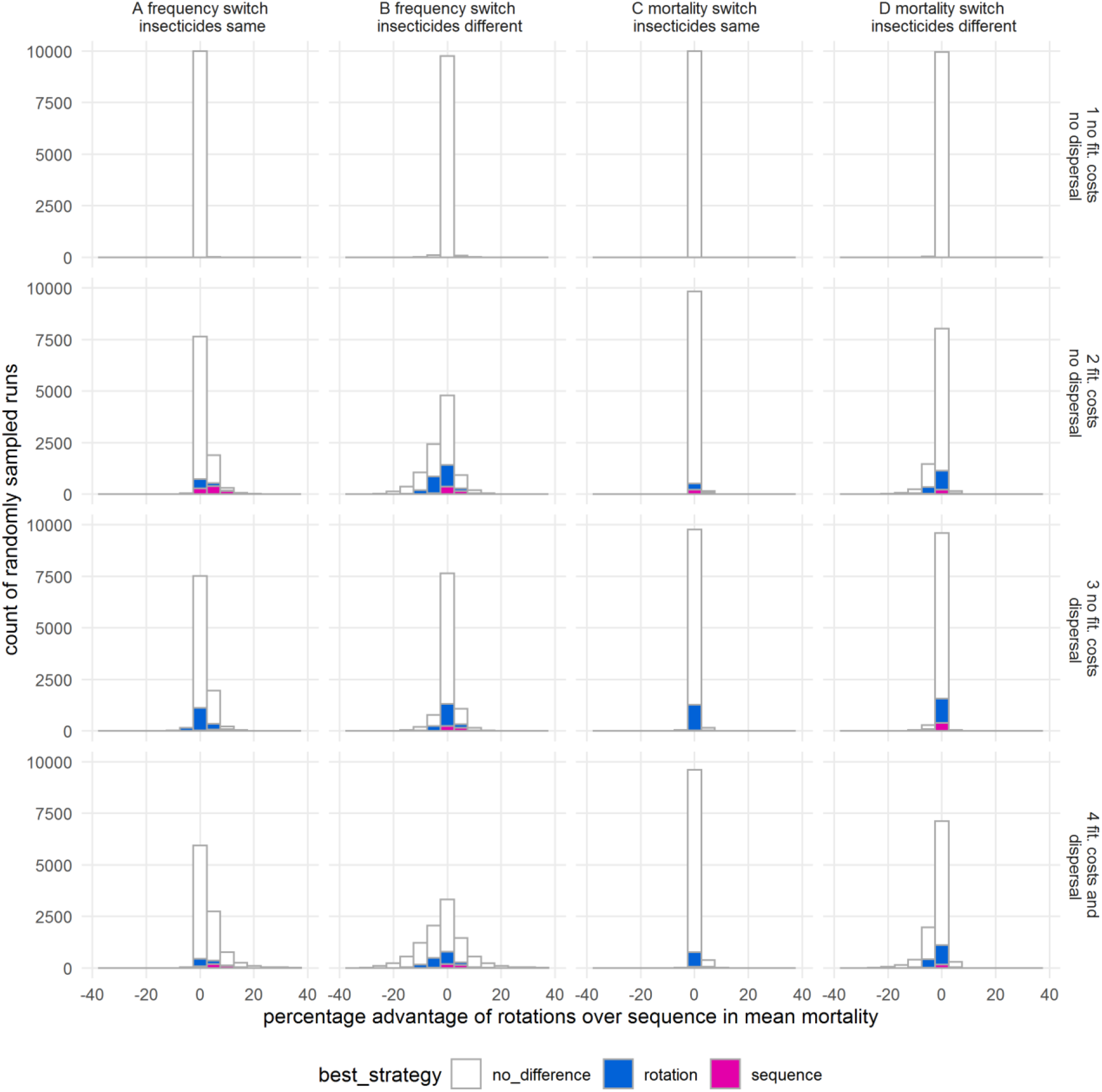
Evaluation of strategies based on the mortality imposed on the insect populations. The figure structure is as for Figure 2 and exactly the same simulations were reanalysed. The difference here is that the strategies are evaluated on the percentage difference in mean mortality achieved over the course of their operational lifespan. The colour coding identifies ‘best-strategies’ identified by the alternative metric, operational lifespan (as shown on Figure 2).

These scales are centred on zero so that positive values of z indicate rotations perform better, negative values of z indicate sequential use performed better and the magnitude of z shows the percentage (dis)advantage. For example, the value z=−10 shows that sequential policy lasted 10% longer than the rotations in that run.

Resistance fitness costs and dispersal between treated and untreated areas have important implications for the evolution of insecticide resistance but their importance in the field is little known and is debated (e.g. [17–20]). We investigate their likely impact by comparing whether the relative performances of sequences and rotations change according to four factors that we suspect may be important and/or have been identified as important in the literature:

- Costs of resistance (present/absent)
- Untreated refugia (present/absent)
- Resistance switching threshold (allele frequency >50% / assay mortality)

Plus the assumption made for analytic clarity:

- Insecticide properties (identical/different)

This gives 16 different scenarios in total and 10,000 runs were run for each scenario using randomly selecting inputs according to the ranges specified in Table 2. We then compared whether rotations were better than sequential use based on the two criteria i.e. duration of the operational lifespans and mean field mortalities during those lifespans.

Sensitivity analyses were conducted using partial rank correlation coefficients (PRCC) and classification trees, and excluded runs where both rotations and sequences reached 500 generations. The reasoning for this exclusion is that including all data would force the analyses to do two things i.e. identify criteria that determine when at least one policy fails before 500 generations, then identify the conditions under which rotations may be favoured or disfavoured; the logic was therefore that omitting runs where both policies lasted >500 generation will allow a more focussed analysis of the conditions under which rotations are favoured.

PRCC analysis was then conducted on those two outcome measures i.e. operational lifespan and mean mortality.

The following parameters were included for PRCC analysis and to construct classification trees in all scenarios:

*Number of insecticides
*Effectiveness
*Resistance_restoration
*Dominance of resistance
Exposure
Male exposure
Starting frequency

Where costs are present, the following parameters also enter the PRCC analysis

*Cost of resistance
*Dominance of cost

Where refugia are present, the following parameters also enter the PRCC analysis

Coverage
Dispersal

Where insecticides differ in their properties, the parameters marked with an asterix are different between insecticides. In this case their mean values are used as input for PRCC an classification trees.

All simulations were run in R version 3.6.3 PRCC analysis was performed using package ppcor (version 1.1) and classification trees using package rpart (version 4.1 - 15).

## 3. Results

Figures and 2 and 3 shows the key results i.e. the relative performance of rotations vs sequences in terms of operational lifespan of the insecticide armoury (Figure 2) or mean mortality imposed on the mosquito population during this lifespan (Figure 3). The clear conclusion is that neither policy is ever advantageous when costs and refugia are absent (top row of Figures 2 and 3) while one policy may be favoured in a small sub-set of simulations when costs and/or refugia are present (lower three rows of Figures 2 and 3). Advantages, if present, are generally less than 10% when evaluated on operational lifespans (Figure 2), and less than around 5% when evaluated by mean mortality (Figure 3).

Figure 4 shows why neither policy is favoured in the absence of costs and refugia. The absence of cost and refugia means that resistance frequencies do not decline when an insecticide is not in use. Consequently, resistance spreads in several large ‘steps’ under sequential use and in more numerous, but smaller, steps in rotations. Importantly, both policies end (because resistance exceeds 50% to all the insecticides in this example) at the same time.

**Figure 4.**
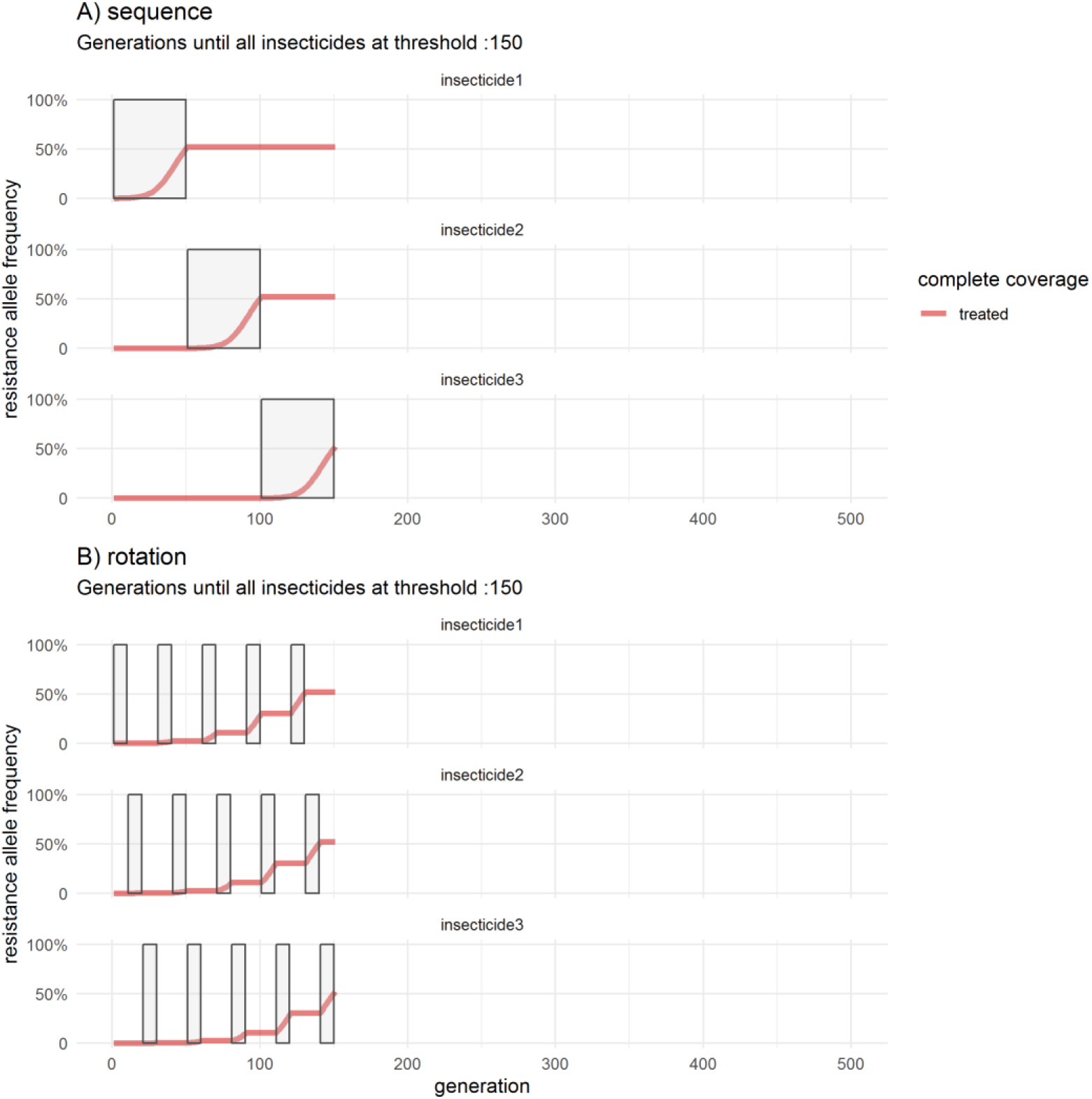
An example of how insecticide resistance evolves under a rotation or a sequence when there are no resistance fitness costs or dispersal from untreated refugia. Three insecticides are available, all of which have identical properties, and are changed when the resistance threshold of 50% is reached. The resistance frequency for each insecticide is shown as red lines in the sub-plots of each panel, and the shaded boxes within each sub-plot indicate when that insecticide was in use. Panel (A) shows the dynamics of resistance spread under sequential use. Panel (B) show the dynamics of resistance to the same three insecticides that occur under a rotation with a regular interval of 10 generations which is approximately a year for Anopheline mosquitoes. Calibration is given in Table 2.

The situation becomes more complicated when costs and/or refugia are present because resistance declines in periods when the corresponding insecticide is not in use. These declines arises from natural selection acting against the fitness cost, or by emigration of resistance allele into the refugia populations.

Figures 5 and 6 show examples where costs are present but refugia absent. The parameters are identical between the plots but the threshold for switching insecticides in sequences is based on resistance allele frequency >50% in Figure 5 and on assay mortality <90% in Figure 6. Resistance allele frequency for each insecticide increases to reach the threshold in one step under sequential use but, in contrast to the previous Figure 4, resistance frequencies decline when the insecticide is not in use due to fitness costs allowing it to be reused later. For the rotation the resistance frequency for each insecticide increases in shorter steps while it is in use and declines while not in use because of fitness costs. In this example, sequential use is slightly favoured over rotations (but note that this is illustrative and does not always occur; see, row 2 of Figures 2 and 3).

**Figure 5.**
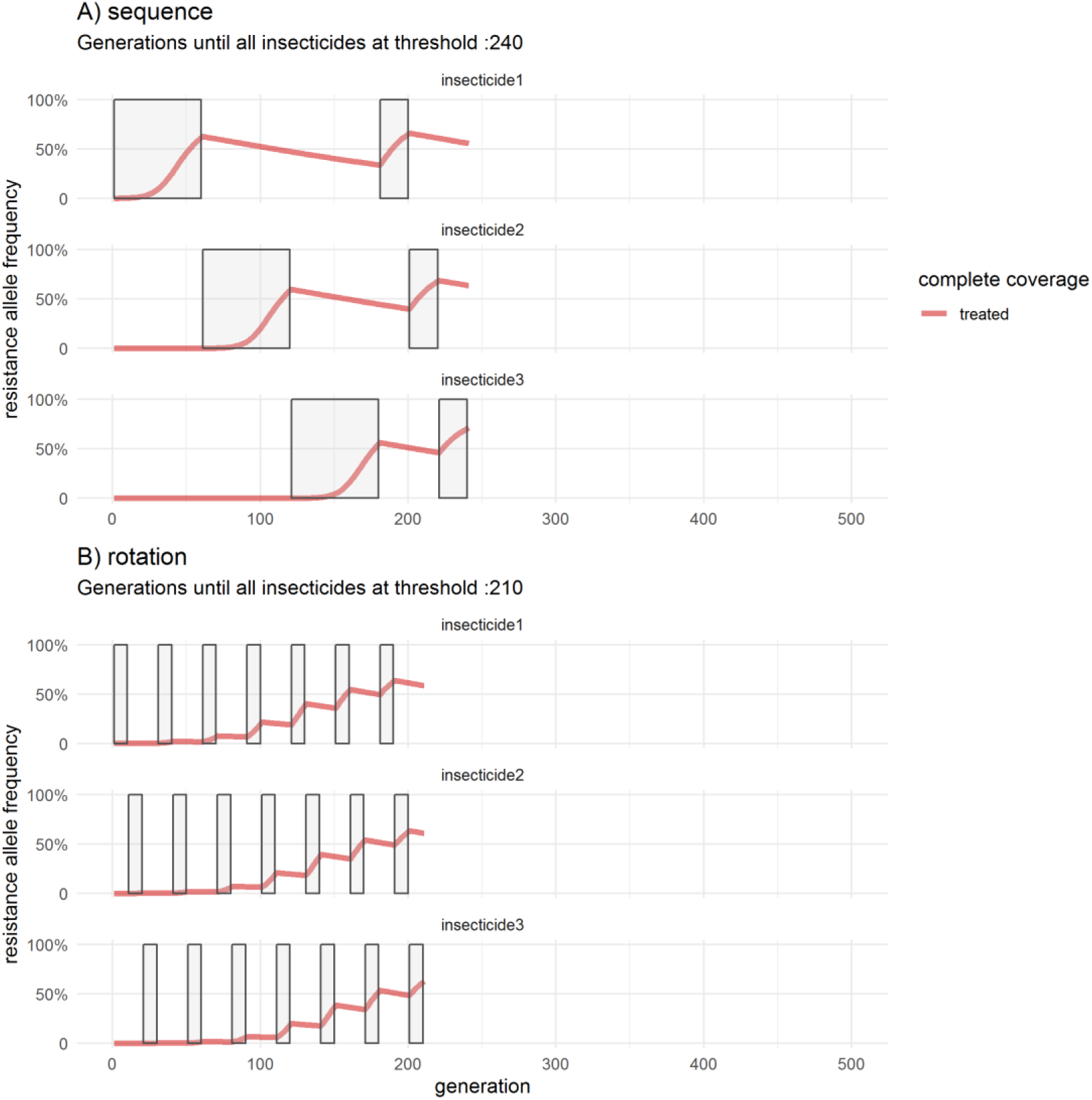
An example of how insecticide resistance evolves under a policy of rotation or sequential use when resistance fitness costs are present but there is no dispersal into/from untreated refugia. This plot follows the same structure as Figure 4 and uses resistance allele frequency of 50% as a switching criterion; note that an insecticide switch can only occur every 10 generations (equivalent to annual deployment) so allele frequency often rises above 50% before the insecticide can be replaced. Calibration is given in Table 2.

**Figure 6.**
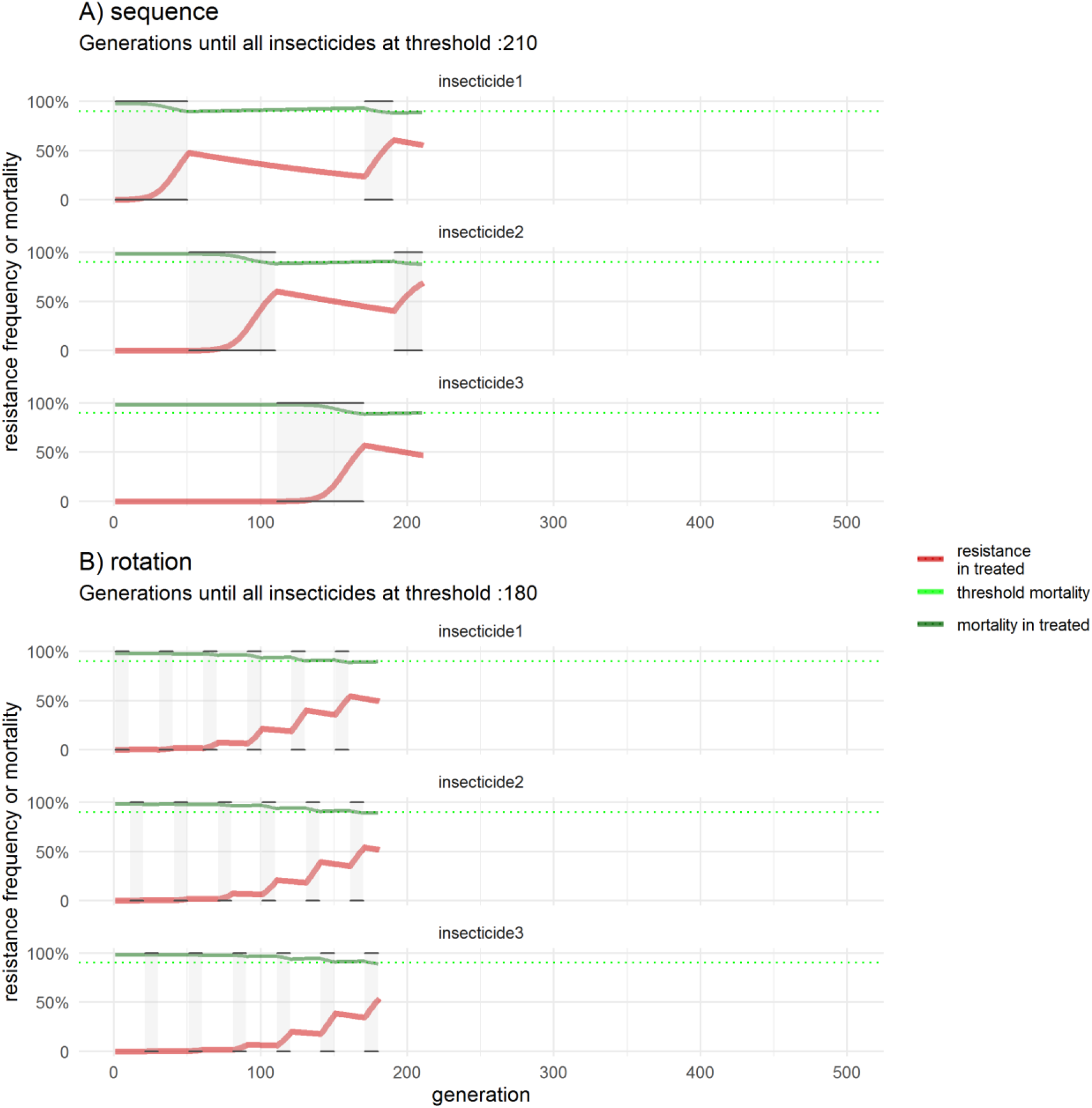
As for Figure 5 but insecticides are replaced in sequential use when their mortality in bioassays falls below 90% (figure 5 use resistance frequency as the switching criterion). As before, resistance frequency in treated areas is given by the red lines, while bioassay mortality, calculated from this frequency, is plotted as green lines (Note that in this calibration, 90% mortality occurs at 45% resistance allele frequency). Calibration is given in Table 2.

Figure 7 show an example where refugia are present but costs are absent. The threshold for switching insecticides in sequences is resistance allele frequency >50%. The presence of refugia and the dispersal of insects between intervention and refugia means that gene flow between the two areas tends to homogenise the allele frequencies in the two areas. This has two noticeable impacts on the dynamics. Firstly, resistance allele frequencies will generally decline in the intervention site when the insecticide is not in use (except in the unlikely case that resistance is actually higher in refugia); this effect can be easily seen in red lines of Figure 7. Secondly, resistance frequencies in the refugia (blue lines in Figure 7) gradually increase due to gene flow from the intervention site(s). Switching insecticides in sequential use occurs sufficiently infrequently that genetic homogeneity is often reached i.e., the resistant allele frequency is identical between intervention and refugia sites (Figure 7A). Insecticide switching occurs so rapidly in rotations that, in this case, homogeneity is rarely reached and resistance frequencies are generally higher in intervention than refugia sites (figure 7B). In this specific example, rotations are slightly better than sequential use (noting, as above, that this is not a universal result; Row 3 of Figures 2 and 3).

**Figure 7.**
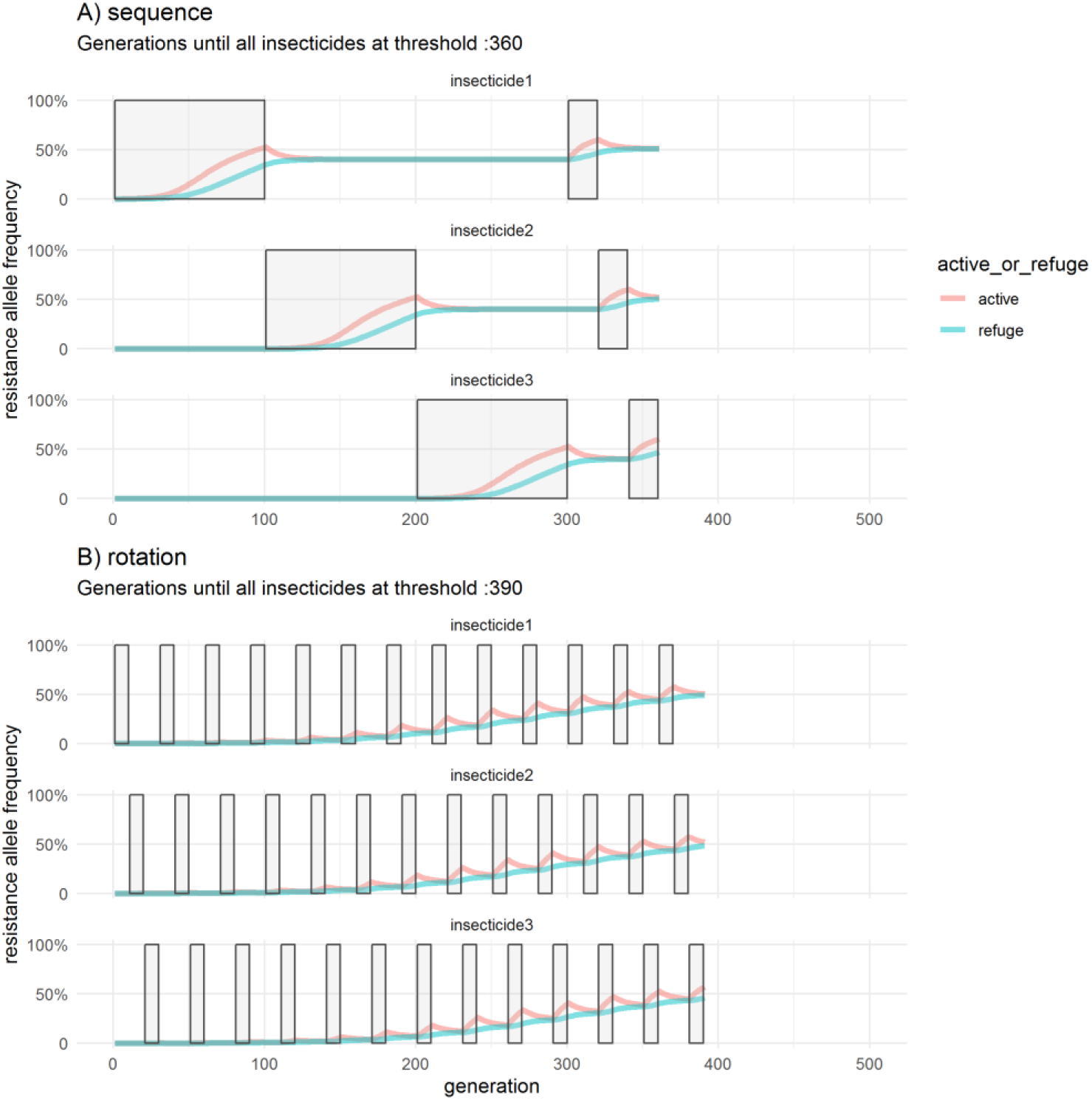
An example of how insecticide resistance evolves under a rotation or a sequence when there is dispersal into/from untreated refugia and resistance fitness costs are absent. Resistance allele frequency of 50% was used as the switching criterion. This plot follows the same structure described for Figure 4 but two resistance allele frequencies are now tracked for each insecticide i.e. frequency in the intervention area (red lines) and frequency in the refugia (green lines). Calibration is given in Table 2.

Figure 8 shows that, as expected, if insecticides differ in their properties then it is the insecticide least susceptible to resistance that dominates the dynamics. In this example both strategies last the full 500 generations (~50 years) of the simulation. Notably, a policy of sequential deployment uses insecticide #2 for longer so that, after 50 years, sequential deployment ends with resistance to all 3 insecticides being below 30% resistance frequency, while a rotations policy ends with two of the insecticides being close to 50% resistance frequency.

**Figure 8.**
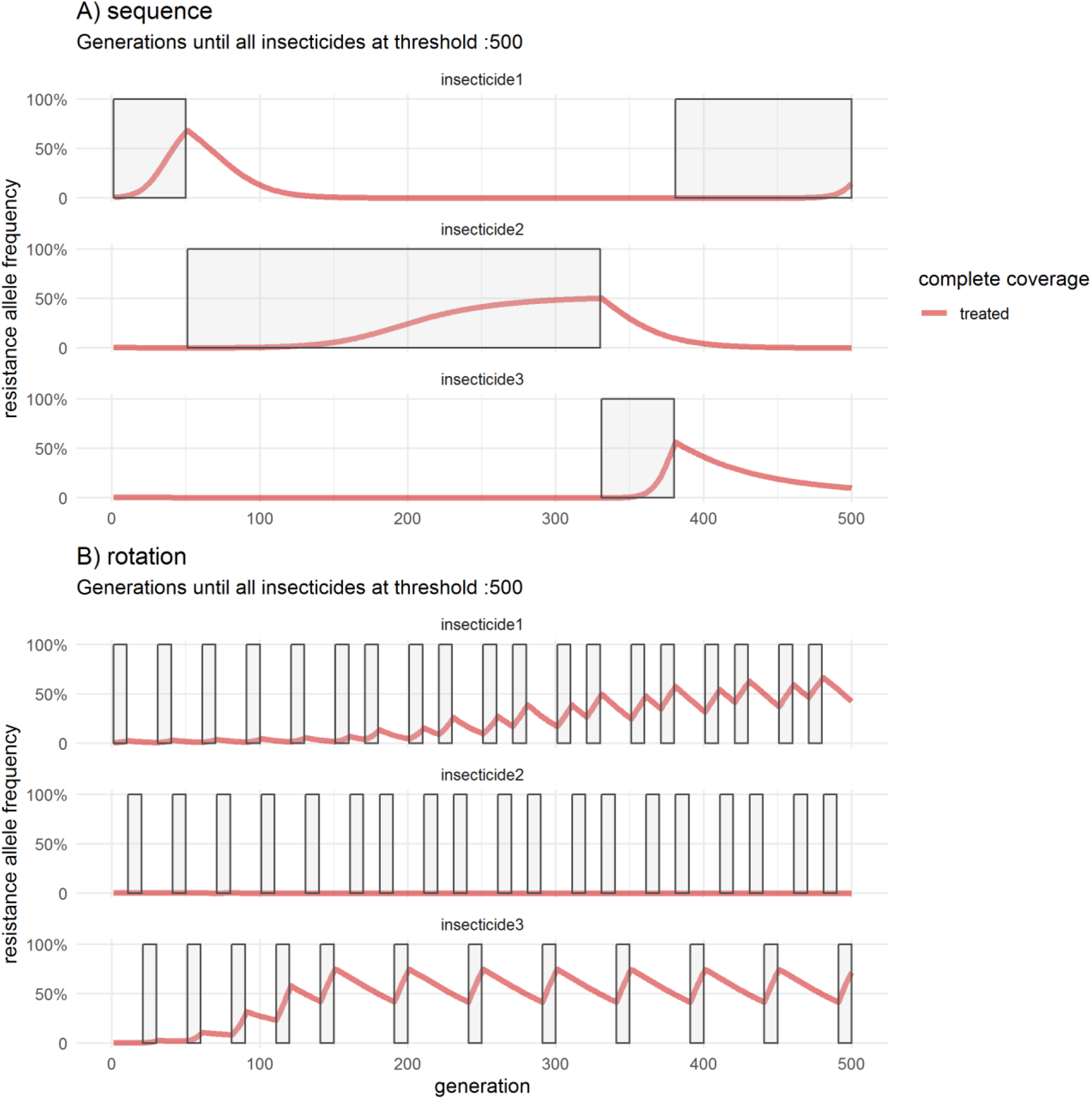
An example of insecticide resistance dynamics when insecticides differ in their properties. This plot is analogous to fig 5, with fitness costs present and refugia absent and uses resistance allele frequency of 50% as a switching criterion. The insecticides have different properties in terms of effectiveness, resistance, costs and dominance (randomly selected from values in table 2). Insecticide2 lasts longer due to its properties, most likely its relatively low value of resistance restoration meaning the RR genotype has low ability to restore fitness of insects exposed to insecticide and hence lower selective advantage. Calibration is given in Table 2.

Figures 9 and 10 shows one of our attempts, based on intuition, to identify specific circumstances under which rotations or sequences are favoured. The figures show the operational lifespan of the insecticides in all the rotation simulation runs, together with coloured bars indicating whether rotations or sequences were favoured by more than 10% in that run. This allows us to check whether the runs in which one strategy was favoured were associated with duration of lifespans. It also shows how lifespans change under different scenarios. For example, comparing rows 1 & 2 shows that the addition of fitness costs increases operational lifespan as expected. Where there are differences between the strategies (rows 3-4) there is a slight suggestion that the runs in which sequences are favoured (pink) are more likely to be those with shorter lifespans and the ones in which rotations are favoured (blue) are more likely to be those with longer lifespans. However, this pattern doesn’t identify any factors that would be operationally useful at this stage.

**Figure 9.**
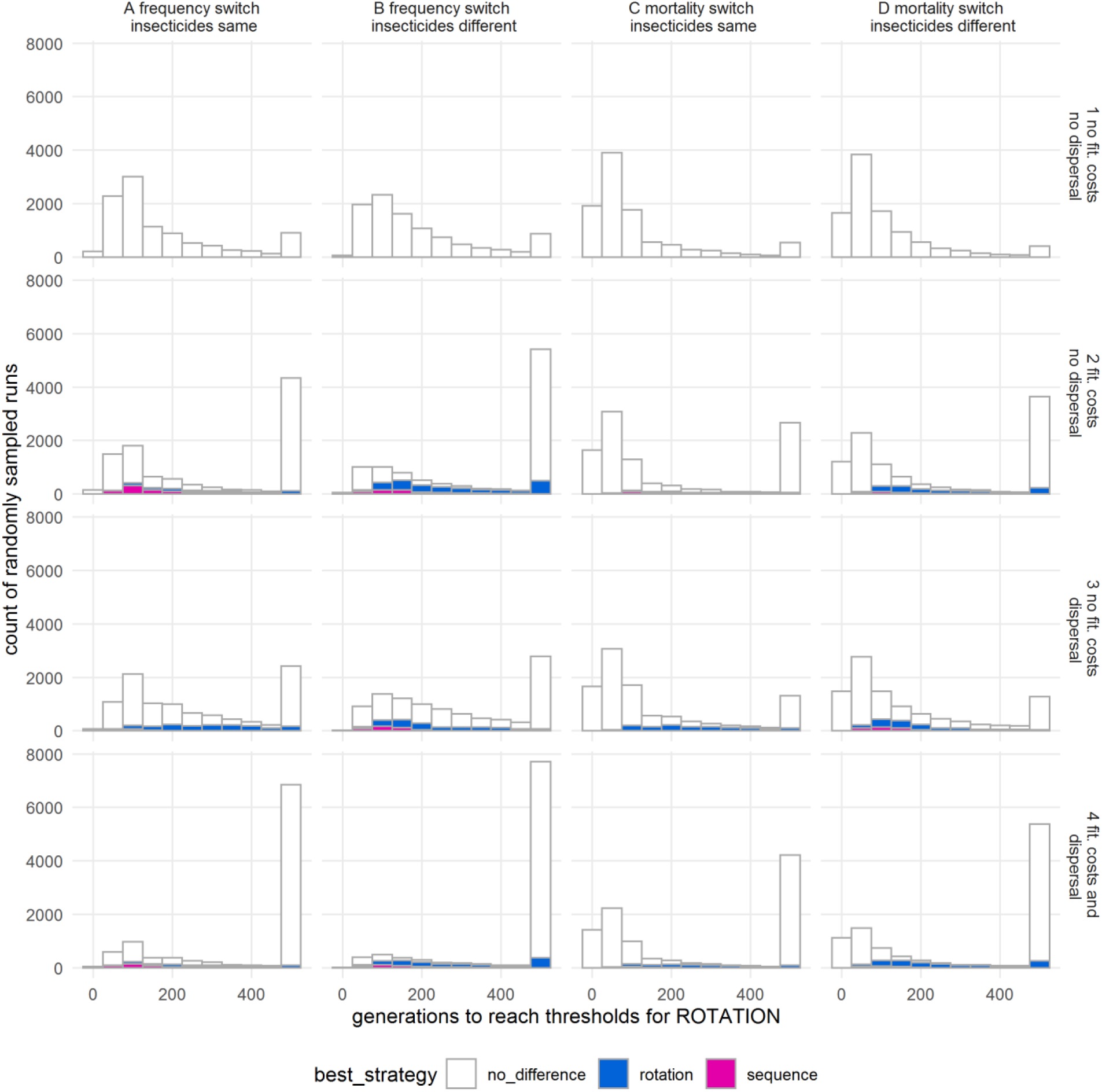
An investigation into whether different polices are favoured depending on their operational lifespan i.e. the number of generations until resistance is present to all insecticides in the repertoire. The same simulations were analysed as in Figures 2 and 3. There is slight tendency for rotations (blue sub-bars) to be favoured when both strategies last a long time with, conversely, sequential use (red sub-bars) being favoured when insecticide resistance evolves rapidly. [“Best” strategies are defined as >10% difference between rotation and sequence. These X axis are operational lifespans under rotations, the analogous plot showing lifespans under sequential deployment is shown in Figure 10]

**Figure 10.**
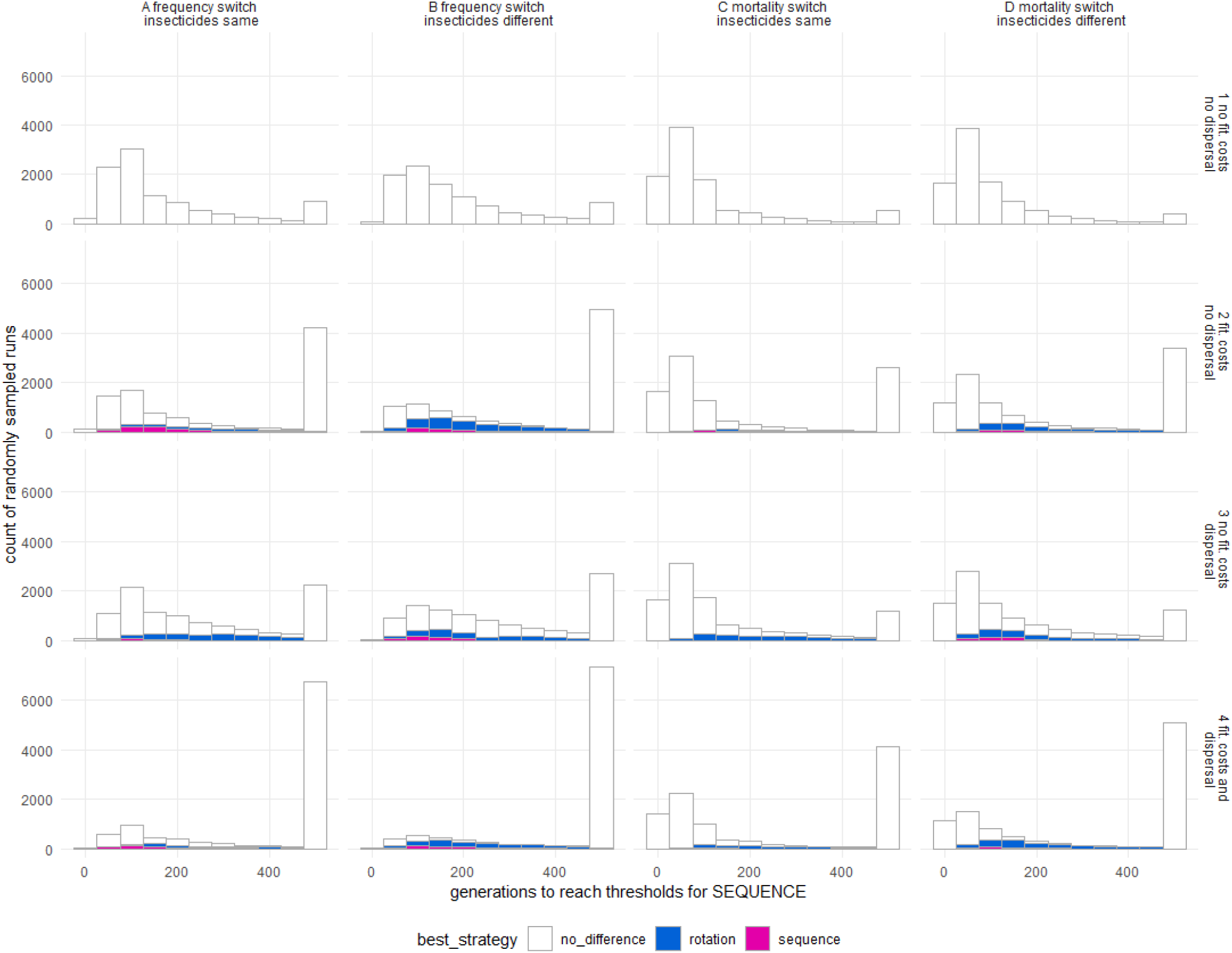
As for Figure 9 but histograms now use an Xaxis of generations to reach threshold for sequences.

The results from the sensitivity analyses are given in SI #2. The PRCC coefficients are low and, more importantly, there seems to be no parameters that consistently predict when one policy is favoured over another. The classifications trees also failed to identify any clear parameter space where one policy was favoured over another (Si #2).

In summary, neither policy is ever favoured under our base scenario of no fitness costs or gene flow with untreated refugia. When costs and/or refugia are present one policy may be favoured in some circumstances but the advantages were small (typically <10%) and there did not appear to be any consistent predictors of when one policy may be favoured over the other.

## 4. Discussion

The results presented above show that use of a rotation or sequential policy has little impact on how long an insecticide repertoire will last before resistance renders it ineffective. The best way to understand this result is to assume *a priori* that rotations have no effect, then to proceed from first principles. It would be expected, intuitively, that when there are no costs of resistance or refugia, the time to resistance would be the same for both policies: it is simply that resistance under sequential deployment evolves in a few, larger steps while resistance in rotations evolves in more numerous smaller steps: resistance in both policies reach the same endpoint (i.e. resistance to all insecticides in the armoury) at the same time. This is exactly what was observed (e.g. Figure 4) and our results show that, in the absence of costs or refugia, both policies last exactly the same amount of time (top row of Figure 2). The two policies may differentially affect the longevity of the insecticide armoury when costs and/or refugia are present (e.g. Figures 5 to 7). However, the differences are slight (typically <10%; Figures 2 and 3) and there seems to be no set of circumstances that consistently favour one policy over another (PRCC and classification trees in the SI).

Another way to understand these results is to recognise that “sequential” use or “rotations” of insecticides can both can be regarded as different types of rotations. In fact, there appears to be three distinct types of rotation in use and they are often confused. We clarify them as follows.

- “Rotate when resistant” is another description of our “sequential” strategy i.e. deploy one insecticide until resistance makes it ineffective and is then replaced by the next insecticide in the repertoire, and so on. Insecticides in our sequential deployment can (as will likely occur in practice) later be redeployed if the population has regained its susceptibility (through costs and/or gene flow with refugia). In essence, sequences can be regarded as “rotate when resistant” through the insecticide repertoire driven by the empirical need to replace an ineffective insecticide.
- “Pre-planned” rotations as considered above i.e. where periodic insecticide changes occur irrespective of local levels of resistance and is driven by the belief that this maximises the lifespan of the insecticide repertoire.

Realising that our “sequential” and “rotations” policies are essentially different types of rotations driven by different triggers for insecticide change helps explain the observation that there is rarely any significant difference in their ability to protect the insecticide armoury (Figures 2 and 3).

Another type of rotations can be added to the two types described above. It is an empirically driven rotation and can be defined as follows:

- “Pre-emptive rotations” occur when insecticides are changed when experience shows that a resistance threshold is almost certain to be reached by a certain point (e.g. by month 3 of the growing season) so a rotation is scheduled to avoid this threshold.

There are many factors in agriculture that drive a policy of frequently treating pest populations with a wide range of insecticides (Table 1). Frequent reapplication of the same insecticide is likely to rapidly drive resistance and it is likely that farmers, recognising this, pre-emptively rotate insecticides in anticipation of this resistance. Similarly, pre-emptive rotations may occur in public health. The best-known example in this context is control of black fly in rivers of Cameroon as a part of the Onchocerciasis control program [2, 21], where seasonal rotation is deployed as experience has shown that resistance rapidly builds up. In summary, it is likely that most rotations used in agriculture occur for operational reasons such as the need to treat different species or lifecycle stages or to minimise ecological impact (Table 1) or are pre-emptive rotations anticipating the likely onset of resistance. These pre-emptive rotations have driven the widespread belief that empirical evidence favours the use of pre-planned rotations as an IRM strategy. In fact, pre-emptive rotations are operationally similar to a sequential rotate-when-resistant strategy with insecticide change driven by resistance reaching a certain threshold, it is simply that rather than monitoring the frequency up to the threshold and then reacting, it is predicted in advance. In summary, pre-emptive rotations share the same philosophy as sequential use as both are driven by the same empirical need to avoid deploying ineffective insecticides and provide little empirical support for any putative benefit of pre-planned rotations over sequential use.

Most of the theoretical studies evaluating the use of IRM strategies (rotations, mosaics, mixtures) date from the 1980s. A recent review [2] cited two (in their Box 1 i.e. [22] and [23] to which we would add the contemporaneous work by Curtis [7] and Mani [24], neither of whom found any advantage of rotations over sequential use, Mani stating “Application of two insecticides alternately in time (commonly referred to as rotation), does not have any advantage over the use of one insecticide till resistance appears followed by a switch to another insecticide”. The only recent theoretical evaluation of rotations we could find was that of Busi et al [25] who modelled herbicide applications strategies. Four herbicides were available and the lifespan of herbicide repertoire was measured as “the number of years until resistance occurred, defined as the first year that crop yields fall below 75% of maximum”. Importantly, their simulation of “sequential” use assumed a single herbicide was used (their Table 2) until “resistance” was reached at which point the timespan was measured. This precluded the possibility of a more realistic sequential use (as used in our modelling) i.e. that one herbicide can be used until resistance is almost reached, then the next one deployed. It therefore, unrealistically, evaluates a single pesticide (their “sequential” deployment) against several pesticides used in a rotation or mixture. Under these assumptions it seems inevitable that sequential use of a single pesticide would fail before a rotation employing two or more pesticides.

Empirical studies on the advantages of rotations give similarly mixed results. The best known example for insecticides in public health was a comparison of IRM strategies to control malaria vectors in Mexico [26] and later e.g. [27]. The authors of these studies did not draw any explicit conclusion on which, if any, IRM policy was best (because resistnce evolved rapidly under all IRM polices). Subsequent authors have tried to draw conclusion which, for example, Dufour et al [2] summarised as follows: “a 3-compound annual rotation, a 2-compound mosaic, and a long-term treatment with a single insecticide were compared. An increase of resistance was observed in all villages, but it demonstrated that mosaic or rotation selected for low resistance levels and remained stable compared with villages with single treatment where resistance increased rapidly and to levels significantly greater”. As with the Busi et al paper, we argue that comparing a 3-compound rotation with a single-compound deployment is unrealistic, as a village with a failing compound could sequentially replace with the other two compounds in turns, so the study cannot be read as empirical support of rotations over sequential use. Finally we note the results of Legata and colleagues [28] who investigated rotations in experimental populations of Chlamydomonas and concluded that “the effects of herbicide cycling on the evolution of resistance may be more complex and less favorable than generally assumed” an empirical result that is clearly consistent with our modeling results.

One important practical reason for using (pre-planned) rotations is that it is operationally robust. It is likely to be operationally challenging to correctly manage a sequential deployment, i.e. to robustly detect that resistance has exceeded the switching threshold, and act on this information by rapidly replacing the current insecticide. If this is not operationally possible, then a pre-planned rotation minimises the time an ineffective insecticide continues to be deployed. In our comparisons, we assumed that insecticides in a sequence would be discontinued as soon as the switching threshold was reached. This is likely to be problematic in many resource-poor regions when there are likely to be challenges organising routine monitoring, acting on data suggesting resistance is present, and factors such as institutional inertia and extended timelines for ordering replacement insecticides. All these factors may delay deploying the next insecticide in a sequence with the result that ineffective insecticides are retained and used for significant amounts of time, possibly with associated failure of disease control. An illustration of how difficult it is to switch policy is illustrated by the antimalarial drug chloroquine which was widely known to be failing in most countries at the start of this century, but remained in widespread use until an article appeared in the Lancet accusing the funders (World Bank, WHO) of “medical malpractice” in funding its purchase [29]. This was inflammatory, but effective and drove the deployment of the new generation of antimalarial drugs, the ACTs. It is important to ease the process of removing and replacing ineffective insecticides and one way to maintain the agility to switch insecticides is to have pre-planned rotations already in place. As ever, Curtis [7] pre-empted this conclusion by several decades, stating that “If one considers that resistance to each available compound is inevitable eventually, there may be practical advantages for user organisations in procuring pre-planned quantities of two or more compounds, rather than having to make a switch in the crisis atmosphere of a control failure due to resistance”. It is usually recognised that the (potential) effectiveness of rotations as IRM depend on the presence of fitness costs (e.g. Dusfour et al [2], their Box 1) but costs are not required for rotations to be operational robust. Rotations, even in the absence of costs, may still be more operationally robust than sequences by eliminating the latter’s high resistance frequencies that occur as the failing insecticide is replaced by an effective replacement.

As ever in model of evolution of insecticide it is impossible to investigate all scenarios in a single paper. It is important to be explicit on this and we highlight the following:

- We assume resistance is encoded by single genes rather than being a polygenic trait. We do not include the potential effects of modifier genes acting on these single genes to reduce the costs of resistance alleles (It has been suggested that rotations could be favoured as a strategy because they keep resistance frequencies lower and restrict selection pressure for modifier genes [5]) as their inclusion would require more detailed calculations tracking linkage disequilibrium between each resistance locus and its modifier. However, there is little evidence to support the existence of such modifier genes in insecticide resistance apart from the well-known example of the Australian sheep blowfly, *Lucilia cuprina* [30]. Even with the existence of modifier genes it is unlikely that they would provide much of an advantage to rotations in the scenarios we describe. In the modelled scenarios the use of an insecticide is stopped when the resistance allele frequency for that insecticide reaches 0.5. Thus in all scenarios resistance frequencies do not remain at high levels for long. In this situation it seems unlikely that modifiers would be selected for. Earlier simulations by Curtis [7] similarly showed that rotations did not give a sizeable advantage over sequential use unless using a resistance threshold of 90% as the trigger to switch insecticide and very high (80%) fitness costs of resistance in the absence of the modifier. The presence of modifiers also relies absolutely on the assumption that significant fitness costs are associated with the mutations encoding IR and this is by no means certain (see recent critical review by ffrench-Constant and Bass [18]).
- We assume there is no cross-resistance between insecticides.
- We assume rotations is every 10 generations (roughly a year for the main malaria vectors). We see no reason why choice of rotation interval should affect the conclusion but intend to confirm this by examining rotations at 30 and 50 generations before formal submission.
- Finally, we stress that we model the evolution of resistance a single population, albeit with intervention and refugia areas. In reality, different counties, or even health regions within countries, may potentially choose to deploy different insecticides resulting a patchwork or “mosaic” of different insecticides with gene flow between mosquito populations in the different countries. Such mosaics have been proposed as IRMs but need to be addressed in a separate work. It is clear that IR does not respect international boundaries and that a co-ordinated international deployment strategies may be superior to local solutions but again, this is clearly not within the remit of the current work.

We stress that these are usual assumptions made in modelling but we highlight them as important extensions that could be addressed in subsequent pieces of work. What we have shown is that modelling does not support the use of rotations on theoretical grounds as an effective, robust IRM.

In summary, we show that any differences between rotations and sequential policies are invariably small, and that any benefits of one strategy over another do not appear to be associated with any particular set of circumstances. Certainly, rotations should not be seen as a panacea capable of significantly reducing the evolution of resistance over a wide range of deployment scenarios. However, pre-planned rotations are probably more operationally robust because changing a front line insecticide may be a lengthy process in resource-poor areas where most vector-borne diseases occur. In this, and previous work (e.g. [10]) we have essentially repeated earlier modelling by authors in the 1980s and 1990s and which are now largely overlooked. We added modern computational power to investigate much wider ranges of parameter space and interrogate this parameter spaces using modern sensitivity analysis methods but basically found the same results. As an example we end with a quoted from Chris Curtis in 1987 who concluded “though there may be valid practical reasons for using pre-planned rotations, population genetics does not give strong support to this policy as against other ways of using two or more pesticides” [7].

## Supporting information

Algebra of Rotations as a IRM strategy

PRCC and classification tree plots

## Acknowledgments

We thank Mark Hoppe for discussion on how insecticides are used in agriculture which formed the basis for Table 1.

## Notes

### Competing Interest Statement

The authors have declared no competing interest.

