## Supplementary material for "Evaluating rotations as a strategy to delay the evolution of insecticide resistance in vectors of human diseases": Algebra of Rotations as a IRM strategy

### Rotations as an IRM stratagem: a quantitative analysis.

Note that, wherever possible, we use the same nomenclature as in Levick et al [1].

#### *(1) Natural selection (fitness costs) in the refugia.*

There is an implicit assumption that males and females of the same genotype have the same fitness costs in both the untreated refugia and the intervention site (the latter is described below). The fitness costs are the same in males and females so

$$F_u^{f,r\prime} = F_u^{m,r\prime} = \frac{\left[F_u^{m,r} F_u^{f,r} w_-^{rr} + \left(F_u^{m,r} F_u^{f,s} + F_u^{m,s} F_u^{f,r}\right) 0.5\, w_-^{rs}\right]}{\bar{W}}$$

Equation 1

$$F_u^{f,s\prime} = F_u^{m,s\prime} = \frac{\left[\mathrm{F}_\mathrm{u}^{\mathrm{m,s}} \mathrm{F}_\mathrm{u}^{\mathrm{f,s}} \mathrm{w}_-^{\mathrm{ss}} + \left(\mathrm{F}_\mathrm{u}^{\mathrm{m,r}} \mathrm{F}_\mathrm{u}^{\mathrm{f,s}} + \mathrm{F}_\mathrm{u}^{\mathrm{m,s}} \mathrm{F}_\mathrm{u}^{\mathrm{f,r}}\right) 0.5\, \mathrm{w}_-^{\mathrm{rs}}\right]}{\bar{W}}$$

Equation 2

Where $\bar{W}$ is a normalising coefficient equal to the sum of the numerators, F represents the frequency of an allele, F being subscripted by 't' or 'u' to indicate the population (treated intervention site or untreated refugia respectively) and the first superscript denoting male (m) or female (f) with the second superscript 'r' or 's' denoting resistant or sensitive alleles. The prime indicates the frequencies after selection. The 0.5 enters the equation because half the gametes produced by the heterozygotes are of each allele type. The symbol w represents the fitness of the genotype where the superscript is the genotype (rr, rs or ss) and the subscript indicates whether it is in the treated intervention site (symbol +) or in the untreated refugia (symbol -) .

These fitness parameters, w, are constructed from the insecticide parameters given in Table 1 of the main text i.e.

$$\mathrm{w}_-^{\mathrm{ss}} = 1$$

$$\mathrm{w}_-^{\mathrm{rs}} = 1 - \text{"}dominance\ of\ resistance * \text{"}cost\ of\ resistnce\text{"}$$

$$\mathrm{w}_-^{\mathrm{rr}} = 1 - \text{"}cost\ of\ resistnce\text{"}$$

***(2) Insecticide selection in the treated area***

There is differential selection between the sexes in the treated population, so we track allele frequencies separately in the sexes. This requires an additional superscript in F to denote male (m) or female(f), resulting in in 4 equations in total.

We use the exposure terms in Table 1 of Levick et al [1], and the fitness of Table 2 of the same paper

As in Levick et al, we allow the mosquitoes to encounter three niches in the treated area i.e. encounters no insecticides (coverage within the treated areas may not be 100%) with probability $\propto_{-}^{m}$ , low concentration of insecticides with probability $\propto_{a}^{m}$ and high levels of insecticide with probability $\propto_{A}^{m}$ where the superscript 'm' indicated males, and is replaced by 'f' when females are considered. We do not utilise this ability to differentiate between high and low insecticide concentrations in the present study but wanted to make the methodology compatible with Levick et al.

First calculate the resistance allele frequency in adult males after selection

$$F_t^{m,r'} = F_t^{m,r} F_t^{f,r}[\propto_{-}^{m} \mathrm{w}_{-}^{\mathrm{rr}} + \propto_{a}^{m} \mathrm{w}_{\mathrm{a}}^{\mathrm{rr}} + \propto_{A}^{m} \mathrm{w}_{\mathrm{A}}^{\mathrm{rr}}] + \left(F_t^{m,r} F_t^{f,s} + F_t^{m,s} F_t^{f,r}\right) 0.5 [\propto_{-}^{m} \mathrm{w}_{-}^{\mathrm{rs}} + \propto_{a}^{m} \mathrm{w}_{\mathrm{a}}^{\mathrm{rs}} + \propto_{A}^{m} \mathrm{w}_{\mathrm{A}}^{\mathrm{rs}}]$$

Equation 3

Similarly, for the sensitive allele frequency

$$F_t^{m,s\prime} = F_t^{m,s} F_t^{f,s}[\propto_{-}^{m} \mathrm{w}_{-}^{\mathrm{ss}} + \propto_{a}^{m} w_{a}^{ss} + \propto_{A}^{m} w_{A}^{ss}] + \left(F_t^{m,r} F_t^{f,s} + F_t^{m,s} F_t^{f,r}\right) 0.5 [\propto_{-}^{m} \mathrm{w}_{-}^{\mathrm{rs}} + \propto_{a}^{m} \mathrm{w}_{\mathrm{a}}^{\mathrm{rs}} + \propto_{A}^{m} \mathrm{w}_{\mathrm{A}}^{\mathrm{rs}}]$$

Equation 4

These male allele frequencies are normalised as

$F_t^{m,r\prime} = \frac{F_t^{m,r\prime}}{W_m}$ and $F_t^{m,s\prime} = \frac{F_t^{m,s\prime}}{W_m}$

where $W_m$ is a normalising coefficient equal to the sum of Equation 3 and Equation 4.

Females differ from males only in their exposure parameters i.e. the superscript in the α terms in Equation 3 and Equation 4, so

$$F_t^{f,r'} = F_t^{m,r} F_t^{f,r}\left[\propto_{-}^{f} \mathrm{w}_{-}^{\mathrm{rr}} + \propto_{a}^{f} \mathrm{w}_{\mathrm{a}}^{\mathrm{rr}} + \propto_{A}^{f} \mathrm{w}_{\mathrm{A}}^{\mathrm{rr}}\right] + \left(F_t^{m,r} F_t^{f,s} + F_t^{m,s} F_t^{f,r}\right) 0.5 [\propto_{-}^{f} \mathrm{w}_{-}^{\mathrm{rs}} + \propto_{a}^{f} \mathrm{w}_{\mathrm{a}}^{\mathrm{rs}} + \propto_{A}^{f} \mathrm{w}_{\mathrm{A}}^{\mathrm{rs}}]$$

Equation 5

And

$$F_t^{f,s\prime} = F_t^{m,s} F_t^{f,s} [\propto_{-}^{f}\ \mathrm{w}_{-}^{\mathrm{ss}} + \propto_{a}^{f}\ w_{a}^{ss} + \propto_{A}^{f}\ \ w_{A}^{ss}] + \left(F_t^{m,r} F_t^{f,s} + F_t^{m,s} F_t^{f,r}\right) 0.5 [\propto_{-}^{f}\ \mathrm{w}_{-}^{\mathrm{rs}} + \propto_{a}^{f}\ \mathrm{w}_{\mathrm{a}}^{\mathrm{rs}} + \propto_{A}^{f}\ \mathrm{w}_{\mathrm{A}}^{\mathrm{rs}}\ ]$$

Equation 6

These female allele frequencies also have to be normalised as

$F_t^{f,r\prime} = \frac{F_t^{f,r\prime}}{W_f}$ and $F_t^{f,s\prime} = \frac{F_t^{f,s\prime}}{W_f}$

where $W_f$ is a normalising coefficient equal to the sum of Equation 5 and Equation 6.

Computational time can be decreased by noting that some factors are common to 2 or more equations so that factor can be calculated once and substituted

Coefficient #1 is common to all Equation 3 to Equation 6:

$$\left(F_t^{m,r} F_t^{f,s} + F_t^{m,s} F_t^{f,r}\right) 0.5$$

Coefficient 2 is common to Equation 3 and Equation 4:

$$[\propto_{-}^{m}\ \mathrm{w}_{-}^{\mathrm{rs}} + \propto_{a}^{m}\ \mathrm{w}_{\mathrm{a}}^{\mathrm{rs}} + \propto_{A}^{m}\ \mathrm{w}_{\mathrm{A}}^{\mathrm{rs}}\ ]$$

Coefficient 3 is common to Equation 5 and Equation 6:

$$[\propto_{-}^{f}\ w_{-}^{rs} + \propto_{a}^{f}\ w_{\mathrm{a}}^{rs} + \propto_{A}^{f}\ w_{\mathrm{A}}^{rs}\ ]$$

Finally, we explicitly state how these fitness parameters ‘w’ are constructed from the insecticide parameters given in Table 1 of the main text. Fitness of the SS genotype is easily calculated as.

$$\mathrm{w}_{\mathrm{A}}^{\mathrm{ss}} = 1 - "Effectiveness"$$

Subsequent calculations are a bit complex and described in Levick et al [1] Table 2 and on the bottom of their page 12. Briefly, the value $\mathrm{w}_{\mathrm{A}}^{\mathrm{ss}}$sets the lower base line (because fitness of the RS and RR genotypes cannot fall below that of the SS in the presence of insecticide). The other baseline is 1 because RR genotypes in the presence of insecticide cannot be more fit that the "reference fitness" i.e. fitness of SS in the absence of insecticide which is assigned the value of 1.

We calculate the selective advantage of the RR genotypes (compared to the SS), $\mathrm{s}_{\mathrm{A}}^{\mathrm{rr}}$ , as

$$\mathrm{s}_{\mathrm{A}}^{\mathrm{rr}} = \text{"effectiveness"} * \text{"}\mathit{resistance\ restoration}\text{"}$$

Which is used to calculate the fitness of the RR genotypes as

$$\mathrm{w}_{\mathrm{A}}^{\mathrm{rr}} = \mathrm{w}_{\mathrm{A}}^{\mathrm{ss}} + s_{A}^{RR}$$

As can be seen by substitution, setting resistance restoration to have a value of 1 makes the fitness of the RR genotype equal to 1 (i.e. the RR genotype is completely unaffected by contact with insecticide) while setting it to zero means the RR genotype has the same fitness as the SS genotype. The SR genotype has a fitness determined by the dominance of the resistance allele i.e.

$$\mathrm{w}_{\mathrm{A}}^{\mathrm{rs}} = \mathrm{w}_{\mathrm{A}}^{\mathrm{ss}} + \text{"}\mathit{dominance\ of\ resistance}\text{"} * s_{A}^{RR}$$

***(3) Non-selection in the treated area***

This occurs when the locus under consideration is not being selected because the insecticide to which it encodes resistance is no longer deployed (i.e. has been rotated out or is not currently used in the sequential deployment). We still have to simulate the dynamics of that locus in the intervention site and can do so using Equation 1 and Equation 2 noting that the subscript in the 'F' terms will have to change from "u" to "t".

**(4) Dispersal**

We need three additional parameters (not in Levick et al) to incorporate dispersal.

- C is "coverage" of the intervention defined as the proportion of mosquitoes that are covered by the intervention (and 1-C is the proportion of the population in the untreated refugia).
- $r_t$ is the dispersal rate into and out of the treated area. It is the proportion of the treated population that disperses. We assume that immigration=emigration.
- An "effective dispersal rate, $\theta_e$, described below.

Obviously, it is the number of mosquitoes dispersing into and of out of the refuge that must be equal, not a proportion. So having defined the dispersal rate as a proportion $r_t$ of the treated areas, we can calculate their equivalent proportion of dispersers in/out the untreated refugia population as

$$r_u = r_t \frac{C}{1-C}$$

Equation 7

For example (and to confirm it works), assume there are 1,000 mosquitoes in the treated area and 9,000 in the refugia, so C=0.1. A proportion $r_t$ =0.05 disperse from the treated area so 0.05*1,000= 50 mosquitoes. The equivalent proportion in the refugia is 0.05*1000/9000=0.0055555. Multiplying by the size of the refugia (i.e. 0.0055555x9000=50) gives 50 mosquitoes coming into the refugia.

Note that there is a biologically plausible upper limit to dispersal which depends on the relative sizes of the treated and refugia populations. For example, if refugia and treatment each constitute 50% of the population then the upper limit of dispersal is 50% i.e. 50% stay and 50% leave and this constitutes random mixing of the population across treated and refugia areas (any values higher than 50% imply mosquitoes are actively leaving the treated area). Conversely, if the treated area is 90% of the proportion then it is implausible to have a dispersal rate of 50% as it assume 0.5*0.9= 45% of the population moves into the 10% of refugia. Algebraically, the dispersal rate that represents free movement is $r_t$=(1-C)

Having different limits of dispersal makes it difficult to incorporate into sensitivity analyses (because it is constrained by another parameter, C, which also varies). Hence, we have a new parameter $\theta_e$ which is the effective dispersal rate and lies between zero, representing no dispersal, and 1, representing free movement. Hence

$$r_t = (1-C)\theta_e$$ Equation 8.

And $r_u$ is obtained by substituting this value into equation i.e.

$$r_u = r_t \frac{C}{1-C} = \frac{(1-c)\theta_e C}{1-C} = \theta_e C$$ Equation 9

It is $\theta_e$ (and C) that are taken as inputs to the model rather than $r_t$ directly.

The allele frequencies after dispersal are designated by a double prime and are the frequencies next generation. They are computed as follows.

Male resistance allele frequency in the intervention site after dispersal:

$$F_t^{m,r''} = (1 - r_t)F_t^{m,r'} + r_t F_u^{m,r'}$$

Equation 10

Female resistance allele frequency in the intervention site after dispersal:

$$F_t^{f,r''} = (1 - r_t)F_t^{f,r'} + r_t F_u^{f,r'}$$

Equation 11

Male resistance allele frequency in refugia after dispersal:

$$F_u^{r''} = (1 - r_u)F_u^{m,r'} + r_u F_t^{m,r'}$$

Equation 12

Female resistance allele frequency in refugia after dispersal:

$$F_u^{r''} = (1 - r_u)F_u^{f,r'} + r_u F_t^{f,r'}$$

Equation 13

The corresponding four sensitive allele frequencies are computed as 1 minus resistance allele frequency.

There are two implicit assumptions built into this dispersal complement.

- Firstly, we assume equal dispersal rates for males and females. In principle we could define $\theta_e$ separately for each gender and use them in Equations 8 to 13. This would be relatively easy but we will stick to the current assumption of no gender difference.

- Secondly, we assume equal numbers of mosquitoes immigrate and emigrate from the intervention site. The objection goes something like this "if we are hammering the population in the intervention site, won't there be few mosquitoes emigrating and a big influx of immigrants". Our first inclination was to include a demographic impact parameter (d) into Equations 8 and 9 but it is really difficult to estimate the demographic impact of insecticide deployment. Population genetic methodology, as used here, tracks allele frequencies and ignores the population sizes in which these frequencies exist so it would mean adding a whole new demographic component that would increase the model complexity and number of input parameters. Demography is a whole new subject and would need a model of population regulation, which will change as resistance spreads (see discussion in [2]). In summary, we don't think we need this new parameter and can argue as follows. Coverage can be (re)defined as the proportion of the population in the intervention site after the intervention has occurred e.g. if the intervention covered 14% of the original population and halved the population in the intervention sites, then coverage now becomes 7% (actually, being picky, its 7/(7+86)= 7.5%). We don't actually define coverage as being before or after intervention and, in practice, we probably have very little idea of its value in the field. Our inclination is therefore to ignore demographic impact, argue it is incorporated into coverage, and maybe revisit if we find coverage has an impact on the results.

1. Levick B, South A, Hastings IM. A Two-Locus Model of the Evolution of Insecticide Resistance to Inform and Optimise Public Health Insecticide Deployment Strategies. PLoS Comp Biol. 2017;13(1):e1005327. doi: 10.1371/journal.pcbi.1005327.
2. Barbosa S, Kay K, Chitnis N, Hastings IM. Modelling the impact of insecticide-based control interventions on the evolution of insecticide resistance and disease transmission. Parasites & Vectors. 2018;11(1):482. doi: 10.1186/s13071-018-3025-z.
