## Supplementary material for "Evaluating rotations as a strategy to delay the evolution of insecticide resistance in vectors of human diseases": PRCC and classification tree plots

2020-05-18

The results shown on Figures 7 and 8 of the main text were subjected to PRCC and decision tree analysis. To recap, the analysis was as follows i.e.

Column A: each insecticide had identical properties; switching criterion was resistance allele frequency exceeding 50% Column B: each insecticide had different properties; switching criterion was resistance allele frequency exceeding 50% Column C: each insecticide had identical properties; switching criterion was mortality <90% Column D: each insecticide had different properties; switching criterion was mortality <90%

Row 1: no costs of resistance or dispersal into/out of an untreated refugia
Row 2: costs are present; untreated areas absent
Row 3: costs absent; refugia present of an untreated refugia
Row 4: costs and refugia and both present

PRCC

The results are given in the graphs at the end of this SI. The wealth of data they contain meant the most convenient method of evaluating the results is to print out each page, physically arrange them by column/row and search for consistent patterns within rows or within columns. It is also possible to compare consistency between evaluation criteria in each row/column by comparing evaluation based on operational lifespan of the insecticide armoury (red crosses) with evaluation based on mean mortality during this lifespan (blue crosses); note that lifespan data are absent in plots of A1 and C1 as each policy had identical results in these circumstances.

There may be some slight patterns (e.g. "coverage" across row 3, "effectiveness" in column C) but these may well be coincidental and there seem no clear, consistent patterns that would identify circumstances under which one policy is consistently favoured over the other.

The results given in D3 and D4 are the most important as they reflect the most realistic deployment scenarios i.e. insecticides differ in their properties, switching is based on mortality), but PRCC values are small and inconsistent both within the plots (i.e. evaluation based on operation lifespan or on mean mortality) or between analyses of D3 and D4.

Finally, note the 'p' value may be very low (the sample size is very large) so the effects are 'real' but small in magnitude and inconsistent across scenarios

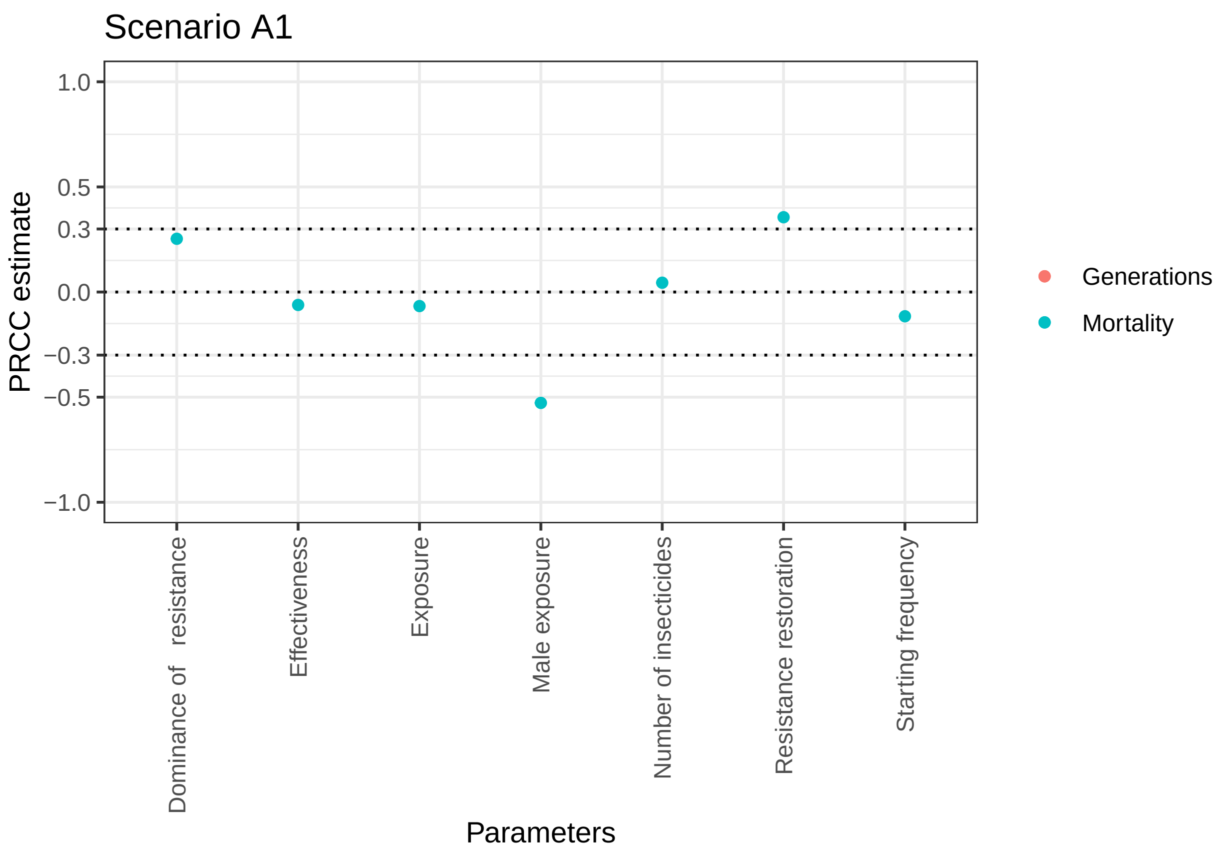


```
## [1] "P values for generations"
##    Number of insecticides             Effectiveness
##                       NaN                       NaN
##    Resistance restoration Dominance of   resistance
##                       NaN                       NaN
##                  Exposure             Male exposure
##                       NaN                       NaN
##        Starting frequency
##                       NaN
## [1] "P values for mortality"
##    Number of insecticides             Effectiveness
##              2.279497e-05              4.132992e-09
##    Resistance restoration Dominance of   resistance
##             2.839076e-271             5.796504e-134
##                  Exposure             Male exposure
##              1.597436e-10              0.000000e+00
##        Starting frequency
##              1.962586e-28
```

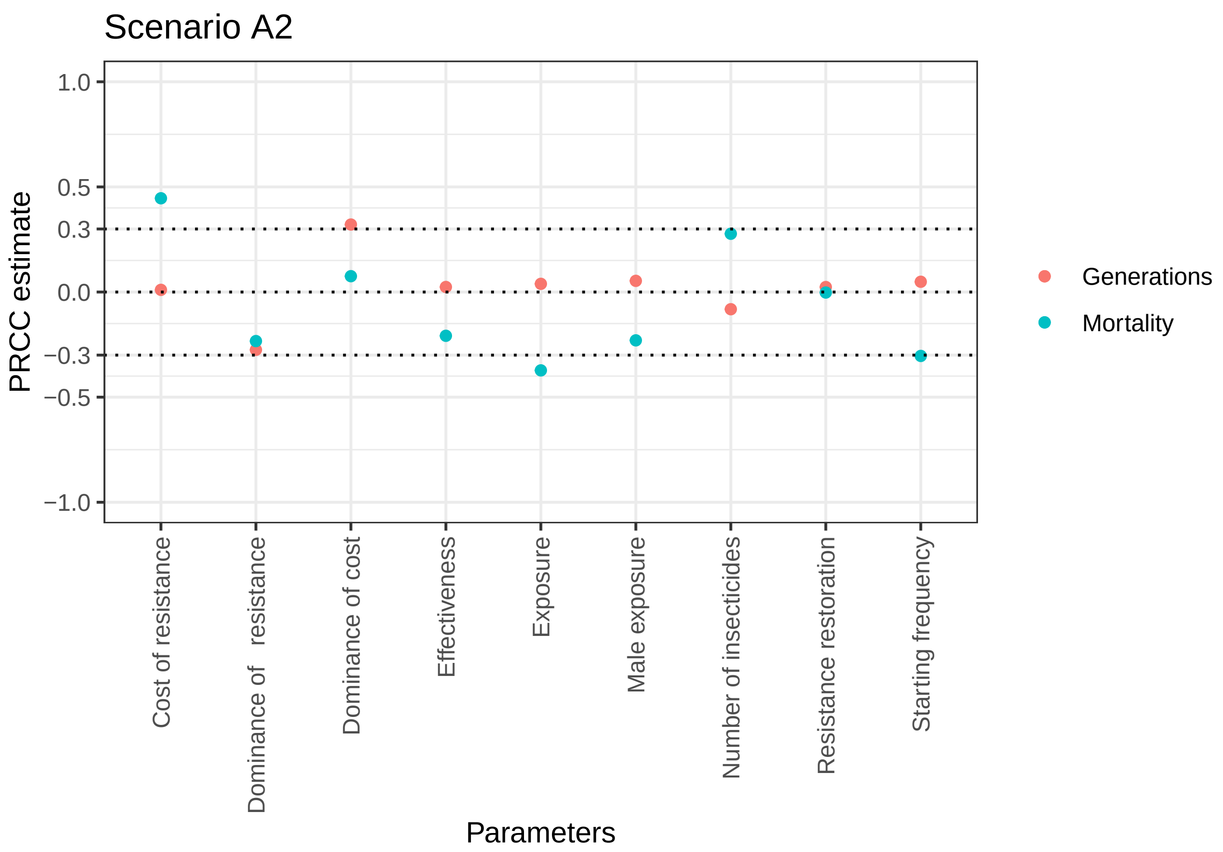


```
## [1] "P values for generations"
##    Number of insecticides             Effectiveness
##              4.431292e-10              6.866538e-02
##    Resistance restoration Dominance of   resistance
##              7.066883e-02             3.410289e-101
##                  Exposure             Male exposure
##              2.987057e-03              5.003460e-05
##        Starting frequency        Cost of resistance
##              2.009100e-04              4.302566e-01
##         Dominance of cost
##             2.149941e-139
## [1] "P values for mortality"
##    Number of insecticides             Effectiveness
##             6.256138e-103              6.819722e-58
##    Resistance restoration Dominance of   resistance
##              8.500737e-01              1.204792e-72
##                  Exposure             Male exposure
##             1.070692e-190              1.688966e-70
##        Starting frequency        Cost of resistance
##             2.058805e-124             6.201080e-282
##         Dominance of cost
##              8.977118e-09
```

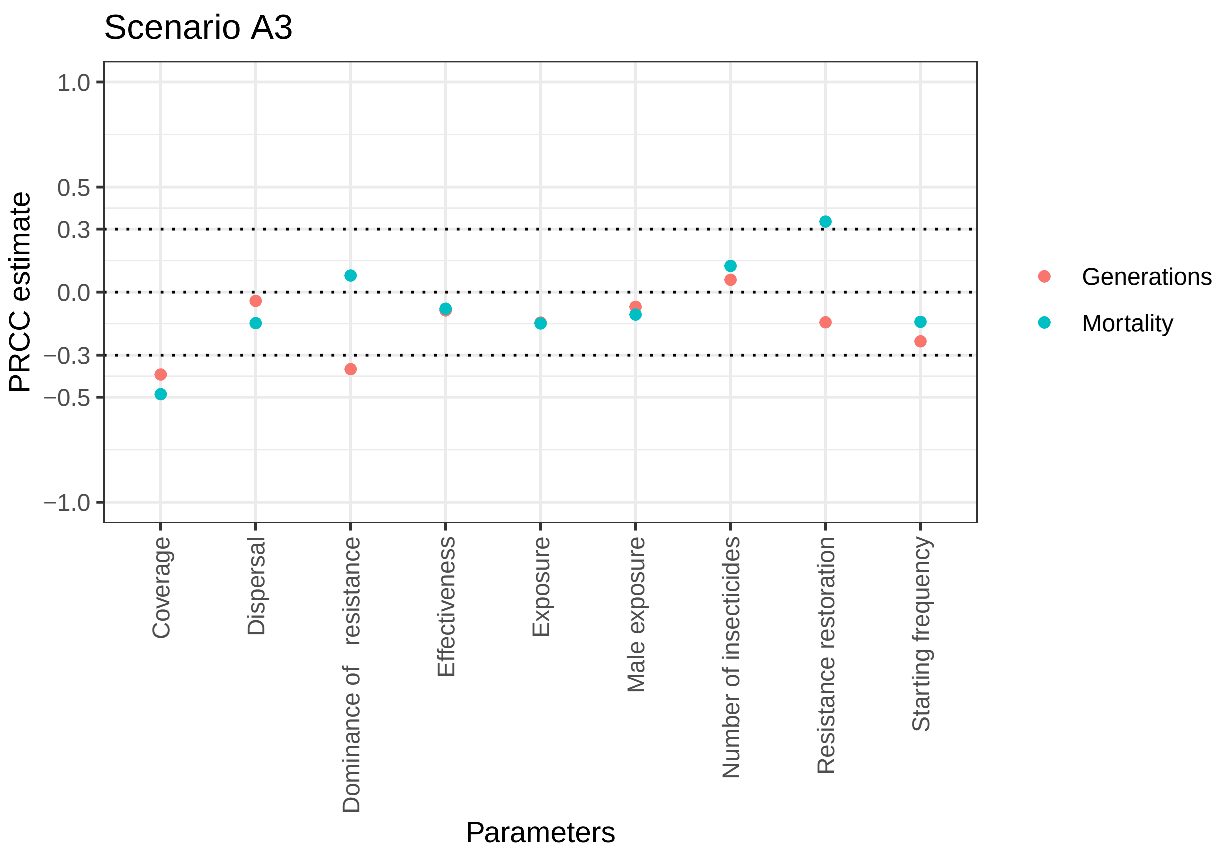


```
## [1] "P values for generations"
##    Number of insecticides             Effectiveness
##              2.040190e-07              8.847019e-15
##    Resistance restoration Dominance of   resistance
##              1.544334e-37             1.590978e-248
##                  Exposure             Male exposure
##              3.102177e-38              5.230439e-10
##        Starting frequency                  Coverage
##              2.436760e-98             3.112878e-287
##                 Dispersal
##              2.048830e-04
## [1] "P values for mortality"
##    Number of insecticides             Effectiveness
##              1.736910e-28              2.438733e-12
##    Resistance restoration Dominance of   resistance
##             4.530652e-206              2.569953e-12
##                  Exposure             Male exposure
##              2.352183e-40              1.960913e-21
##        Starting frequency                  Coverage
##              1.982413e-36              0.000000e+00
##                 Dispersal
##              1.339645e-39
```

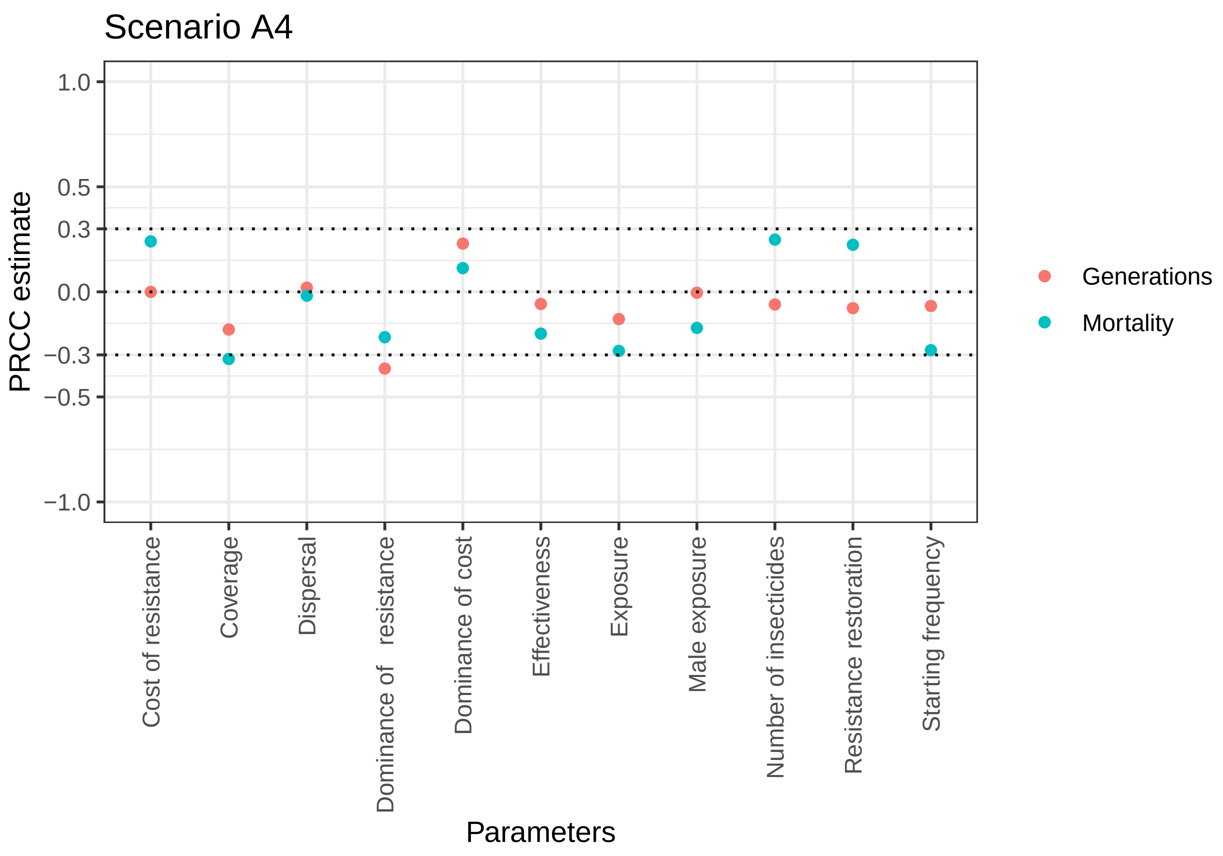


```
## [1] "P values for generations"
##    Number of insecticides             Effectiveness
##              5.964994e-04              9.664349e-04
##    Resistance restoration Dominance of   resistance
##              9.390170e-06             1.232597e-103
##                  Exposure              Male exposure
##              1.023258e-13              8.119258e-01
##        Starting frequency        Cost of resistance
##              1.217907e-04              9.860850e-01
##         Dominance of cost                  Coverage
##              2.460829e-40              5.473628e-25
##                 Dispersal
##              2.552781e-01
## [1] "P values for mortality"
##    Number of insecticides             Effectiveness
##              3.420232e-47              1.749062e-30
##    Resistance restoration Dominance of   resistance
##              1.830962e-38              1.030718e-35
##                  Exposure              Male exposure
##              2.399366e-60              5.004423e-23
##        Starting frequency        Cost of resistance
##              7.936141e-59              4.081867e-44
##         Dominance of cost                  Coverage
##              9.055371e-11              1.993505e-78
##                 Dispersal
##              2.970499e-01
```

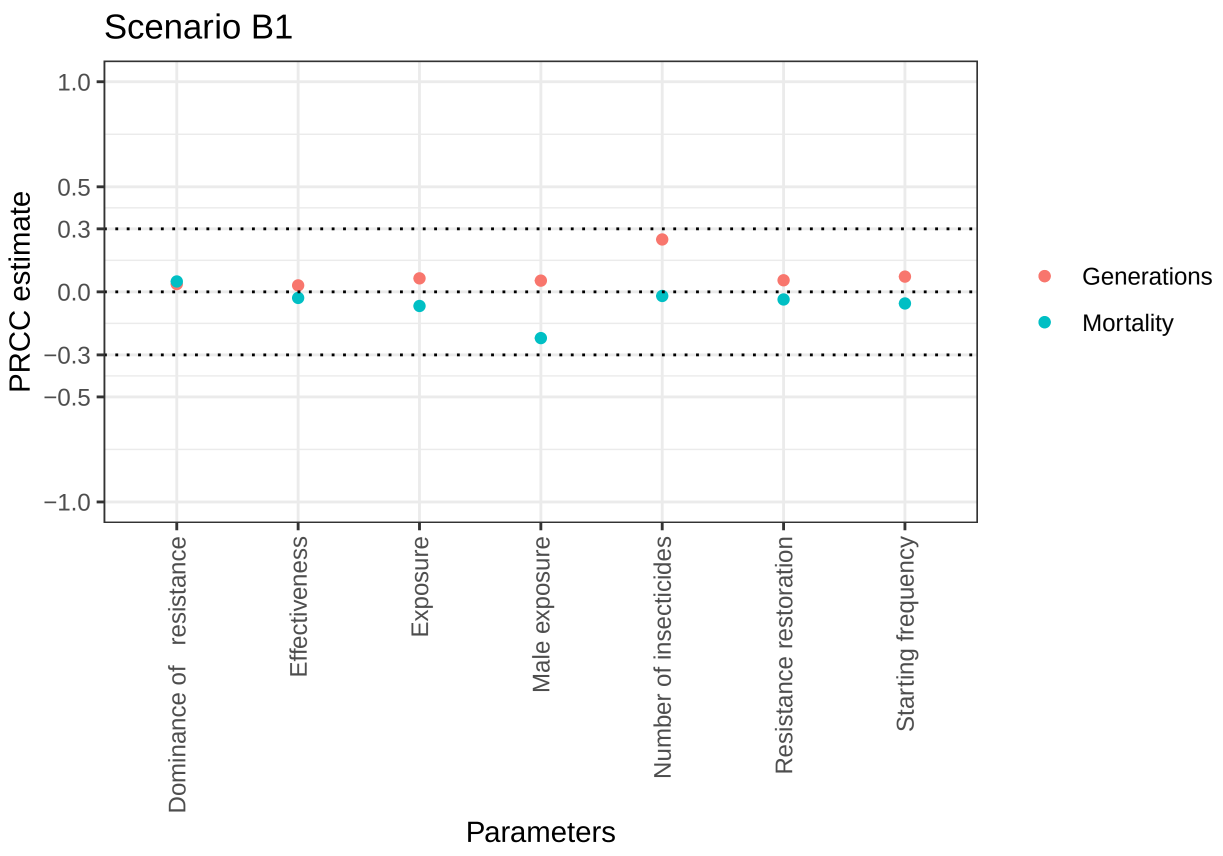


```
## [1] "P values for generations"
##    Number of insecticides              Effectiveness
##             2.793885e-130               3.186603e-03
##    Resistance restoration Dominance of   resistance
##              1.227383e-07               3.781611e-04
##                  Exposure              Male exposure
##              6.593703e-10               2.926757e-07
##        Starting frequency
##              3.391630e-12
## [1] "P values for mortality"
##    Number of insecticides              Effectiveness
##              5.986633e-02               5.692249e-03
##    Resistance restoration Dominance of   resistance
##              5.154546e-04               2.034030e-06
##                  Exposure              Male exposure
##              1.120616e-10              3.306465e-101
##        Starting frequency
##              1.193420e-07
```

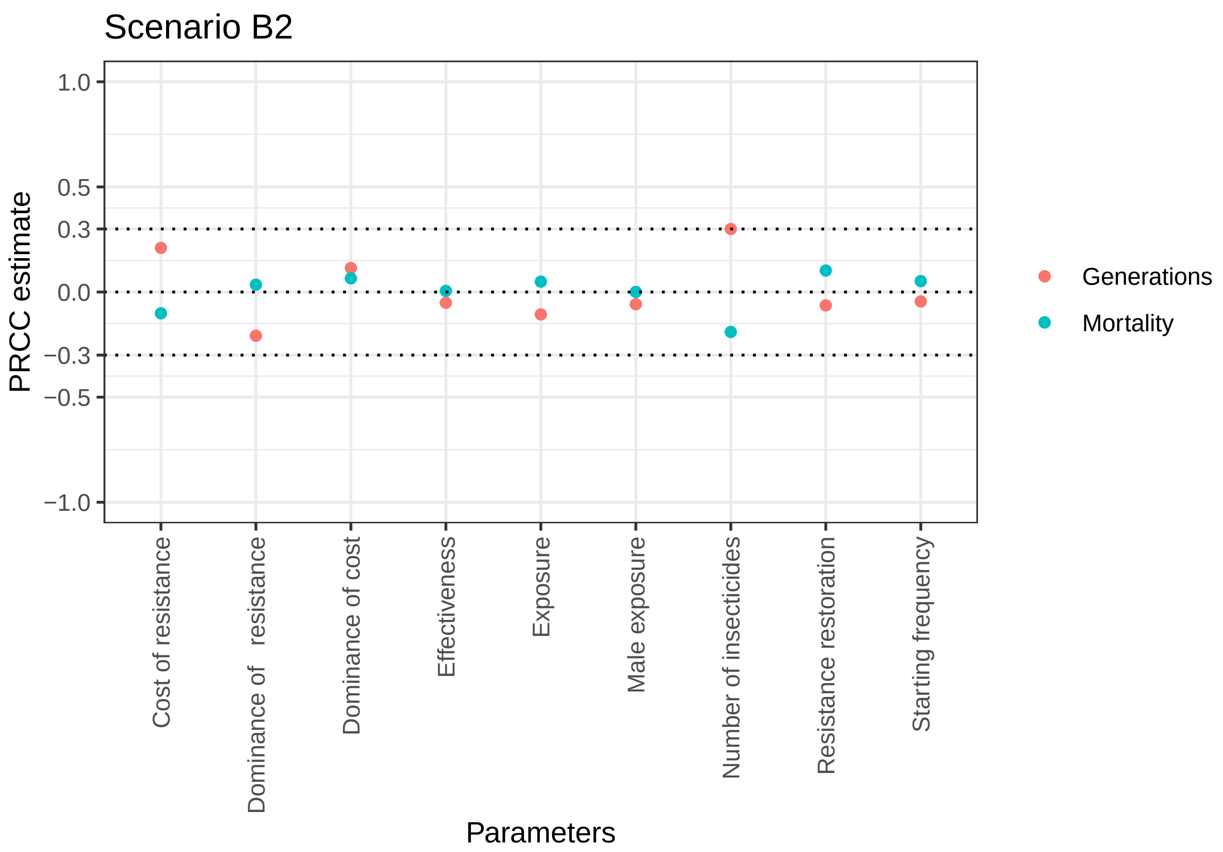


```
## [1] "P values for generations"
##    Number of insecticides             Effectiveness
##             3.445174e-107              2.530469e-04
##    Resistance restoration Dominance of   resistance
##              5.221642e-06              3.489091e-51
##                  Exposure             Male exposure
##              2.074492e-14              3.151796e-05
##        Starting frequency        Cost of resistance
##              1.292085e-03              2.661099e-52
##         Dominance of cost
##              2.604509e-16
## [1] "P values for mortality"
##    Number of insecticides             Effectiveness
##              7.385269e-43              6.990974e-01
##    Resistance restoration Dominance of   resistance
##              2.016125e-13              1.221363e-02
##                  Exposure             Male exposure
##              3.963614e-04              9.717417e-01
##        Starting frequency        Cost of resistance
##              1.914308e-04              3.310871e-13
##         Dominance of cost
##              3.074197e-06
```

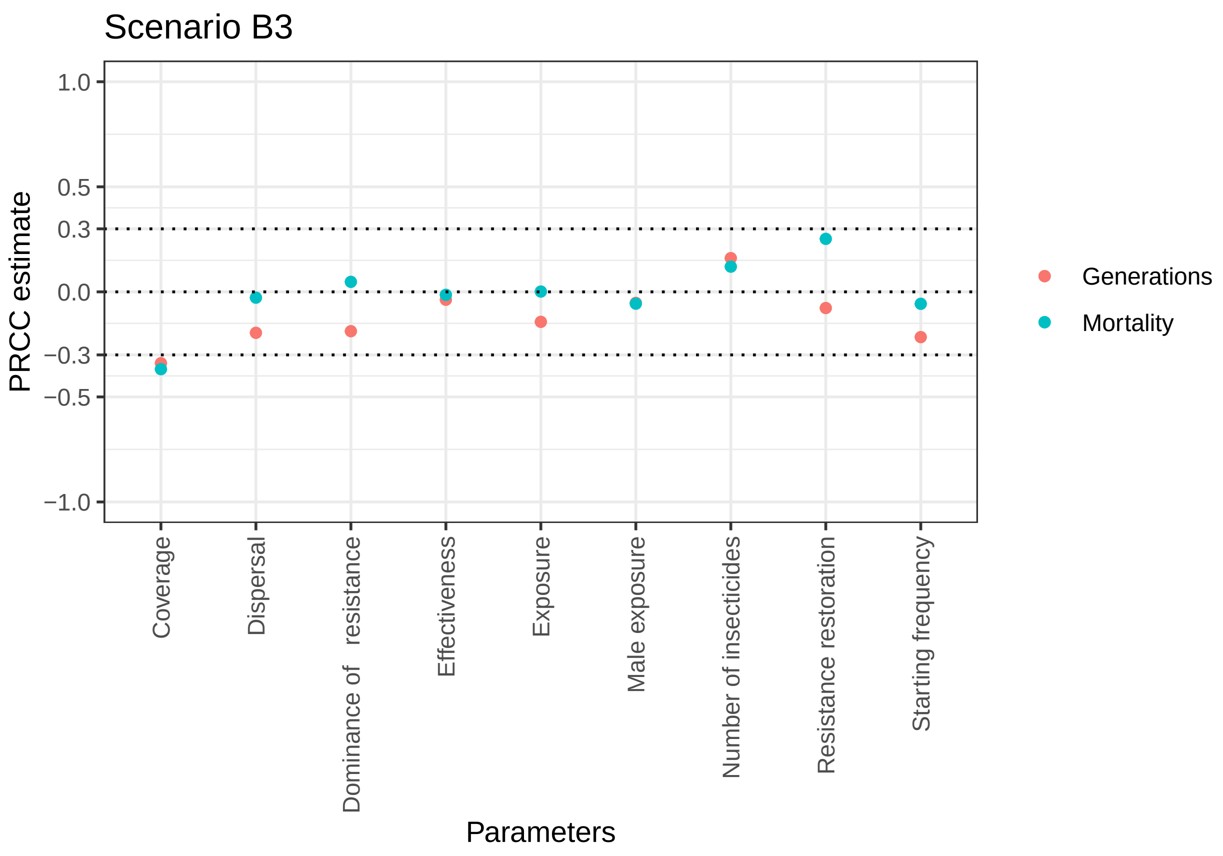


```
## [1] "P values for generations"
##    Number of insecticides             Effectiveness
##              4.621072e-44              1.190614e-03
##    Resistance restoration Dominance of   resistance
##              3.178174e-11              1.321768e-59
##                  Exposure             Male exposure
##              4.243877e-35              5.891333e-06
##        Starting frequency                  Coverage
##              1.081656e-78             7.213789e-200
##                 Dispersal
##              1.513218e-64
## [1] "P values for mortality"
##    Number of insecticides             Effectiveness
##              2.922073e-25              2.194519e-01
##    Resistance restoration Dominance of   resistance
##             4.420165e-108              4.123880e-05
##                  Exposure             Male exposure
##              8.941555e-01              1.221618e-06
##        Starting frequency                  Coverage
##              9.166678e-07             6.332364e-237
##                 Dispersal
##              1.705509e-02
```

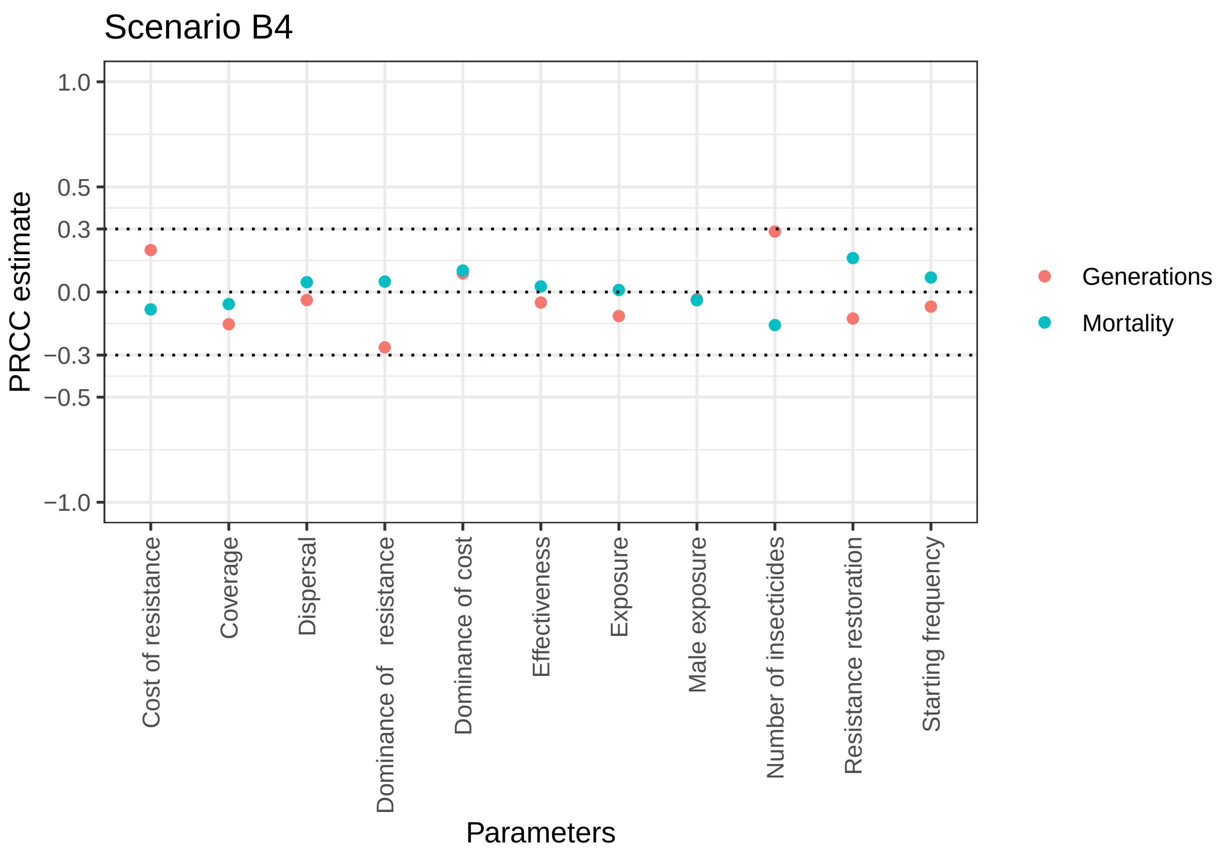


```
## [1] "P values for generations"
##    Number of insecticides            Effectiveness
##              3.091741e-52             9.523047e-03
##    Resistance restoration Dominance of   resistance
##              5.946935e-11             9.285600e-44
##                  Exposure            Male exposure
##              2.692993e-09             7.693886e-02
##        Starting frequency       Cost of resistance
##              3.210231e-04             1.814614e-25
##         Dominance of cost                 Coverage
##              4.636453e-06             1.513121e-15
##                 Dispersal
##              4.841748e-02
## [1] "P values for mortality"
##    Number of insecticides            Effectiveness
##              2.574257e-16             1.705323e-01
##    Resistance restoration Dominance of   resistance
##              3.938262e-17             1.015848e-02
##                  Exposure            Male exposure
##              6.186243e-01             4.336015e-02
##        Starting frequency       Cost of resistance
##              3.418948e-04             1.734081e-05
##         Dominance of cost                 Coverage
##              1.140939e-07             2.918941e-03
##                 Dispersal
##              1.555878e-02
```

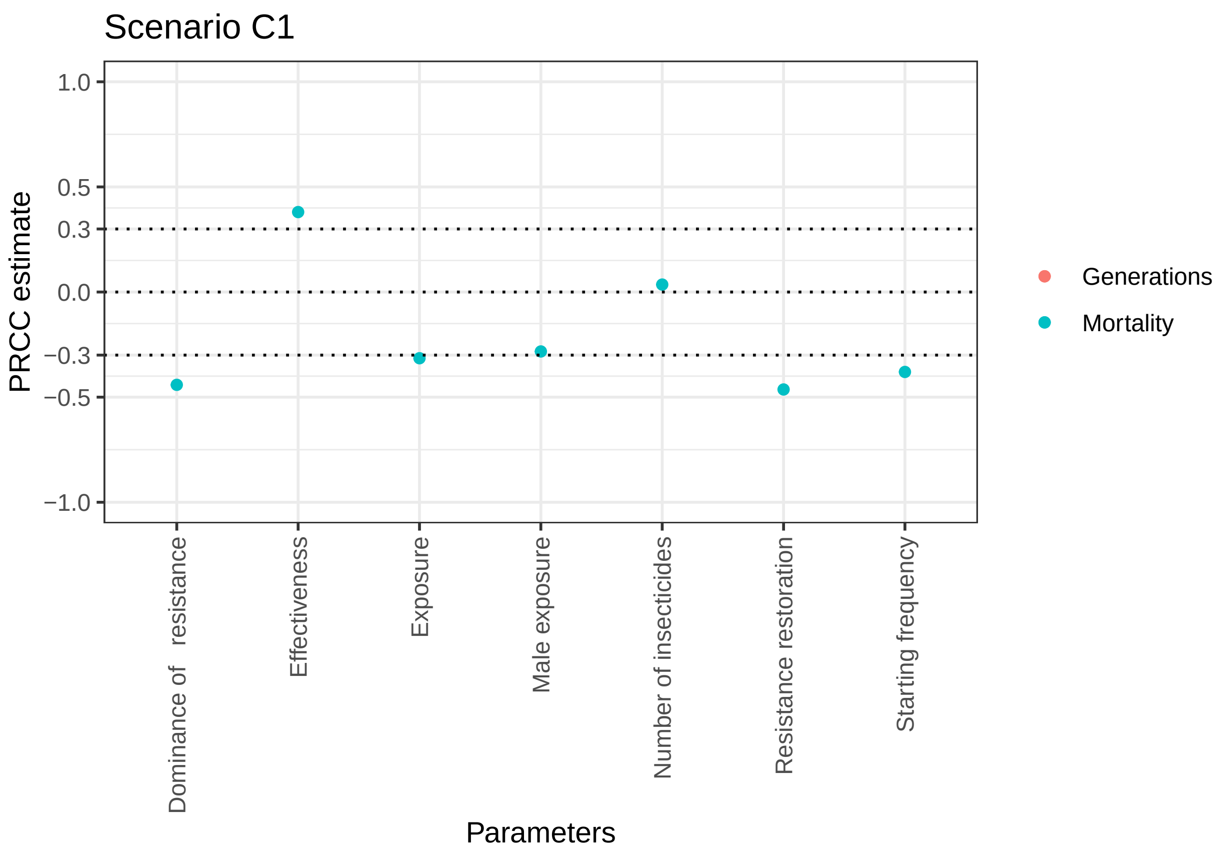


```
## [1] "P values for generations"
##    Number of insecticides              Effectiveness
##                       NaN                        NaN
##    Resistance restoration Dominance of   resistance
##                       NaN                        NaN
##                  Exposure              Male exposure
##                       NaN                        NaN
##        Starting frequency
##                       NaN
## [1] "P values for mortality"
##    Number of insecticides              Effectiveness
##              5.536947e-04               0.000000e+00
##    Resistance restoration Dominance of   resistance
##              0.000000e+00               0.000000e+00
##                  Exposure              Male exposure
##             2.891056e-217              9.843181e-174
##        Starting frequency
##             1.976263e-323
```

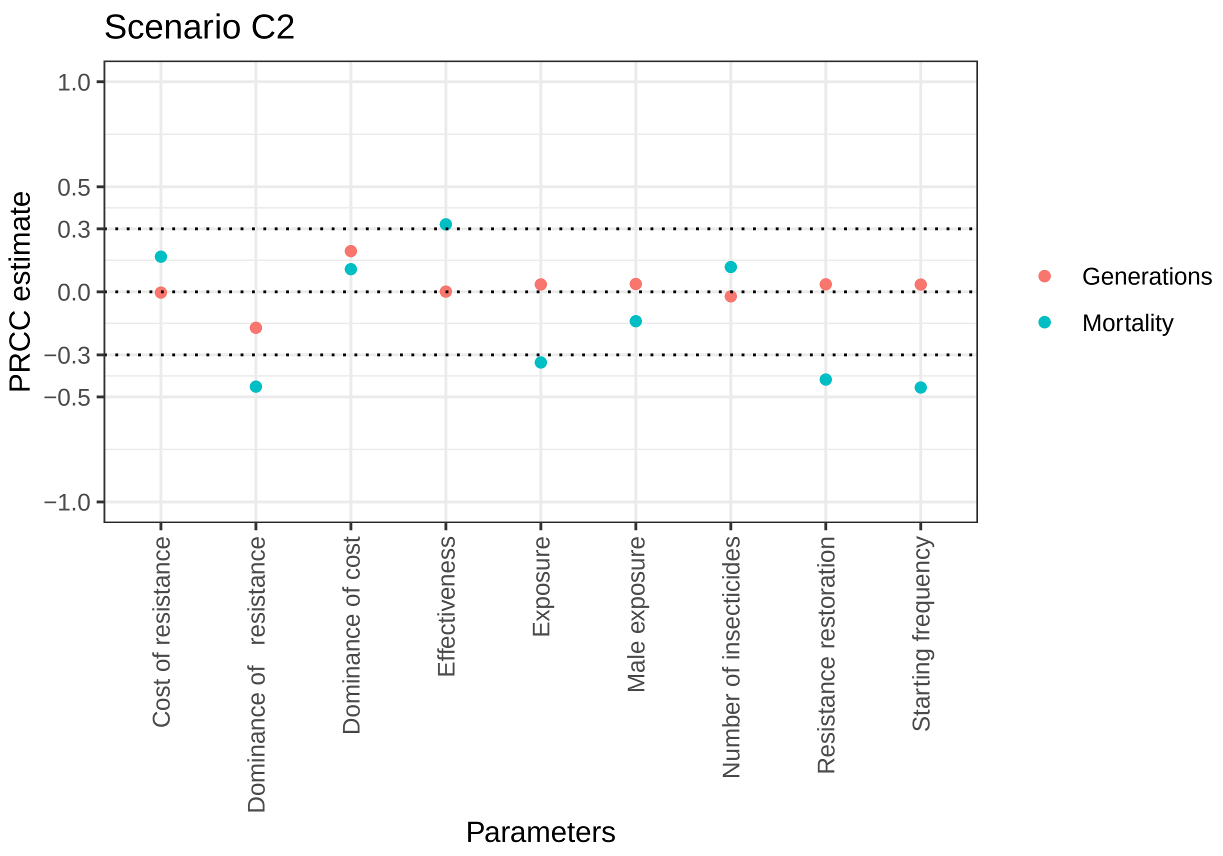


```
## [1] "P values for generations"
##    Number of insecticides             Effectiveness
##              6.613208e-02              9.126458e-01
##    Resistance restoration Dominance of   resistance
##              2.107584e-03              9.689983e-50
##                  Exposure             Male exposure
##              2.363406e-03              1.312496e-03
##        Starting frequency        Cost of resistance
##              2.947348e-03              7.551858e-01
##         Dominance of cost
##              1.364583e-63
## [1] "P values for mortality"
##    Number of insecticides             Effectiveness
##              1.901525e-24             3.162762e-177
##    Resistance restoration Dominance of   resistance
##             1.467414e-309              0.000000e+00
##                  Exposure             Male exposure
##             9.265694e-195              1.352644e-33
##        Starting frequency        Cost of resistance
##              0.000000e+00              1.442660e-47
##         Dominance of cost
##              1.341460e-20
```

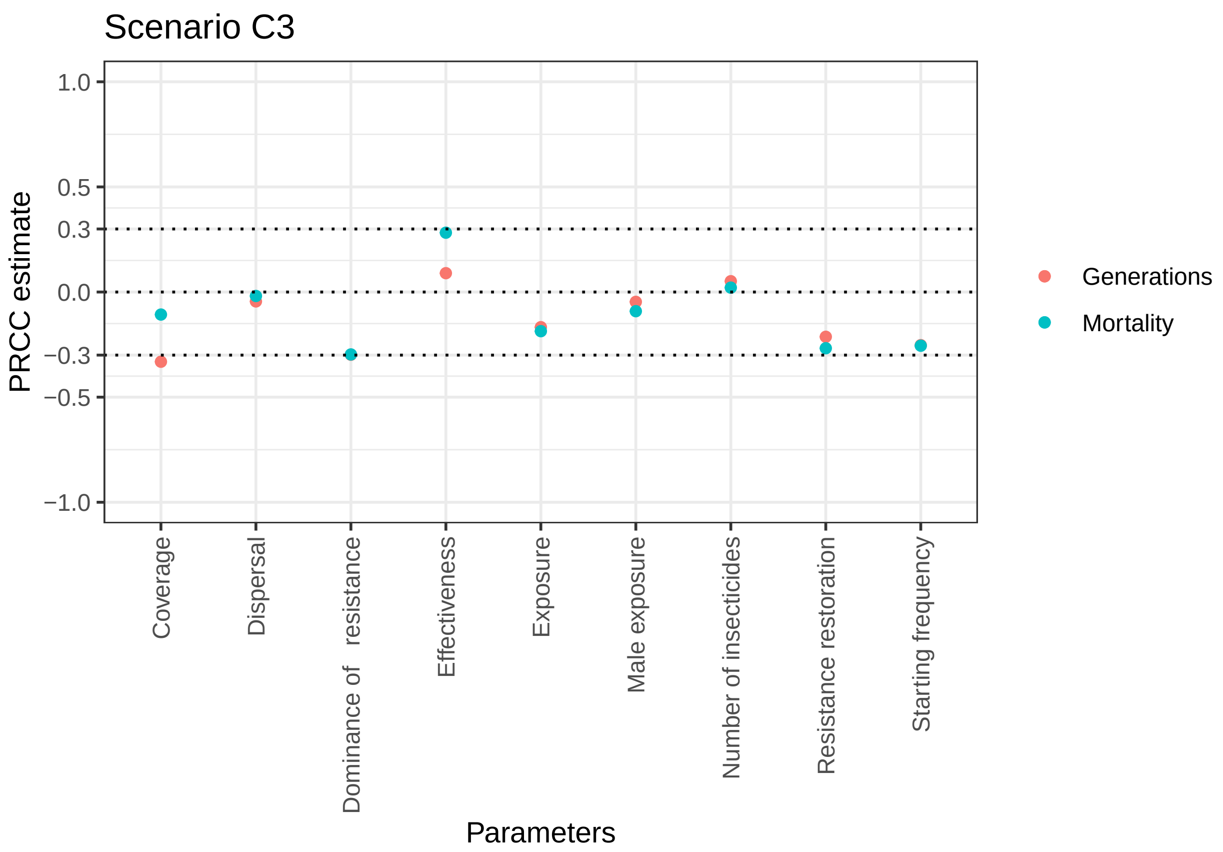


```
## [1] "P values for generations"
##    Number of insecticides             Effectiveness
##              1.324957e-06              3.097366e-17
##    Resistance restoration Dominance of   resistance
##              5.149826e-91             6.974852e-181
##                  Exposure             Male exposure
##              2.268646e-56              1.139877e-05
##        Starting frequency                  Coverage
##             9.857767e-129             1.725897e-225
##                 Dispersal
##              2.580508e-05
## [1] "P values for mortality"
##    Number of insecticides             Effectiveness
##              4.962626e-02             2.666263e-161
##    Resistance restoration Dominance of   resistance
##             2.505642e-144             4.098326e-179
##                  Exposure             Male exposure
##              9.857408e-70              9.155748e-18
##        Starting frequency                  Coverage
##             1.562930e-131              5.436548e-24
##                 Dispersal
##              7.645491e-02
```

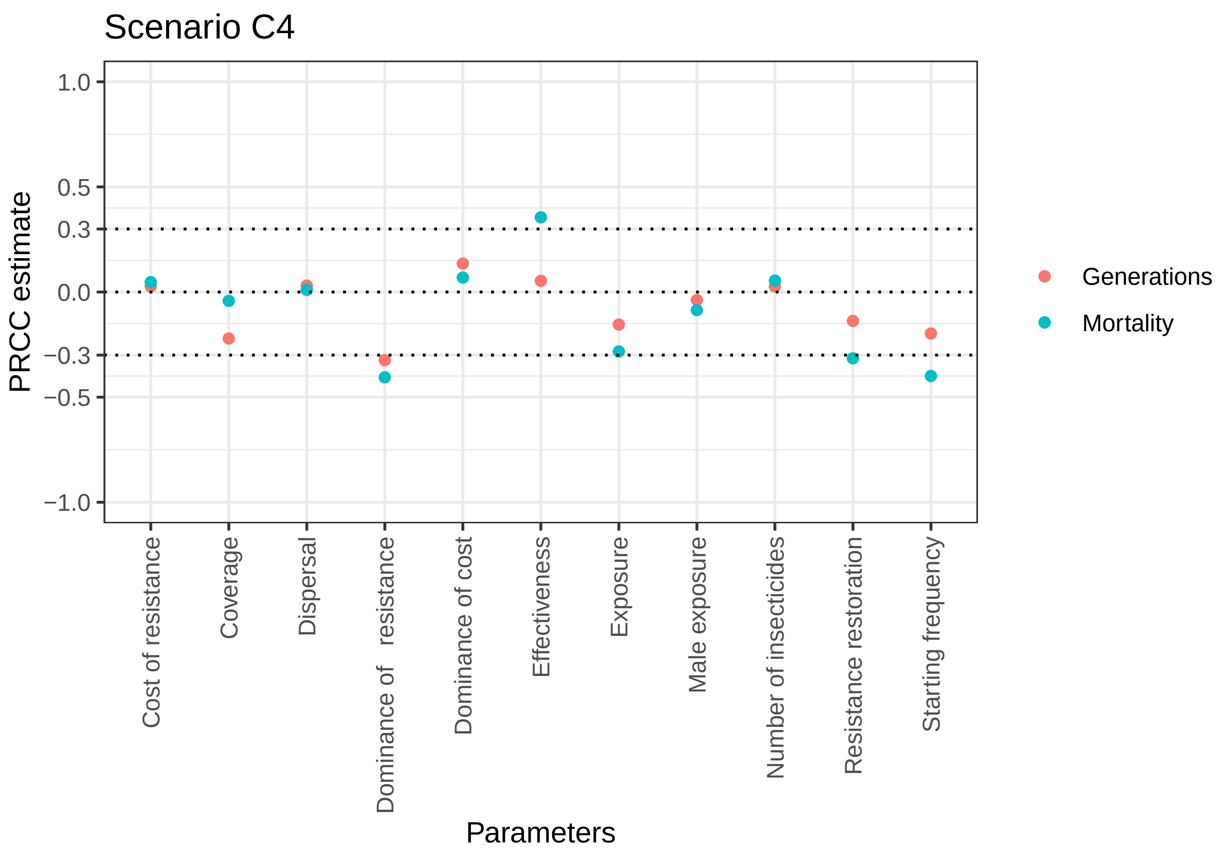


```
## [1] "P values for generations"
##    Number of insecticides              Effectiveness
##              4.121770e-02               4.043657e-05
##    Resistance restoration Dominance of   resistance
##              2.535163e-26              7.777280e-145
##                  Exposure              Male exposure
##              5.690995e-33               3.447412e-03
##        Starting frequency         Cost of resistance
##              7.900528e-53               3.439374e-02
##         Dominance of cost                   Coverage
##              1.318700e-25               2.240292e-66
##                 Dispersal
##              1.772746e-02
## [1] "P values for mortality"
##    Number of insecticides              Effectiveness
##              2.831730e-05              2.018771e-175
##    Resistance restoration Dominance of   resistance
##             1.446793e-136              2.082715e-232
##                  Exposure              Male exposure
##             5.502352e-109               3.536926e-11
##        Starting frequency         Cost of resistance
##             4.439919e-225               3.034425e-04
##         Dominance of cost                   Coverage
##              1.179055e-07               1.230631e-03
##                 Dispersal
##              4.419448e-01
```

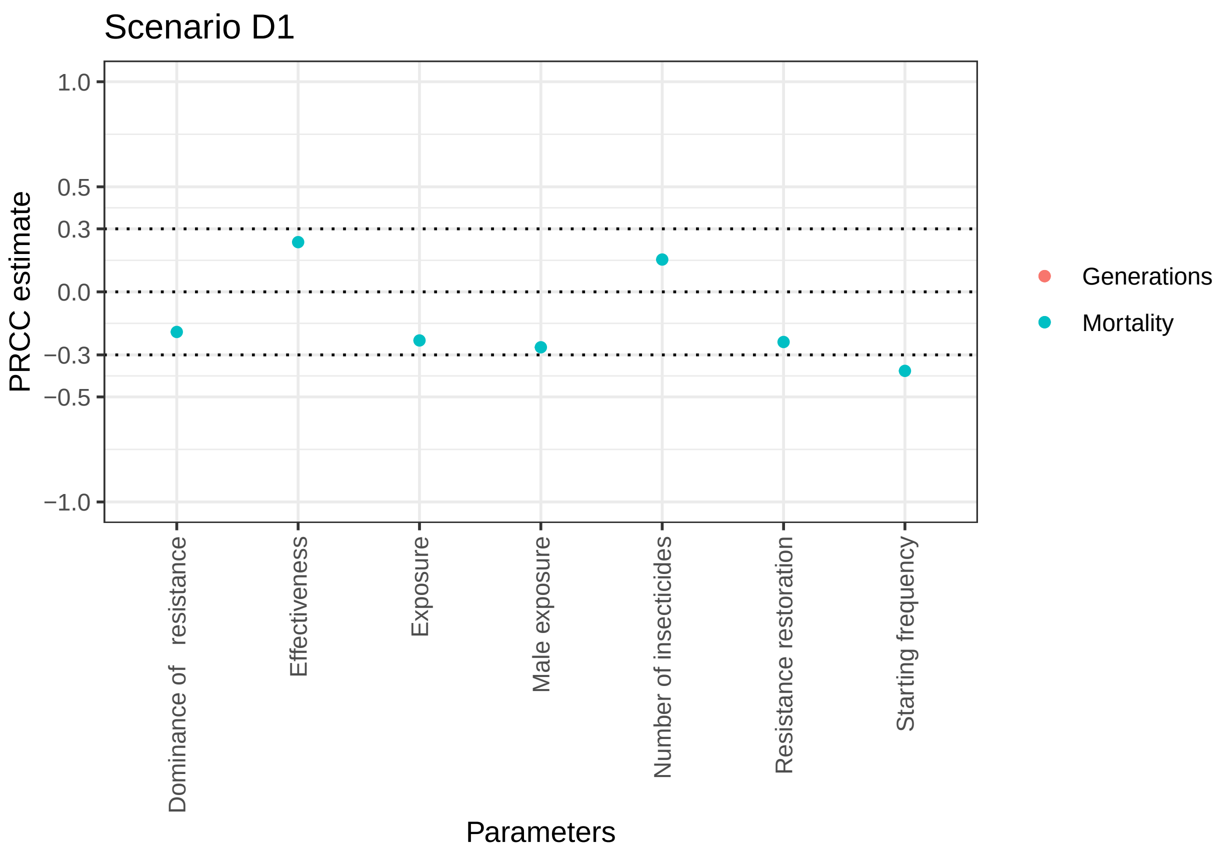


```
## [1] "P values for generations"
##    Number of insecticides             Effectiveness
##                       NaN                       NaN
##    Resistance restoration Dominance of   resistance
##                       NaN                       NaN
##                  Exposure             Male exposure
##                       NaN                       NaN
##        Starting frequency
##                       NaN
## [1] "P values for mortality"
##    Number of insecticides             Effectiveness
##              6.242765e-52             1.633581e-122
##    Resistance restoration Dominance of   resistance
##             1.296183e-124              1.561774e-79
##                  Exposure             Male exposure
##             7.049997e-117             9.553510e-153
##        Starting frequency
##             1.035562e-320
```

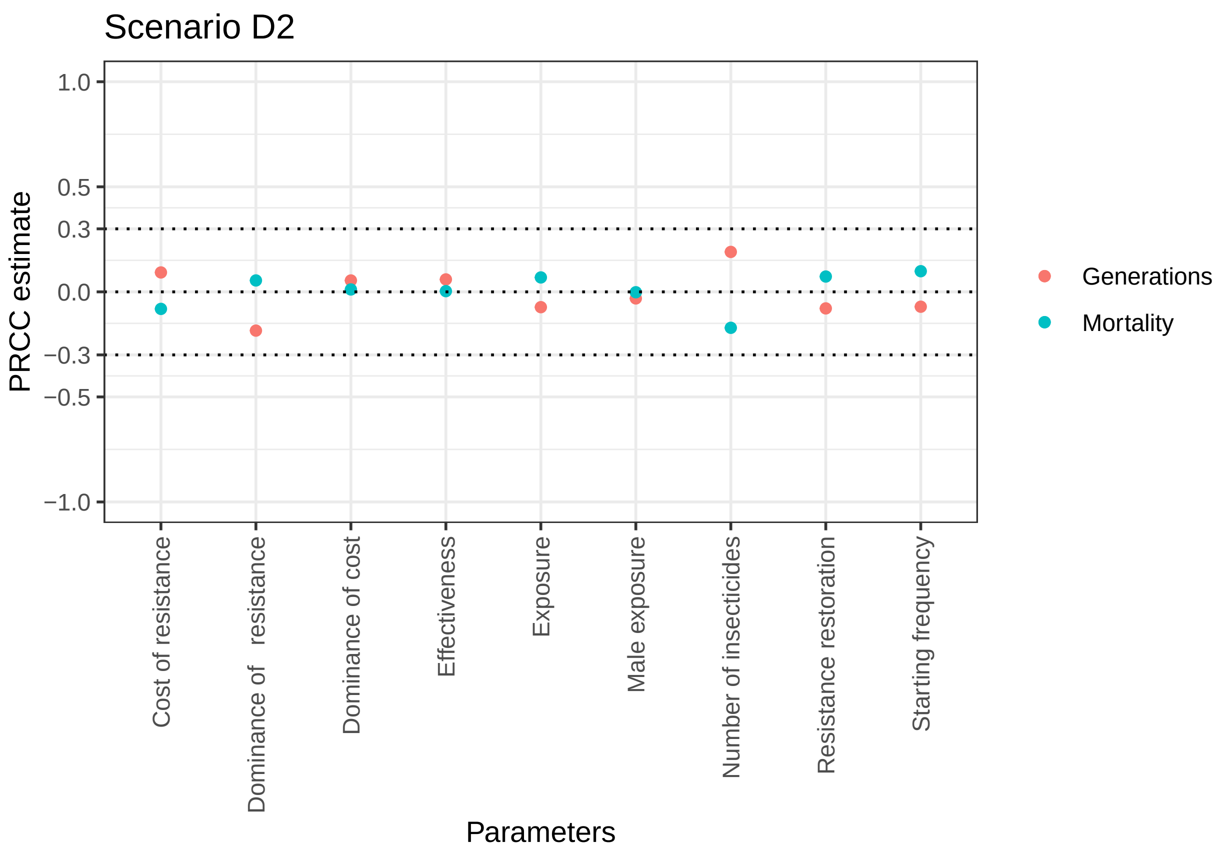


```
## [1] "P values for generations"
##    Number of insecticides            Effectiveness
##              8.200503e-55             1.443596e-06
##    Resistance restoration Dominance of   resistance
##              1.441594e-10             1.201341e-51
##                  Exposure            Male exposure
##              2.943605e-09             1.116616e-02
##        Starting frequency       Cost of resistance
##              8.701257e-09             5.228213e-14
##         Dominance of cost
##              1.053013e-05
## [1] "P values for mortality"
##    Number of insecticides            Effectiveness
##              1.325812e-44             7.853500e-01
##    Resistance restoration Dominance of   resistance
##              2.859244e-09             1.031419e-05
##                  Exposure            Male exposure
##              2.370490e-08             8.829040e-01
##        Starting frequency       Cost of resistance
##              1.159202e-15             3.963494e-11
##         Dominance of cost
##              3.563846e-01
```

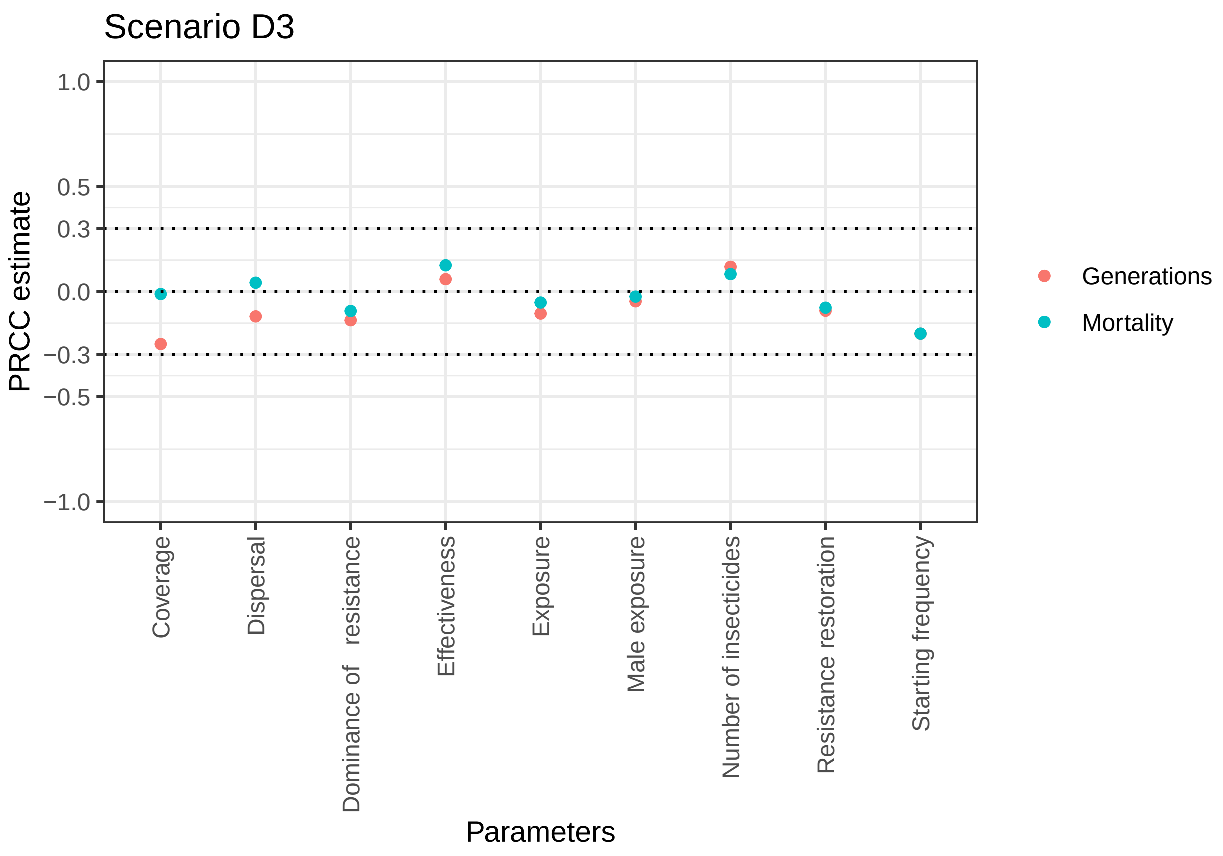


```
## [1] "P values for generations"
##    Number of insecticides             Effectiveness
##              7.434045e-29              2.582068e-08
##    Resistance restoration Dominance of   resistance
##              9.351446e-18              6.528407e-38
##                  Exposure             Male exposure
##              6.660790e-23              1.426250e-05
##        Starting frequency                  Coverage
##              9.760751e-81             1.531881e-125
##                 Dispersal
##              9.664162e-29
## [1] "P values for mortality"
##    Number of insecticides             Effectiveness
##              4.154050e-15              2.993334e-32
##    Resistance restoration Dominance of   resistance
##              5.412344e-13              3.481640e-18
##                  Exposure             Male exposure
##              9.576041e-07              2.529119e-02
##        Starting frequency                  Coverage
##              2.029387e-80              2.767899e-01
##                 Dispersal
##              6.645464e-05
```

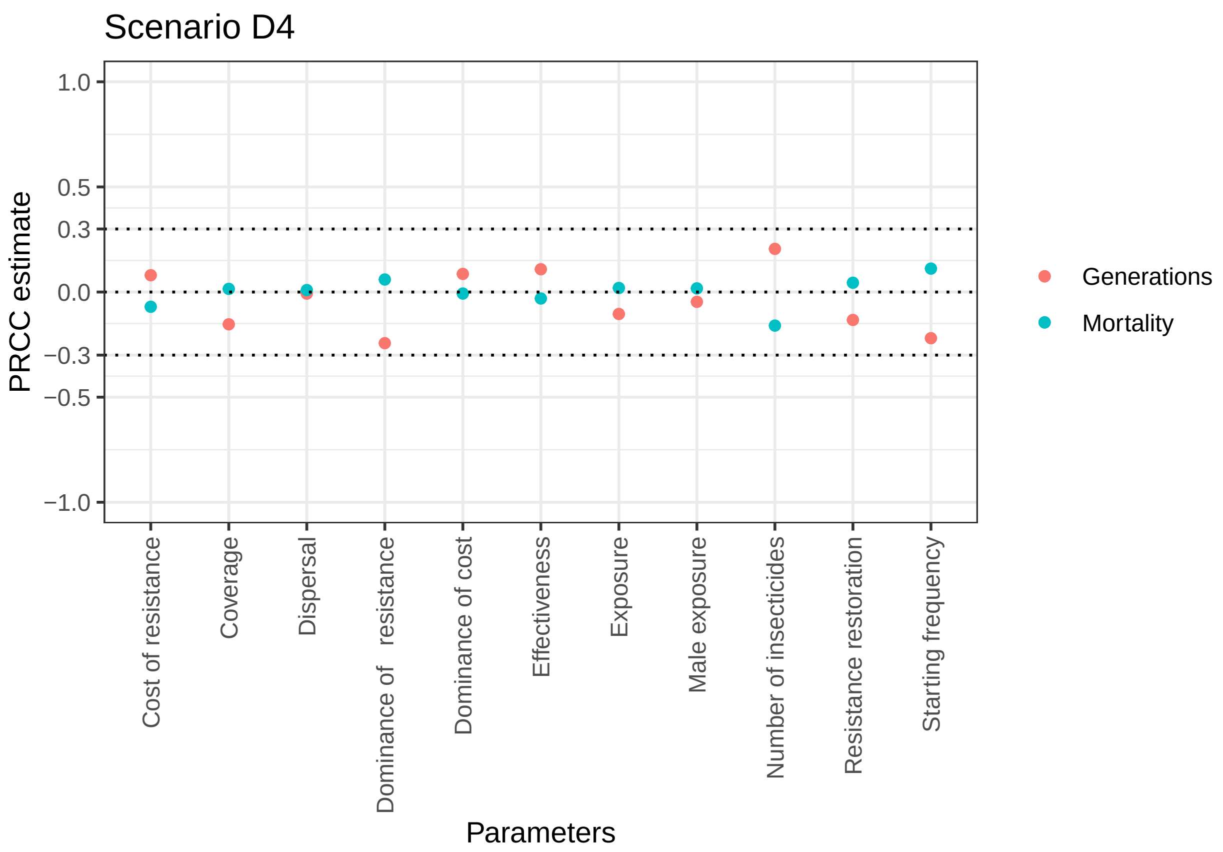


```
## [1] "P values for generations"
##    Number of insecticides            Effectiveness
##              7.191772e-48             2.713605e-14
##    Resistance restoration Dominance of   resistance
##              8.983791e-21             4.919087e-67
##                  Exposure            Male exposure
##              2.272904e-13             1.101868e-03
##        Starting frequency       Cost of resistance
##              1.097601e-54             2.038884e-08
##         Dominance of cost                 Coverage
##              1.484036e-09             2.970703e-27
##                 Dispersal
##              6.020441e-01
## [1] "P values for mortality"
##    Number of insecticides            Effectiveness
##              2.143110e-29             3.130298e-02
##    Resistance restoration Dominance of   resistance
##              1.966945e-03             2.879619e-05
##                  Exposure            Male exposure
##              1.735534e-01             2.238551e-01
##        Starting frequency       Cost of resistance
##              4.868563e-15             8.983808e-07
##         Dominance of cost                 Coverage
##              6.133150e-01             2.946694e-01
##                 Dispersal
##              5.137393e-01
```

Classification trees

Classification trees were run on the same datasets, using the R package Rpart. When evaluated on generations, those runs with a difference of 10 generations or less were regarded as “equivalent”. A significant subset of runs evaluated on mortality were also equivalent as they had the same overall mortality under both policies (as might be expected if both policies reached the same endpoint but by different routes). The trees are interpreted as follows: • The numbers under each box give the numbers of runs in the box which were equivalent, favoured rotations, favours sequences, in that order. • The percentage given below each box is the percentage of runs that fall into that box

As for PRCC, the only reasonable way to look for consistency seems to require printing the graphs, spreading them on the floor in columns/rows as in Figs 7&8 in main ms and look for consistent patterns across row and down columns. There was a tendency for low exposure to favour rotations in Column A when evaluated on mean moratility but in general there was no consistent patterns.

As noted above, scenario A3 and A4 reflect the most realistic deployment scenarios but there was little consistence y between their trees.

Even when classification did occur they classes did not consistent identify parameter spaces in which one policy was clearly favoured. For example, Scenario D4 evaluated on mortality was relatively simple and identified parameter spaces in which sequences, rotations or equivalence was most likely. However, the numbers beneath each box show that although one policy or equivalence tended to be favoured, there were a large number of runs in the classification box which were contrary to the favoured policy/equivalence. For example, rotations was the favoured policy in the bottom right box of this figure with 134 run favouring rotations but the classification box still contained 68 runs where the policies were equivalent and 91 runs where sequences were favoured.

```
[1] "Scenario A1 Generations tree cannot run"
```

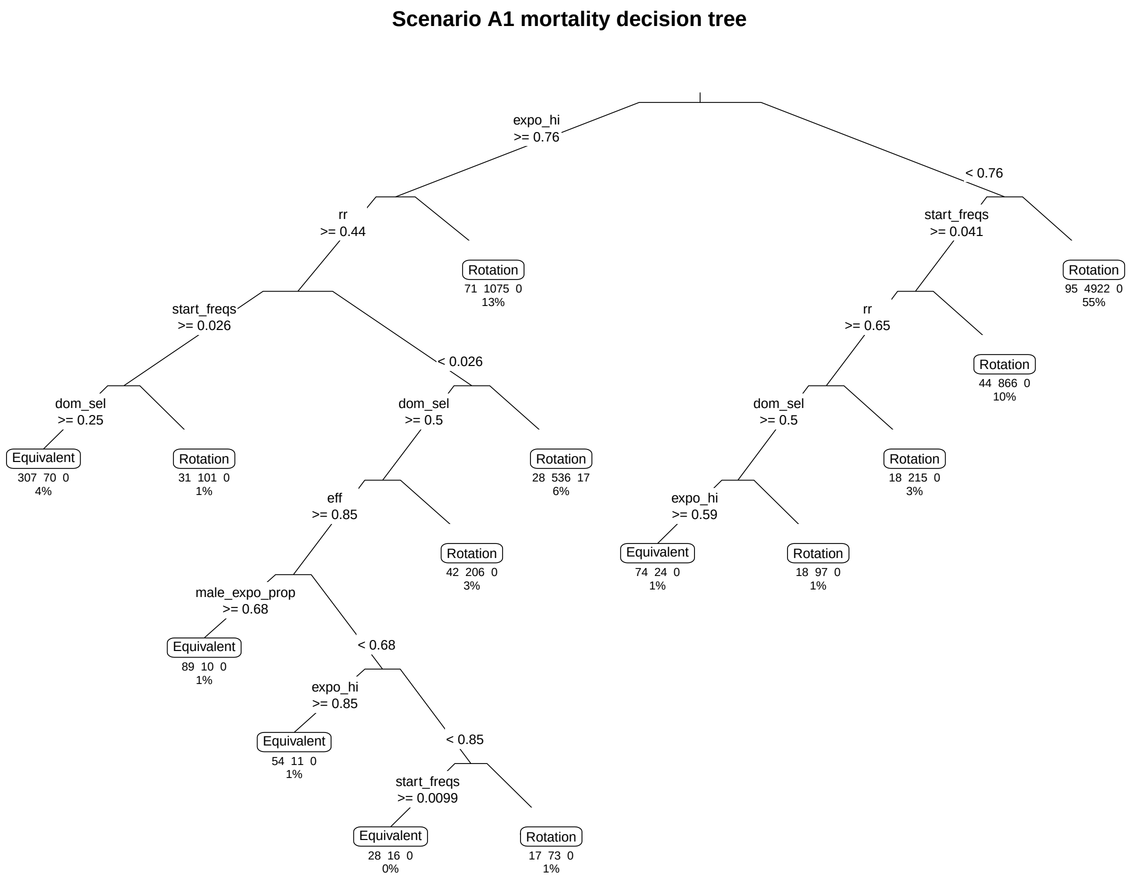

**Scenario A2 generations decision tree**

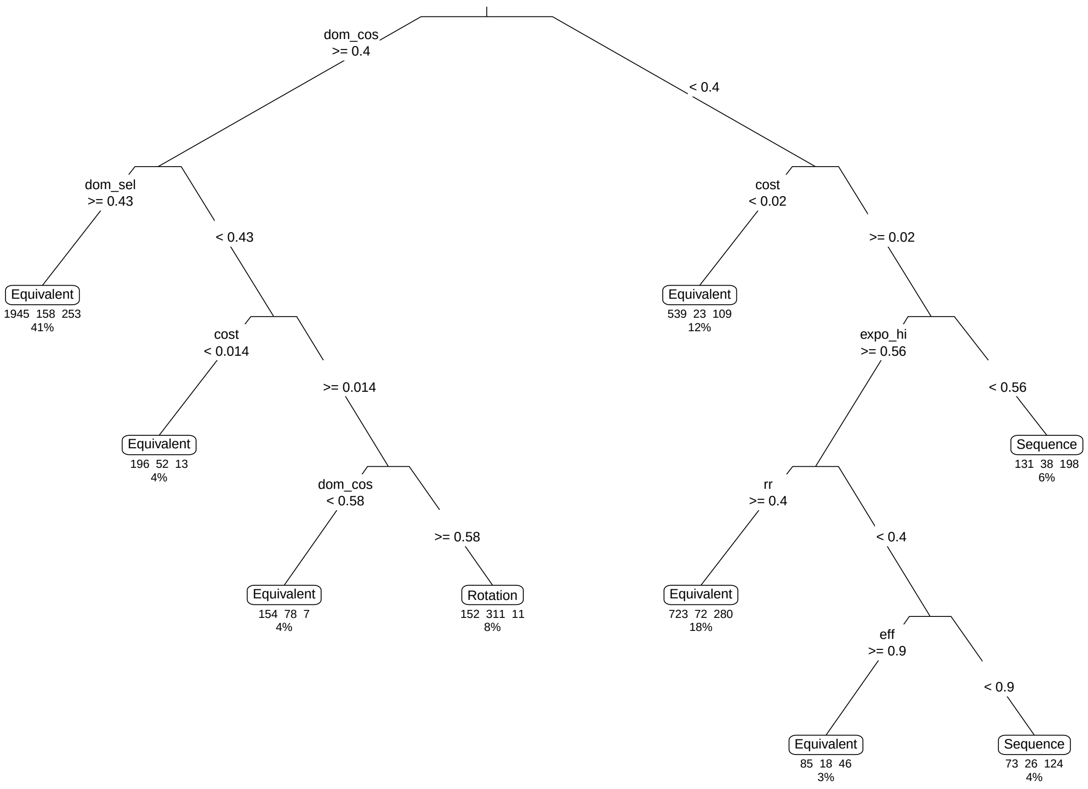

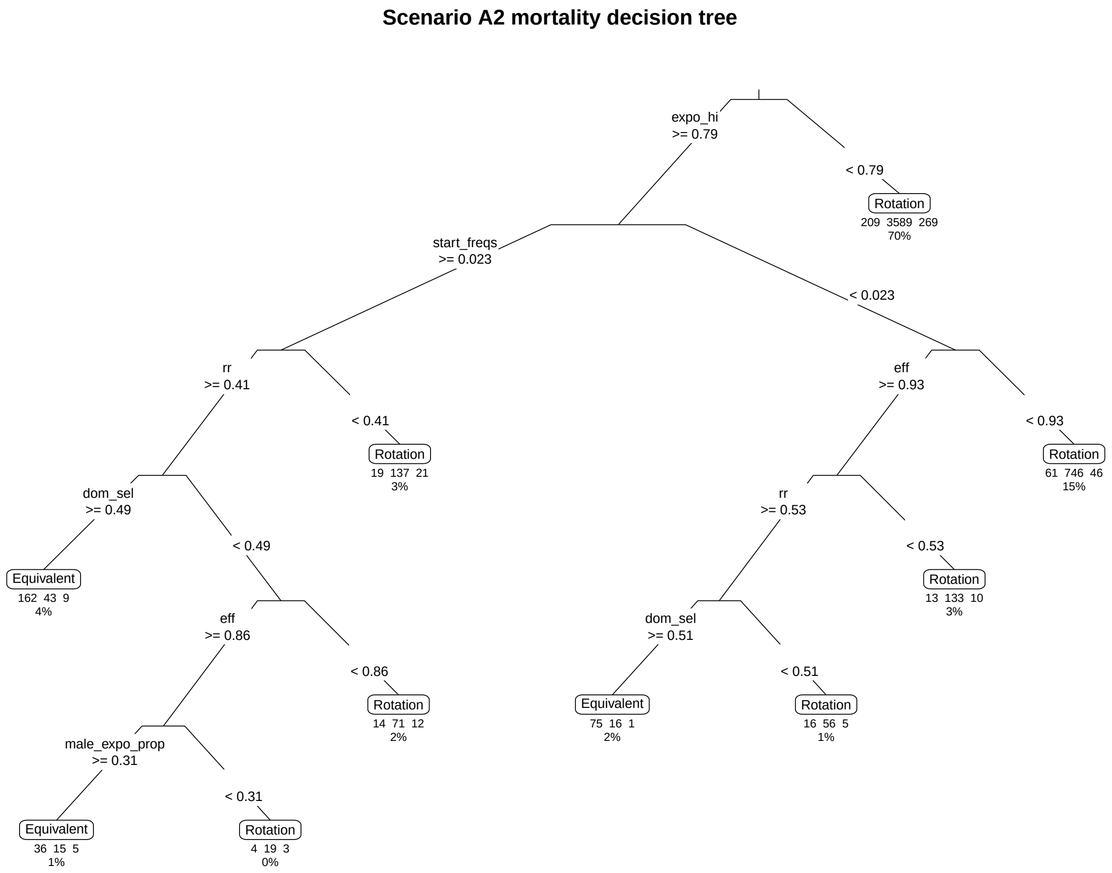
Scenario A2 mortality decision tree
expo_hi
>= 0.79
< 0.79
Rotation
209 3589 269
70%
start_freqs
>= 0.023
< 0.023
rr
>= 0.41
< 0.41
Rotation
19 137 21
3%
eff
>= 0.93
< 0.93
Rotation
61 746 46
15%
dom_sel
>= 0.49
< 0.49
Equivalent
162 43 9
4%
rr
>= 0.53
< 0.53
Rotation
13 133 10
3%
eff
>= 0.86
< 0.86
Rotation
14 71 12
2%
dom_sel
>= 0.51
< 0.51
Equivalent
75 16 1
2%
Rotation
16 56 5
1%
male_expo_prop
>= 0.31
< 0.31
Equivalent
36 15 5
1%
Rotation
4 19 3
0%

### Scenario A3 generations decision tree

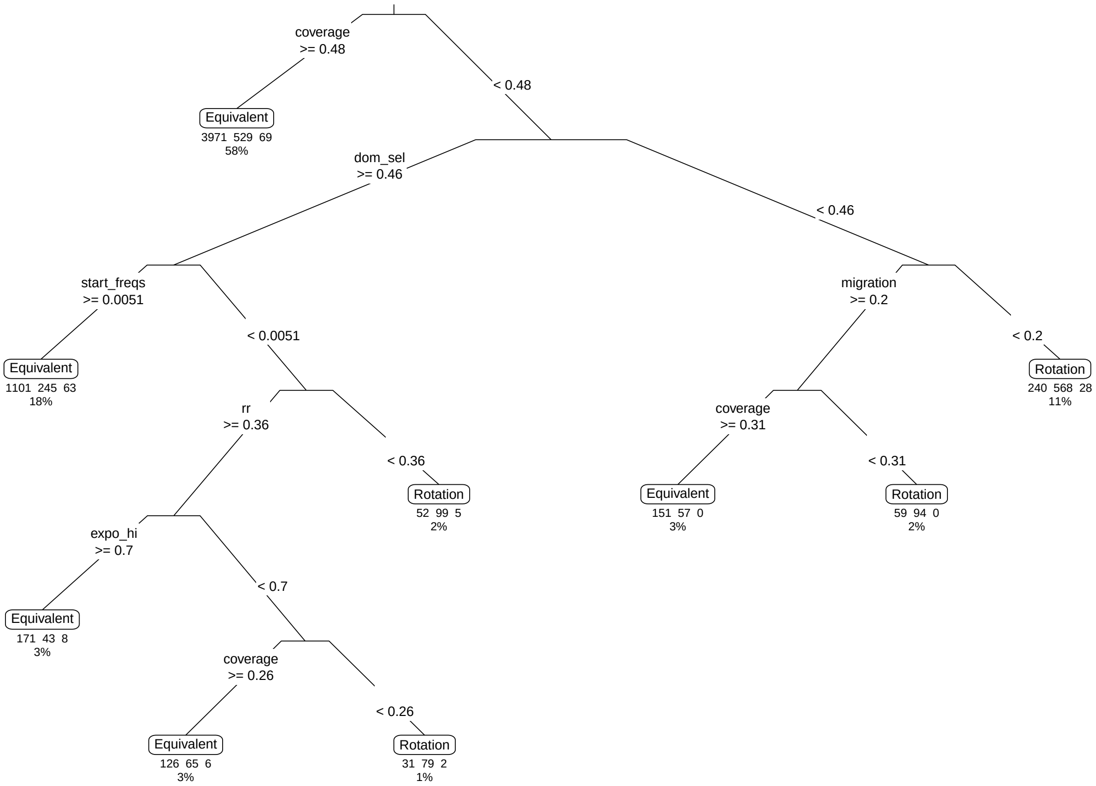

**Scenario A3 mortality decision tree**

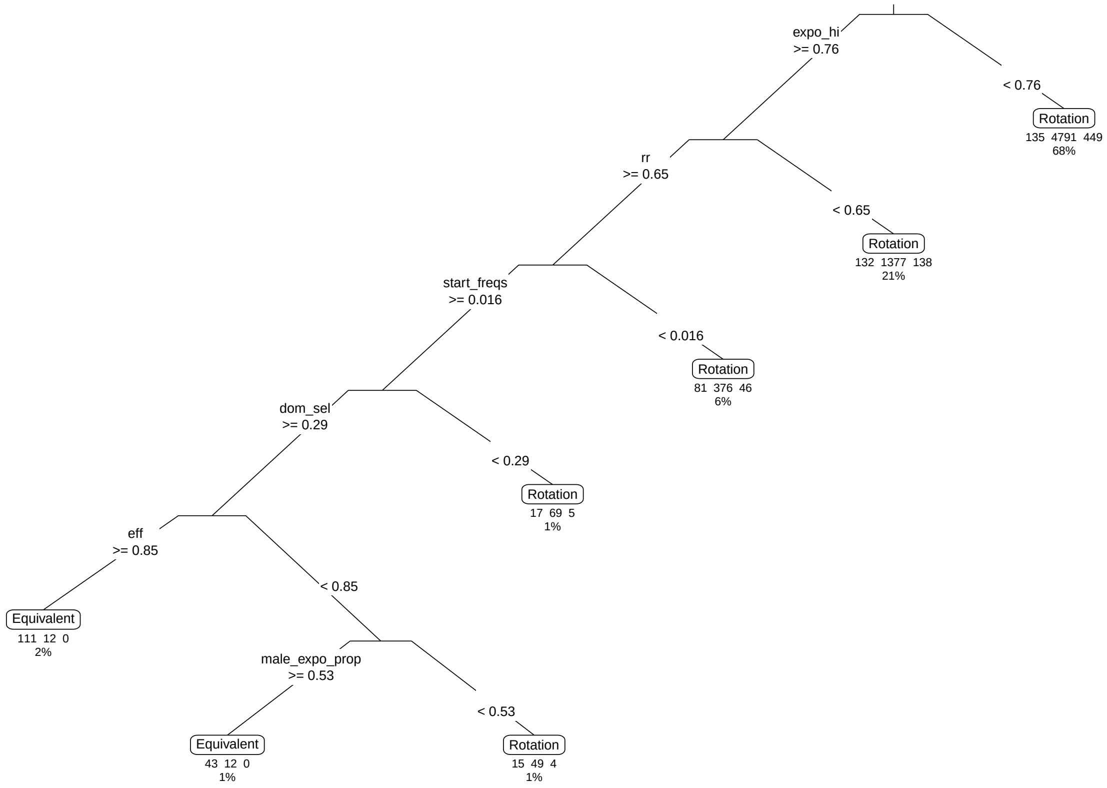

**Scenario A4 generations decision tree**

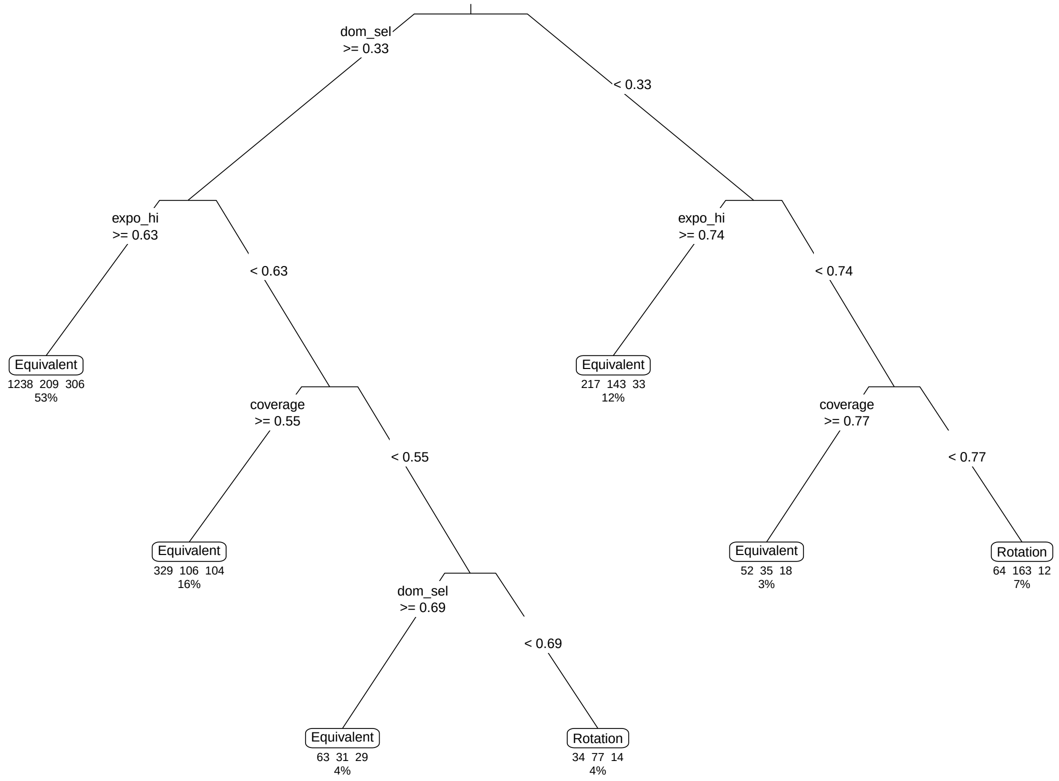

**Scenario A4 mortality decision tree**

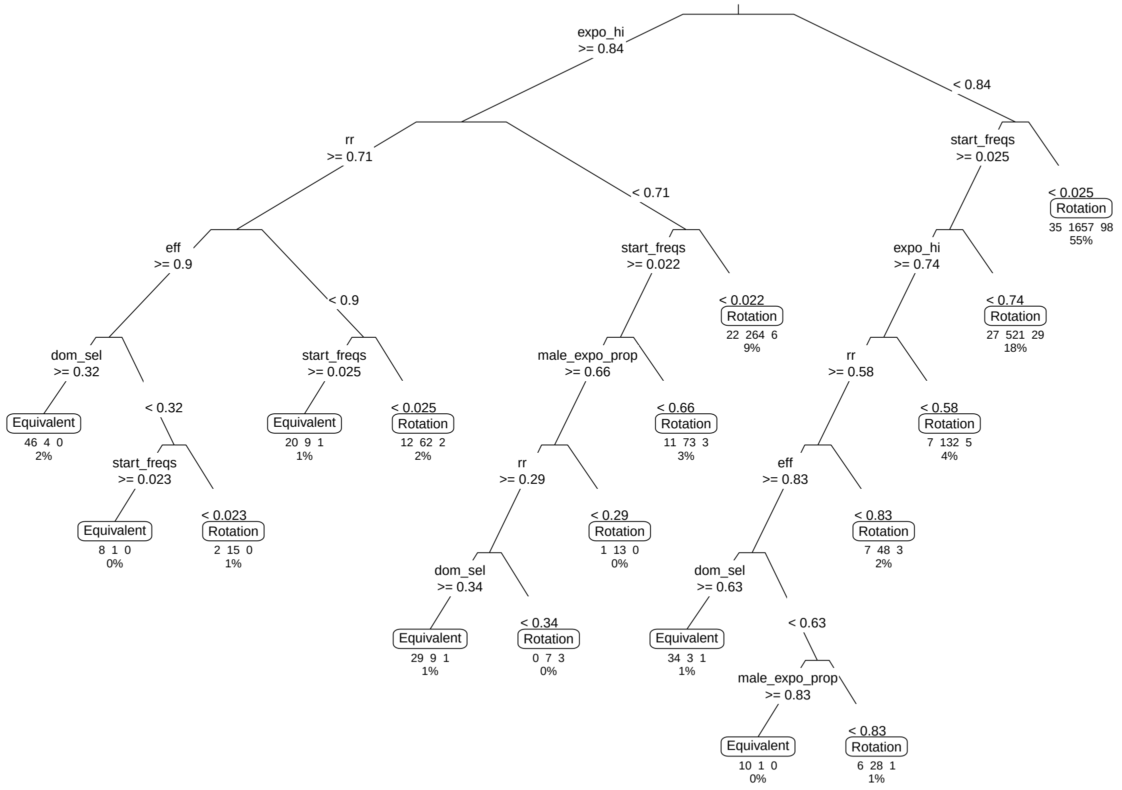

**Scenario B1 generations decision tree**

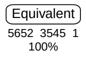

### Scenario B1 mortality decision tree

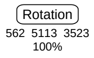

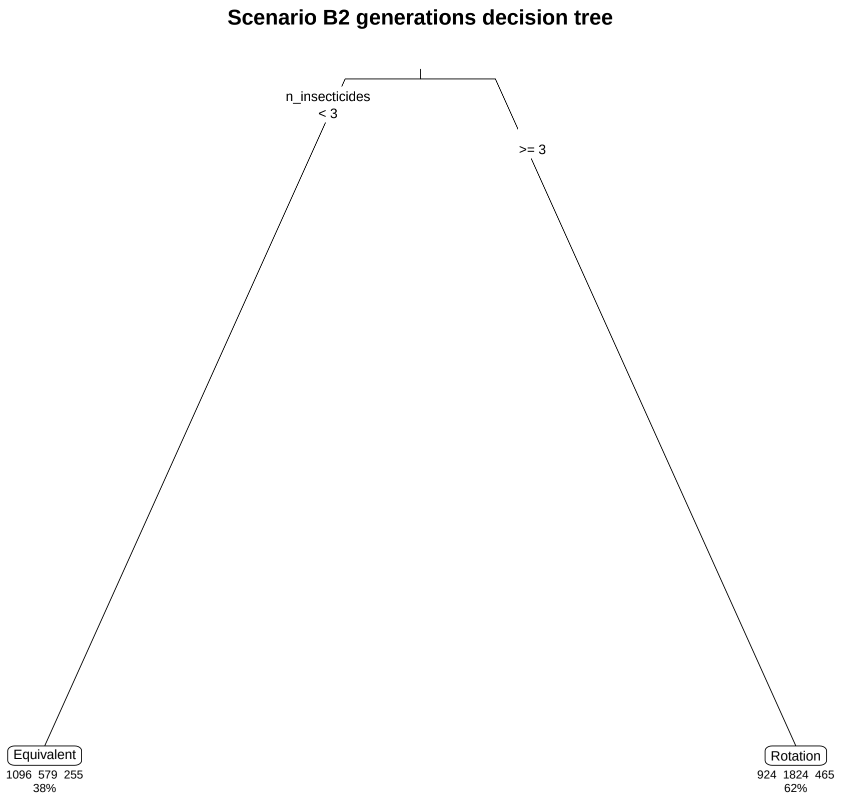
Scenario B2 generations decision tree
n_insecticides
< 3
>= 3
Equivalent
1096 579 255
38%
Rotation
924 1824 465
62%

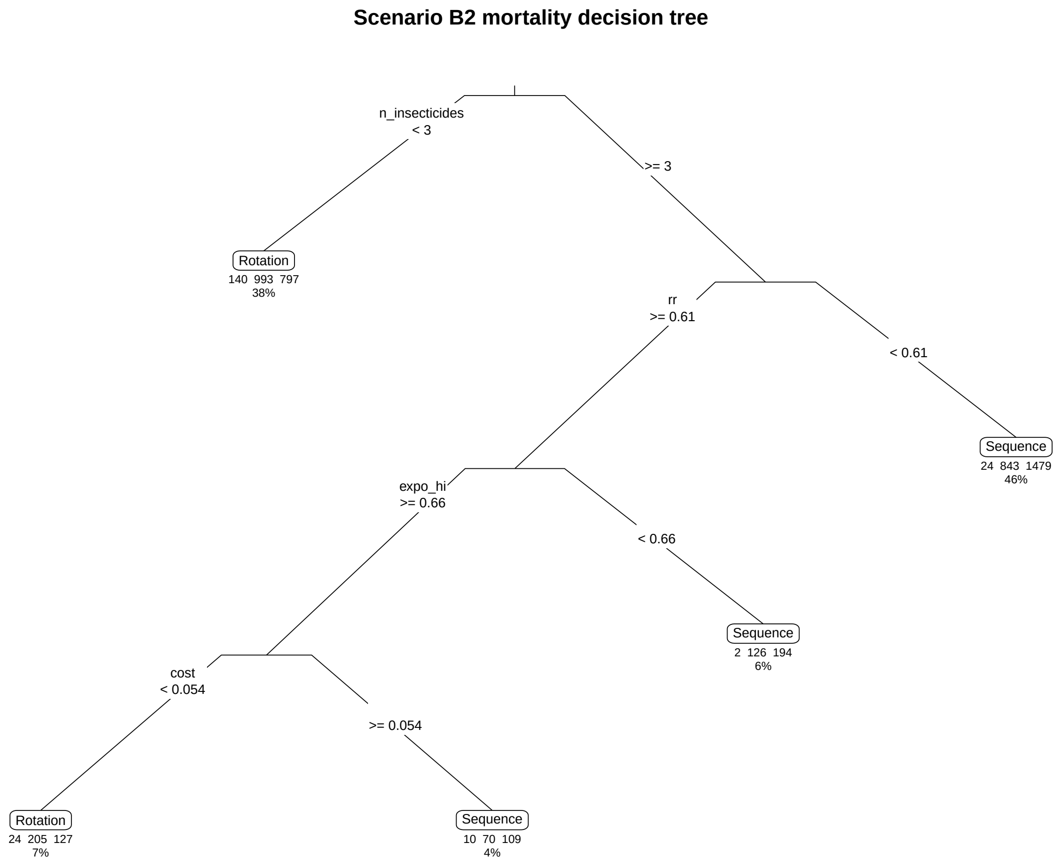
Scenario B2 mortality decision tree
n_insecticides
< 3
>= 3
Rotation
140 993 797
38%
rr
>= 0.61
< 0.61
Sequence
24 843 1479
46%
expo_hi
>= 0.66
< 0.66
Sequence
2 126 194
6%
cost
< 0.054
>= 0.054
Rotation
24 205 127
7%
Sequence
10 70 109
4%

**Scenario B3 generations decision tree**

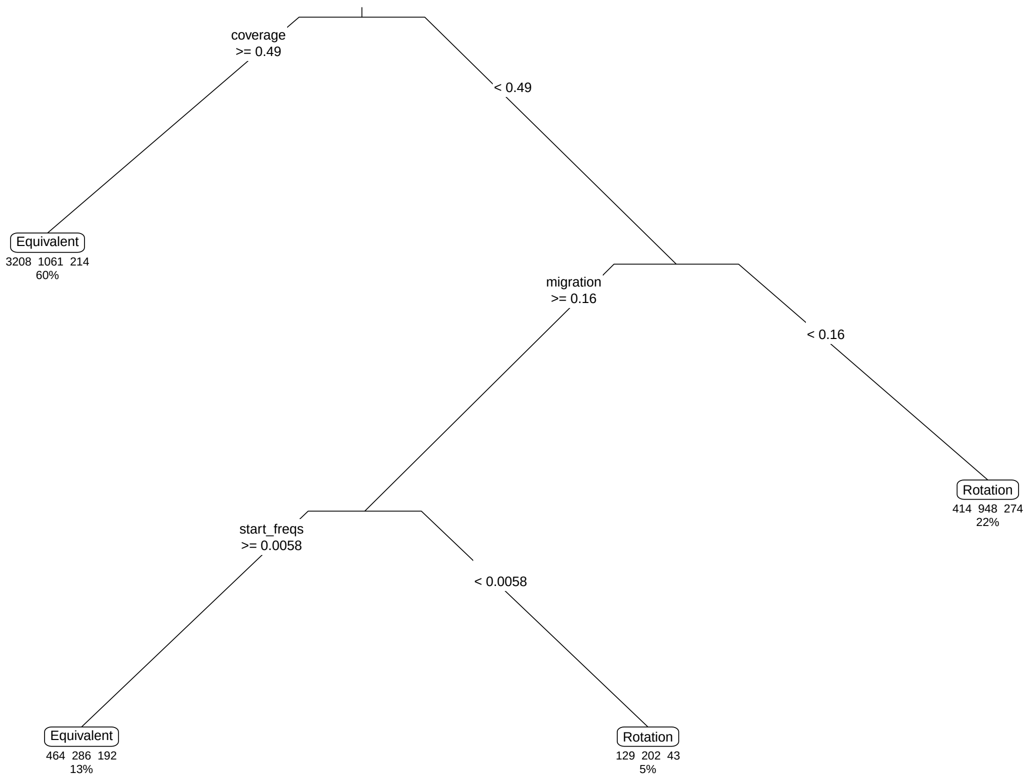

### Scenario B3 mortality decision tree

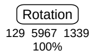

### Scenario B4 generations decision tree

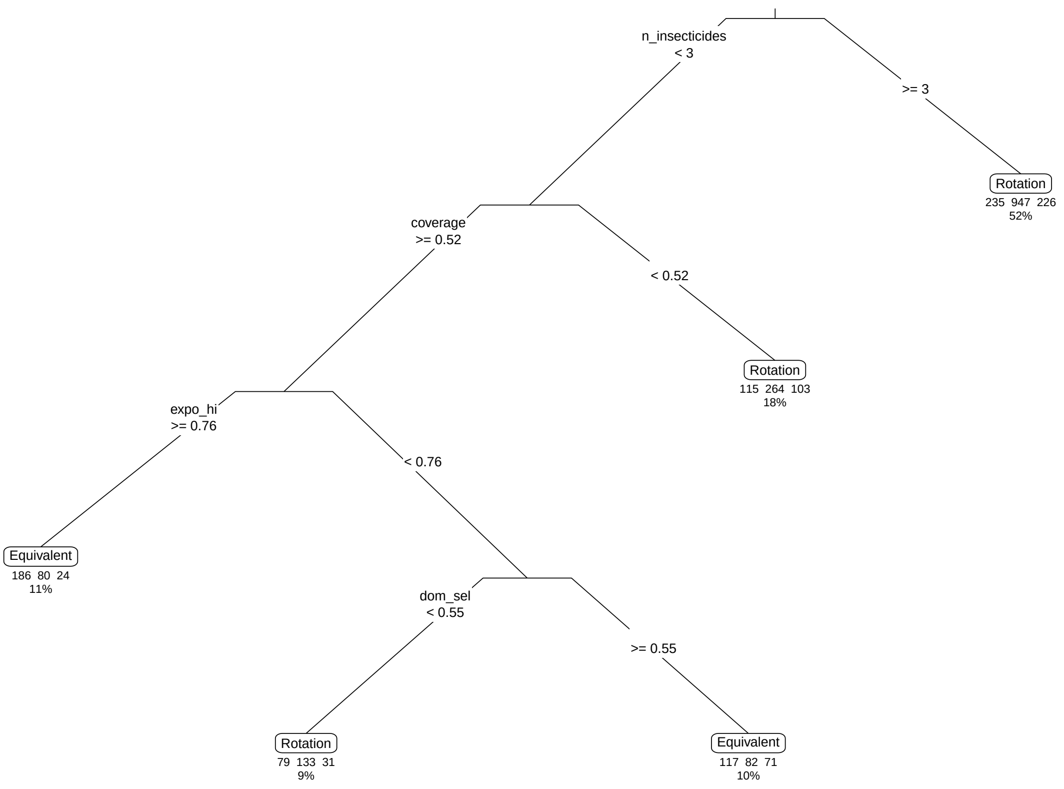

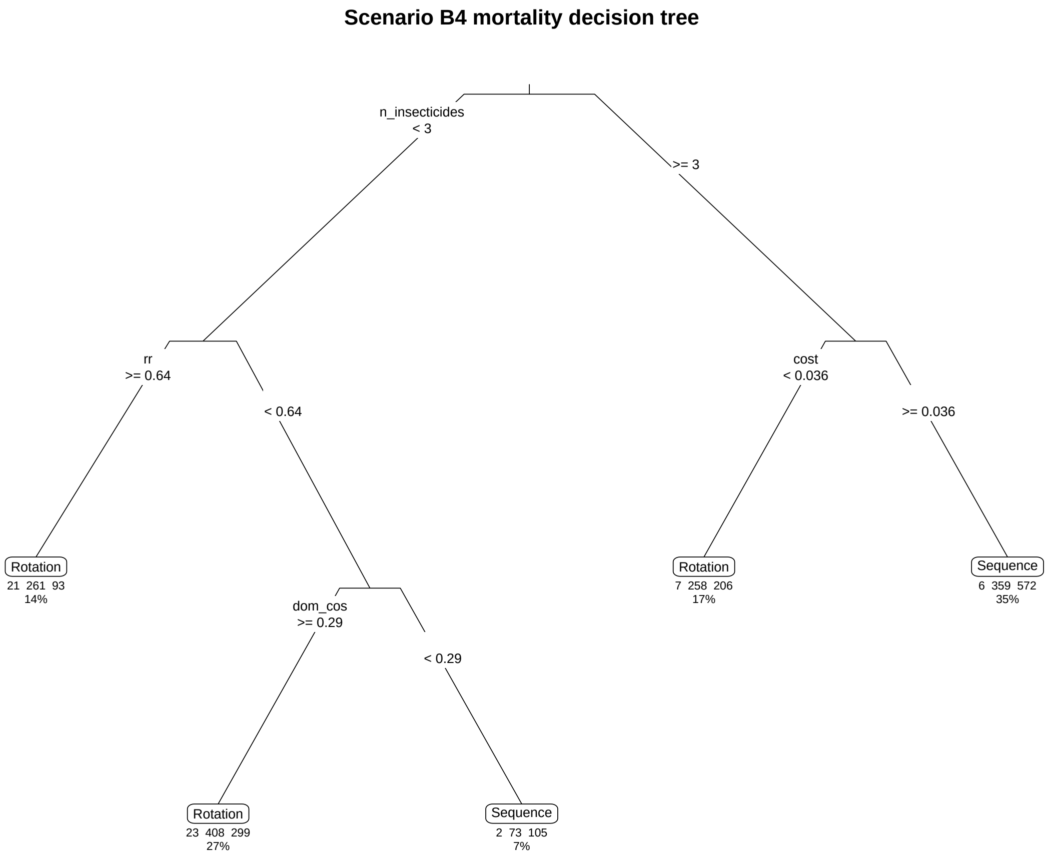


```
[1] "Scenario C1 Generations tree cannot run"
```

**Scenario C1 mortality decision tree**

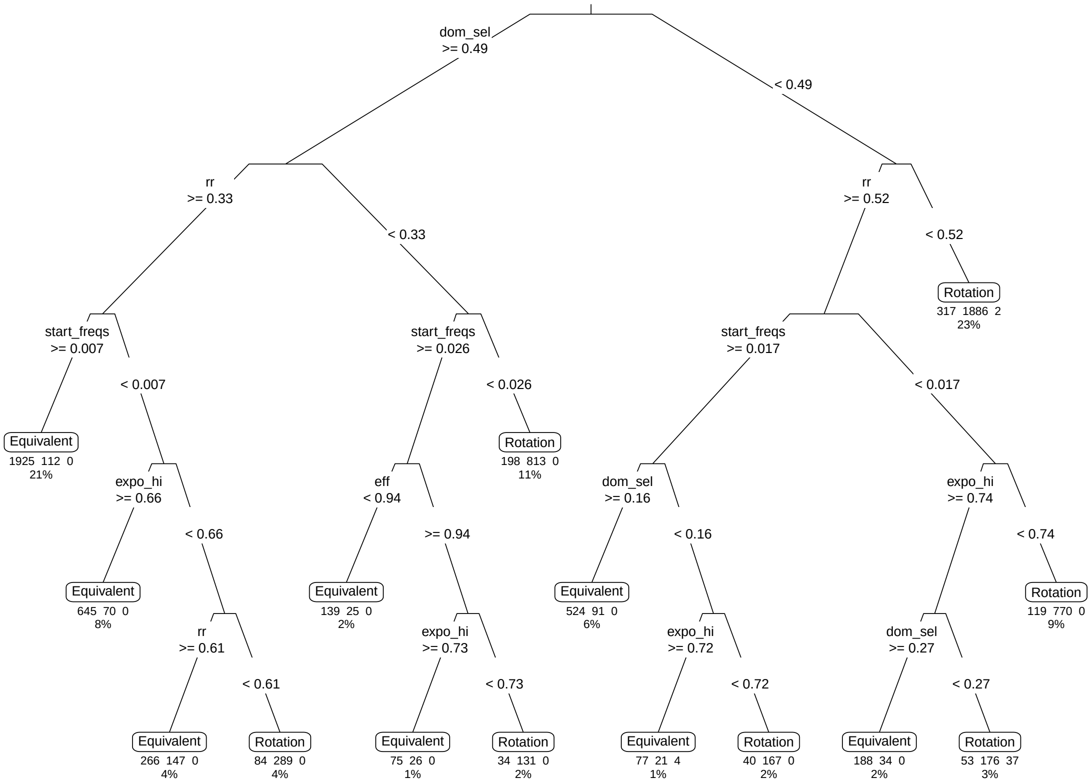

### Scenario C2 generations decision tree

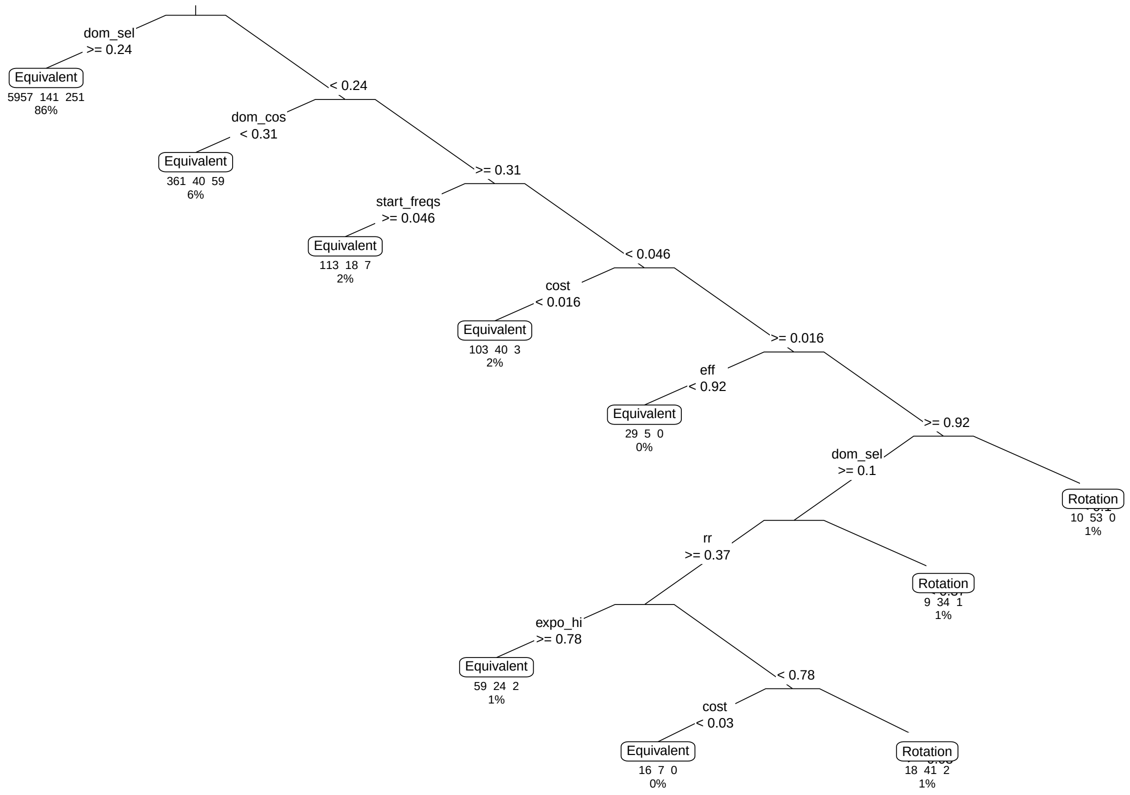

### Scenario C2 mortality decision tree

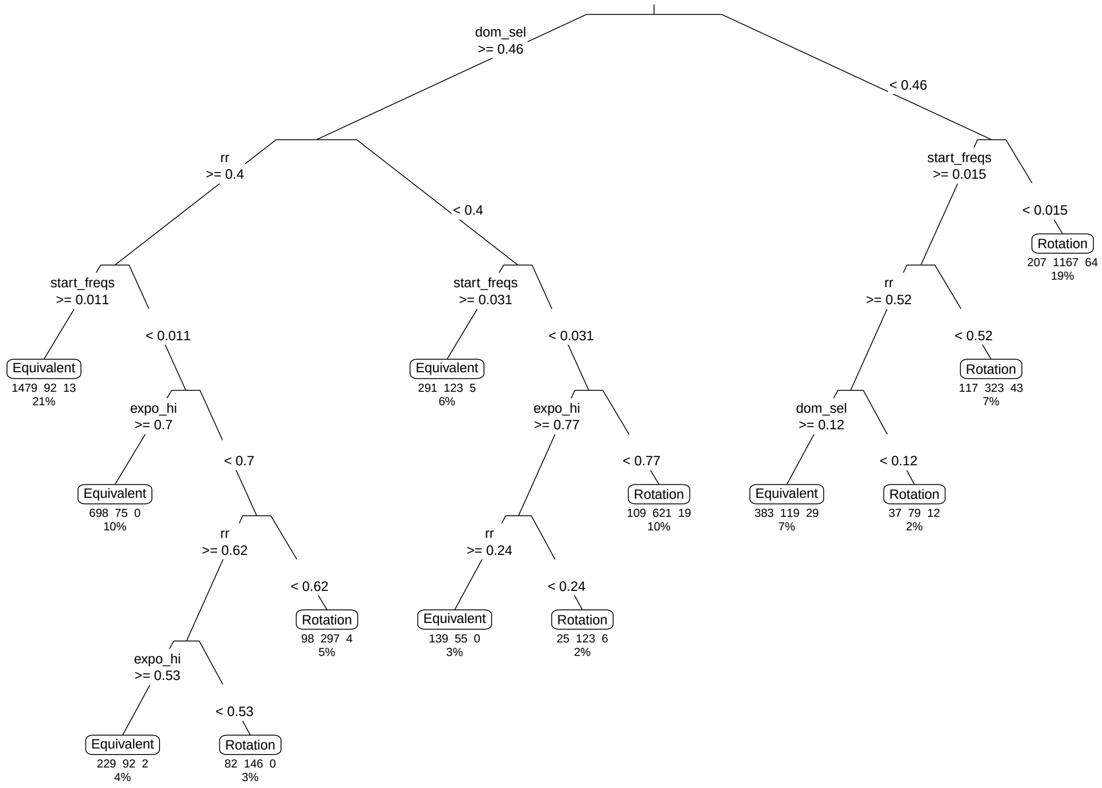

### Scenario C3 generations decision tree

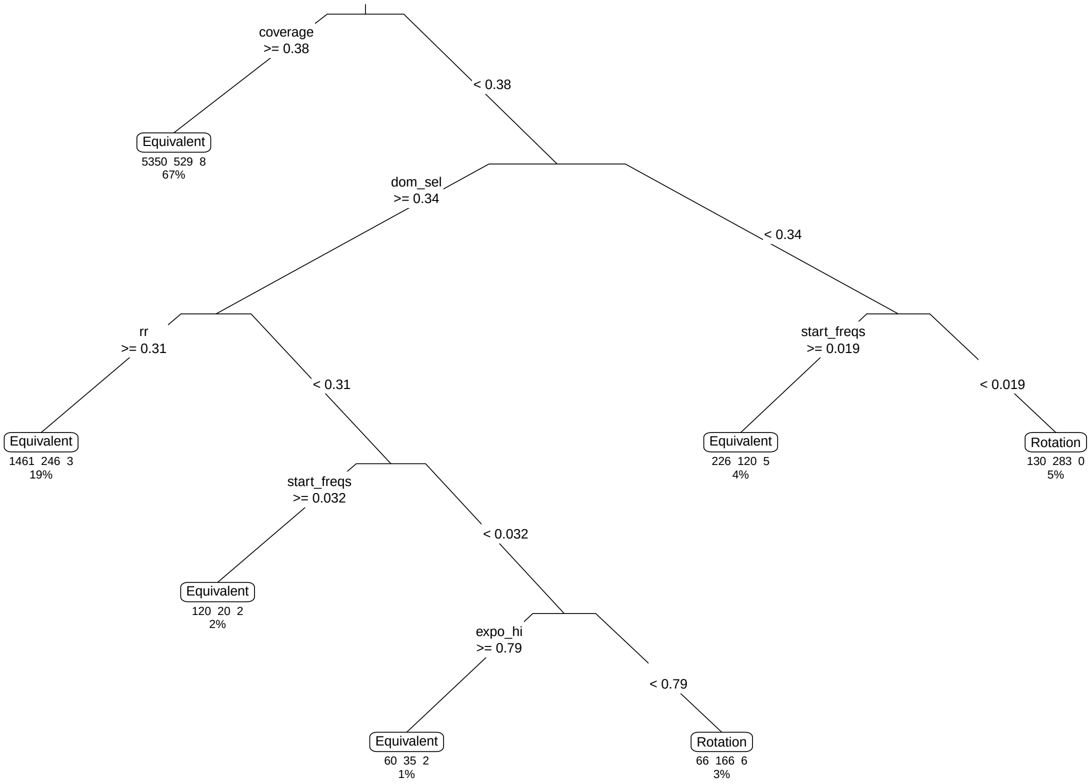

### Scenario C3 mortality decision tree

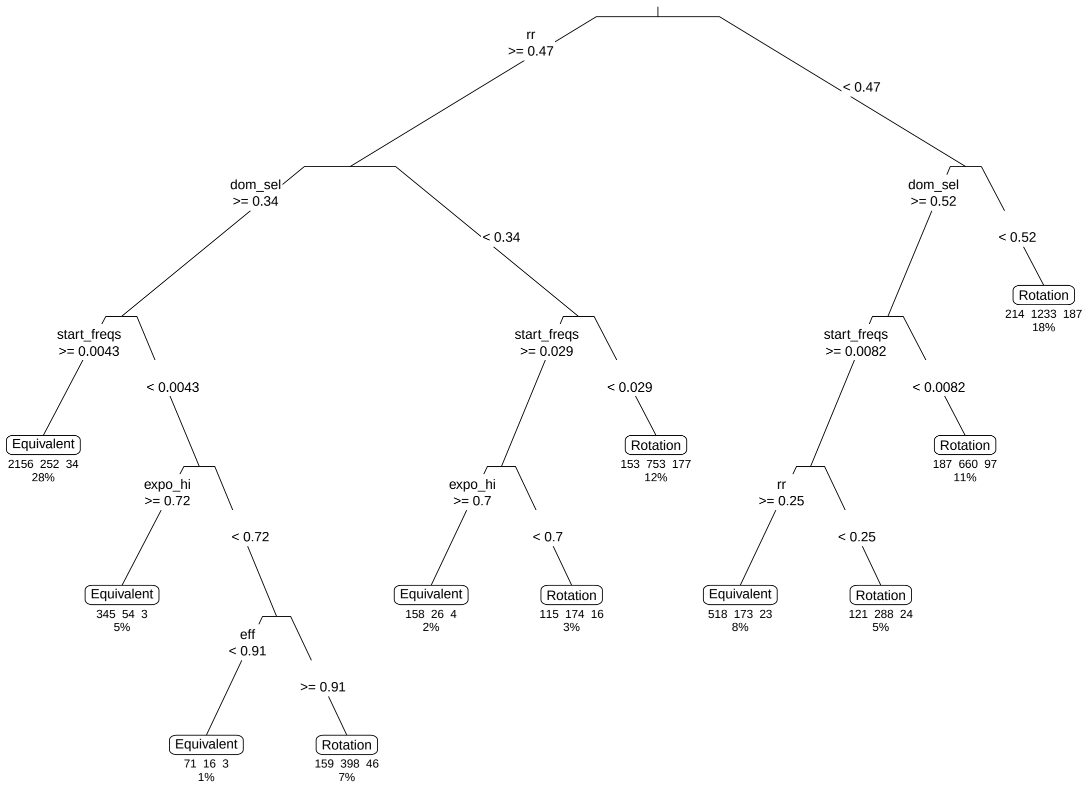

**Scenario C4 generations decision tree**

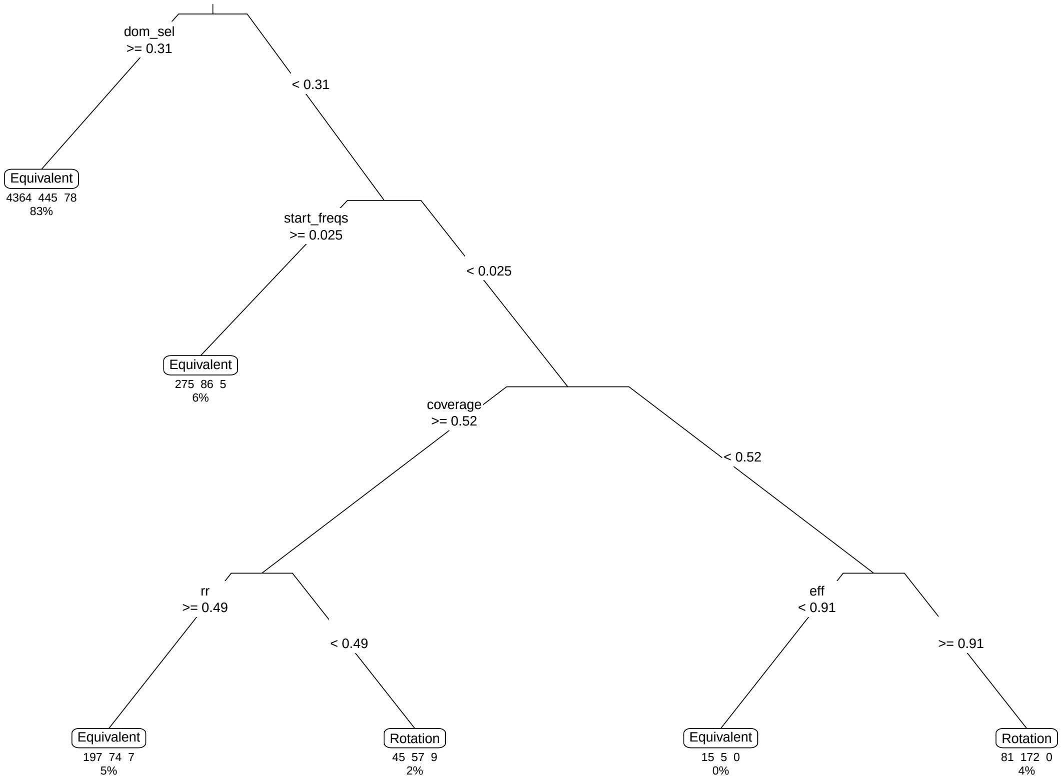

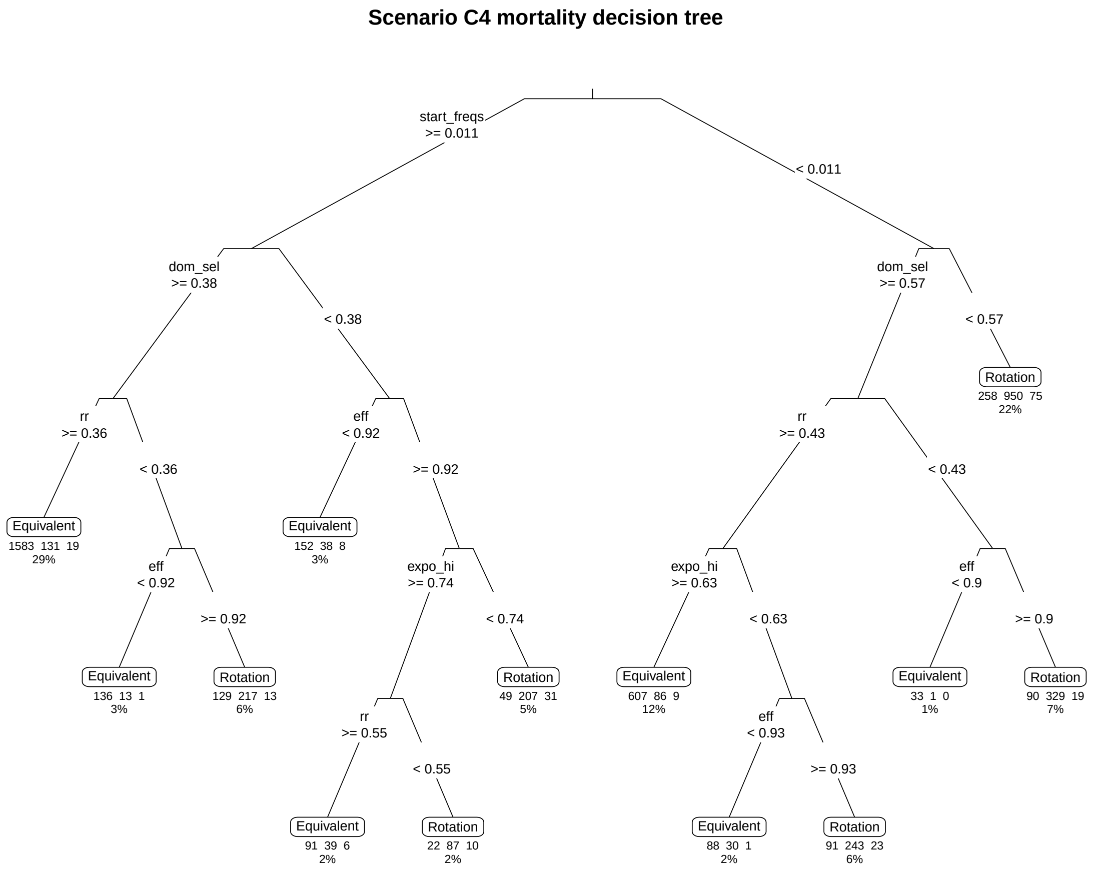


```
[1] "Scenario D1 Generations tree cannot run"
```

### Scenario D1 mortality decision tree

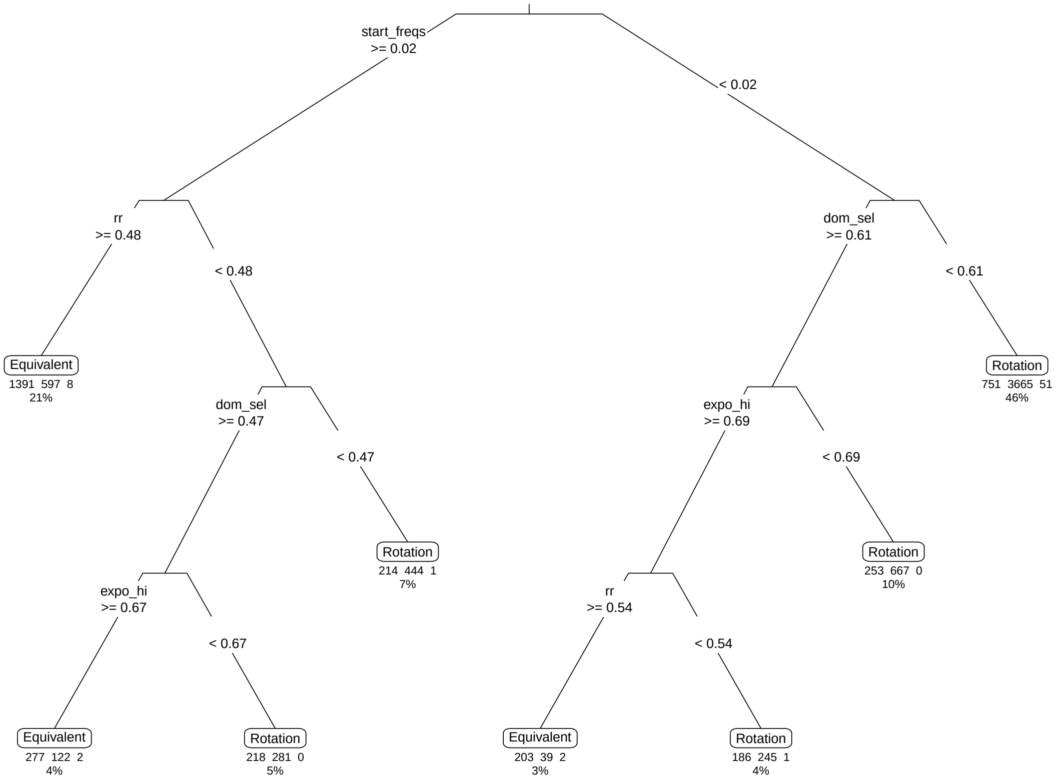

**Scenario D2 generations decision tree**

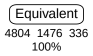

**Scenario D2 mortality decision tree**

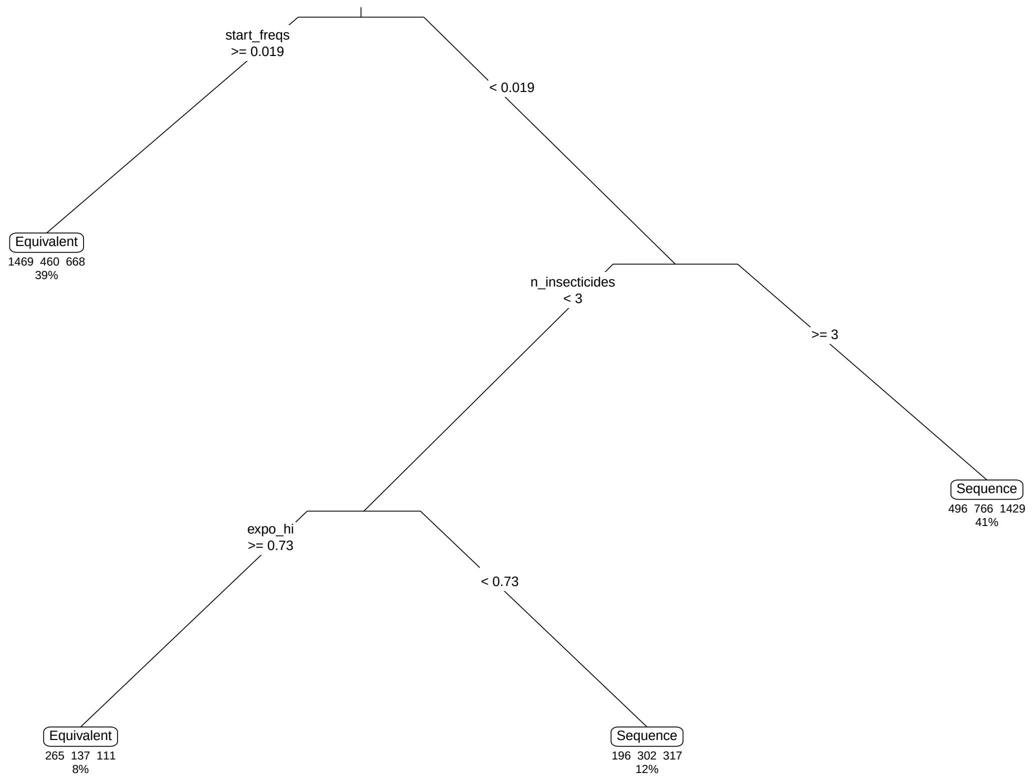

### Scenario D3 generations decision tree

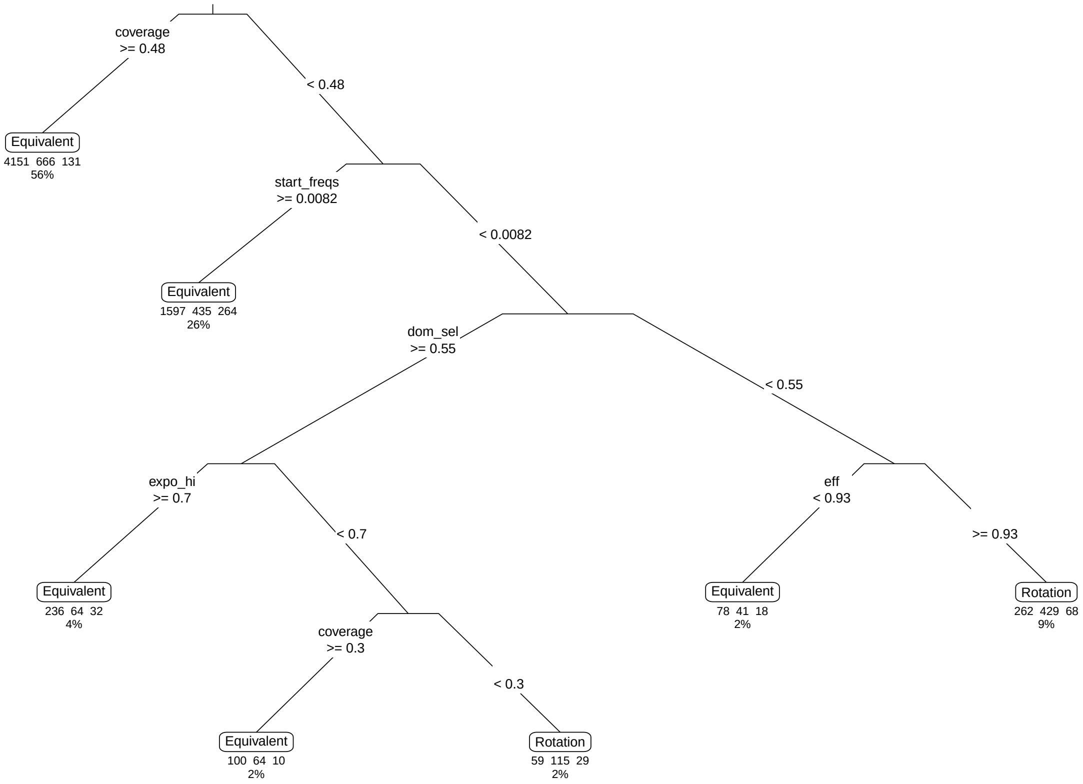

### Scenario D3 mortality decision tree

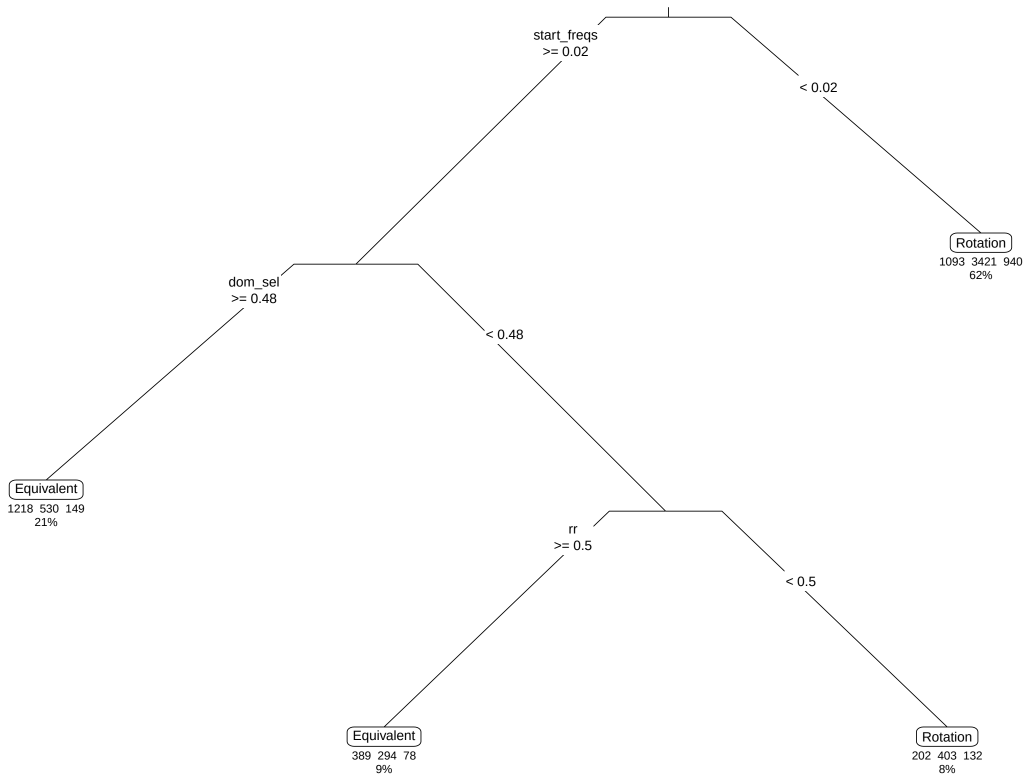

### Scenario D4 generations decision tree

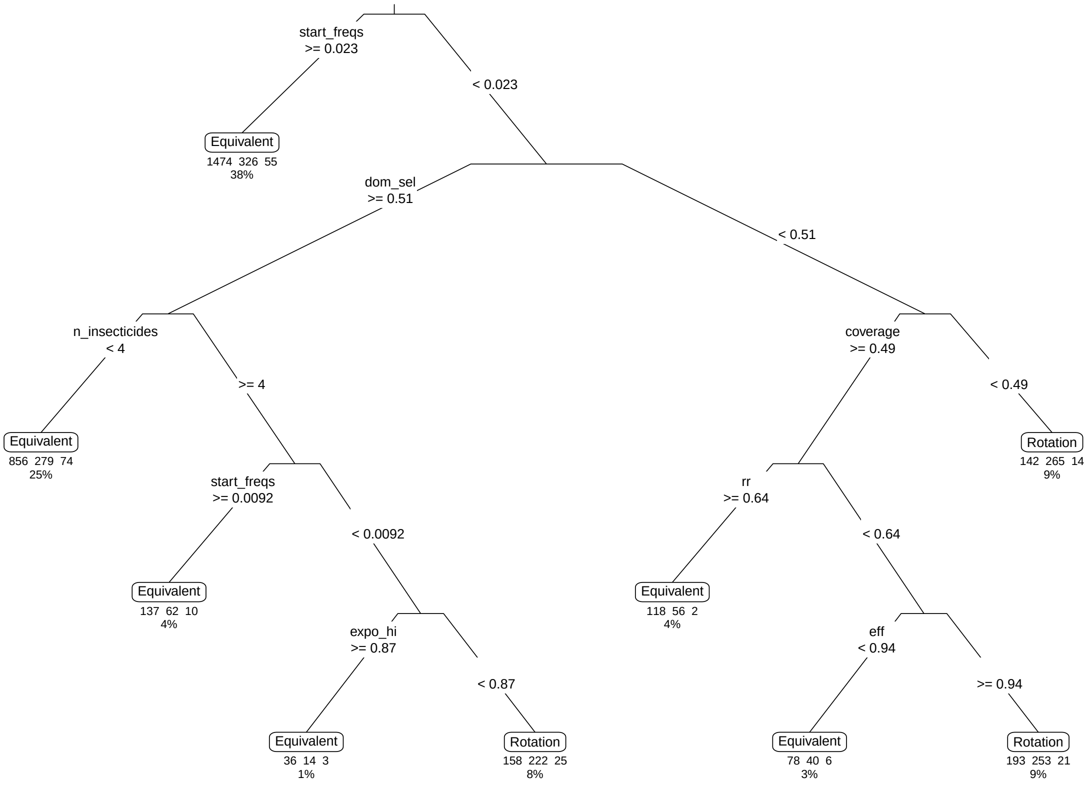

**Scenario D4 mortality decision tree**

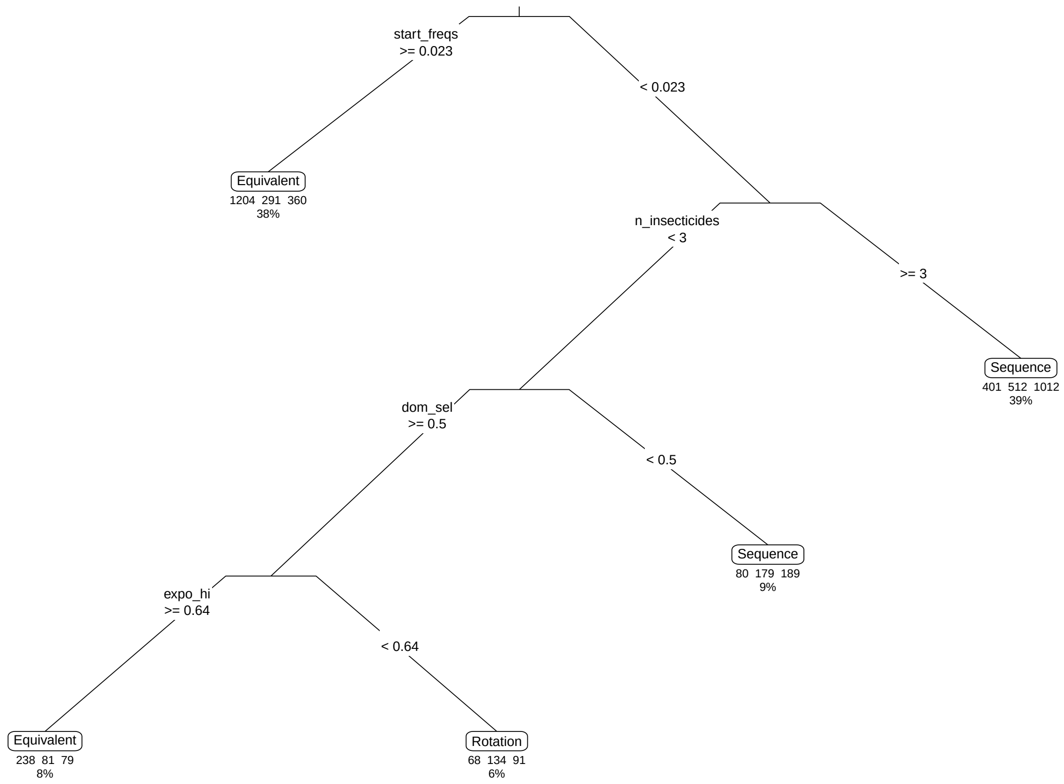
